# Commonly used Bayesian diversification-rate models produce biologically meaningful differences on empirical phylogenies

**DOI:** 10.1101/2023.05.17.541228

**Authors:** Jesús Martínez-Gómez, Michael J. Song, Carrie M. Tribble, Bjørn T. Kopperud, William A. Freyman, Sebastian Höhna, Chelsea D. Specht, Carl J. Rothfels

## Abstract

Identifying along which lineages shifts in diversification rates occur is a central goal of comparative phylogenetics; these shifts may coincide with key evolutionary events such as the development of novel morphological characters, the acquisition of adaptive traits, polyploidization or other structural genomic changes, or dispersal to a new habitat and subsequent increase in environmental niche space. However, while multiple methods now exist to estimate diversification rates and identify shifts using phylogenetic topologies, the appropriate use and accuracy of these methods is hotly debated. Here we test whether five Bayesian methods—Bayesian Analysis of Macroevolutionary Mixtures (`BAMM`), two implementations of the Lineage-Specific Birth-Death-Shift model (`LSBDS` and `PESTO`), the approximate Multi-Type Birth-Death model (`MTBD`; implemented in `BEAST2`), and the cladogenetic diversification rate shift model (`CLaDS2`)—produce comparable results. We apply each of these methods to a set of 65 empirical time-calibrated phylogenies and compare inferences of speciation rate, extinction rate, and net diversification rate. We find that the five methods often infer different speciation, extinction, and net-diversification rates. Consequently, these different estimates may lead to different interpretations of the macroevolutionary dynamics. The different estimates can be attributed to fundamental differences among the compared models. Therefore, the inference of shifts in diversification rates is strongly method-dependent. We advise biologists to apply multiple methods to test the robustness of the conclusions or to carefully select the method based on the validity of the underlying model assumptions to their particular empirical system.

**Lay Summary:** Understanding why some groups of organisms have more species than others is key to understanding the origin of biodiversity. Theory and empirical evidence suggest that multiple distinct historical events—such as the evolution of particular morphological features (e.g., the flower, the tetrapod limb) and competition amongst species—can produce this pattern of divergent species richness. Identifying when and where on the tree of life shifts in diversification rates occur is important for explaining the origin of modern-day biodiversity and understanding how disparity among species evolves. Several statistical methods have been developed to infer diversification rates and identify these shifts. While these methods each attempt to make inferences about changes in the tempo of diversification, they differ in their underlying statistical models and assumptions. Here we test if these methods draw similar conclusions using a dataset of 65 time-calibrated phylogenies from across multicellular life. We find that inferences of where rate shifts occur strongly depends on the chosen method. Therefore, biologists should choose the model whose assumptions they believe to be the most valid and justify their model choice *a priori*, or consider using several independent methods to test an evolutionary hypothesis.

## Introduction

Understanding the patterns and processes that shape the tree of life is one of the central pursuits of biology. However, inferring the tempo of evolution among lineages— the patterns of speciation and extinction that gave rise to our extant biodiversity—remains a difficult problem both theoretically and computationally (Rabosky, 2010; Moore et al., 2016; Louca and Pennell, 2020).

Several methods estimate diversification rates (speciation and extinction rates, individually) assuming that rates are constant across the tree (Morlon, 2014). Recently developed methods have built upon constant-rate models by allowing diversification parameters to vary depending on the state of a focal character (Maddison et al., 2007) or, even more recently, among branches of the phylogeny, which allows for lineage-specific diversification rate estimates (*e.g*., Rabosky, 2014; Hö hna et al., 2019; Barido-Sottani et al., 2020; Maliet and Morlon, 2022).

Such lineage-specific methods have the potential to offer powerful insights into our understanding of evolution, such as the potential time-dependency of macroevolutionary diversification (Henao Diaz et al., 2019), the macroecological and macroevolutionary causes of the latitudinal diversity gradient (Givnish et al., 2018; Rabosky et al., 2018), and macroevolutionary support of Darwinian and Simpsonian theories of microevolution within adaptive zones (Cooney et al., 2017).

The application of these methods, however, has been marred by controversy over their implementation (Moore et al., 2016; Rabosky et al., 2017; Meyer and Wiens, 2018; Meyer et al., 2018; Rabosky, 2018) and by theoretical findings that seemingly undermine the general reliability of inferring diversification parameters from phylogenies of extant species (Louca and Pennell, 2020; Helmstetter et al., 2021). These issues are liable to discourage empiricists, who may wonder if the disagreements among model developers and theorists correspond with biologically relevant inference differences in empirical studies.

To address this question, we assess how inferences under five leading contemporary Bayesian methods— Bayesian Analysis of Macroevolutionary Mixtures (`BAMM;` Rabosky, 2014); the Lineage-Specific Birth-Death-Shift model (LSBDS; Hö hna et al., 2019) and its MCMC-free implementation: Phylogenetic Estimation of Shifts in the Tempo of Origination (`PESTO`; Kopperud et al., 2023a); the approximate Multi-Type Birth-Death model (`MTBD`; Barido-Sottani et al., 2020); and the Cladogenetic Diversification Rate Shift model (CLaDS2; Maliet et al., 2019)— compare to each other.

While all five methods aim to estimate lineage-specific diversification rates, they differ in how and where rate shifts are allowed to occur.

1. `BAMM` models diversification rates as varying across lineages by testing among models that include different numbers of diversification-rate regimes (sets of speciation and extinction parameters) and different placements of those regimes in the tree; however `BAMM` does not model rate shifts on extinct (thus unobserved) branches (Rabosky, 2014).
2. The `LSBDS` model, as implemented in RevBayes (Hö hna et al., 2016), samples rate regimes from a prior distribution discretized into a fixed number of rate categories; this discretization facilitates computation and allows the method to model shifts on extinct branches (Hö hna et al., 2019).
3. `PESTO` is a new implementation of the `LSBDS` model that analytically computes the posterior mean speciation and extinction rates conditional on a set of hyperparameters without the need for Monte Carlo sampling (Kopperud et al., 2023a).
4. The `MTBD` method is based on a multitype birth-death process that infers the number of rate regimes as well as the transition rate *γ* between rate regimes (Barido-Sottani et al., 2020). This approach allows for the same rate regime to be present in different parts of the tree. The approximate `MTBD`, tested here, assumes that no rate changes occur in the extinct parts of the tree; this approximation, when applied with a high transition rate prior, has been found to not substantially differ from the exact `MTBD` method, which allows rates changes along extinct lineages (Barido-Sottani et al., 2020).
5. Finally, in the `CLaDS2` model, diversification rates only change at speciation events. Descendant lineages inherit the speciation rate via a stochastic process that is influenced by the *α* parameter, which represents the long-term trend (i.e., increase or decreases) of the speciation rate (Maliet et al., 2019). This model results in many small and frequent shifts in diversification rates regimes, unlike the other methods, which tend to infer a few large shifts in rate regimes (Maliet et al., 2019; Maliet and Morlon, 2022). Another aspect of `CLaDS2` is that extinction rates are not inferred per branch. Instead, the model estimates a global turnover parameter (*e* = *μ*_*i*_/*λ*_*i*_). However, shifts are allowed to occur along extinct branches.

Other methods, not tested here, leverage hidden states using a maximum likelihood framework (*e.g*., Vasconcelos et al., 2022).

To assess whether the theoretical and computational differences among these methods result in biologically meaningful differences, we reanalyze 65 empirical datasets, compiled from Henao Diaz et al. (2019), using each of `BAMM, LSBDS, PESTO, MTBD,` and `CLaDS2`. We address the question: do different analytical methods for estimating branch-specific diversification rates produce significantly different results across an array of empirical datasets?

## Methods

### Empirical Data

Our empirical data are derived from from the set of 104 chronograms compiled and analyzed with `BAMM` by Henao Diaz et al. (2019). From the Henao Diaz et al. set we excluded trees with fewer than 30 extant taxa in order to concentrate on more informative datasets, resulting in our final set of 76 chronograms.

### Model Settings

Our goal was to apply each method as a typical diligent user might. For each chronogram, we used the incomplete-sampling fraction collected from the original study by Henao Diaz et al. (2019), and applied that sampling fraction when we ran each of the five inference methods. While the methods differ in their specific parameterizations of the birth-death process, we attempted to use comparable settings and priors across methods.

For `BAMM` analyses, we modified the control files from Henao Diaz et al. (2019). We set lambda to be time-constant rather than time-variable in order to more closely match the inferences of other methods and given concerns about the statistical validity of timevarying diversification analyses (Louca and Pennell, 2020). We set `BAMM` priors for each phylogeny using the set`BAMM`priors() function in the `BAMM`tools R package (Rabosky et al., 2014b). This function computes dataset-specific priors by estimating metrics from the dataset such as the root age of the chronogram and then estimating reasonable and broad expectations for shifts and rates. We ran `BAMM` v. 2.5.0 using the priors BAMMtools and control files, which determined the phylogeny-specific number of generations for a single MCMC chain. We removed the first 10% of the MCMC samples as burnin and assessed convergence by computing estimates of effective sample size (ESS) using the R package Coda (Plummer et al., 2006). We specifically looked for convergence of the log-likelihood parameter and the ‘number of distinct regimes’ parameter, as is recommended (Rabosky et al., 2014a). Analyses that did not reach convergence were run for additional generations until they converged.

For LSBDS analyses we used the same set of priors for all phylogenies (except for sampling fraction) with eight categories for speciation and for extinction (64 total rate categories). The number of rate categories was chosen after performing a test on one representative phylogeny, which found that increasing the rate categories above 64 did not result in a significant change in model fit. For each chronogram we ran four MCMC chains for 5,000 generations. Convergence was assessed for each chain by checking that the ESS values for all model parameters in the log files were greater than 200 using the R package Coda (Plummer et al., 2006). Chains that did not reach convergence were restarted and run for an additional 5000 generations. We merged the posteriors, retaining the last 4000 generations from the MCMC (10% burnin for restarts-non and 60% burnin for restarts).

We applied `PESTO` in a three-step fashion. First, we estimated the parameters of a constant-rate birth-death process and treated these as hyperparameters: the speciation rate (*λ*) and the extinction rate (*μ*). Second, we set up a state-dependent speciation-extinction (SSE) model. In this model, we used rate values that correspond to ten quantiles of two lognormal distributions with medians *λ* and *μ*, and standard deviation 0.587. In the SSE model, we used all pairwise comparisons of these (i.e. 100 rate categories). Further, we estimated the shift rate parameter *η* conditional on the speciation and extinction rates, using maximum likelihood. Third, we calculated the posterior state probabilities along each branch. Finally, we plotted the posterior mean rates averaged over the time span for each individual branch.

We ran the `MTBD` model under default priors (implemented in `BEAST2;` Bouckaert et al., 2014; Barido-Sottani et al., 2020). We ran three MCMC chains for 100,000,000 generations per phylogeny. We removed the first 25% as burnin and assessed MCMC convergence by checking that ESS values were higher than 200 for all rates.

We ran `CLaDS2` using the default priors (as described in Maliet and Morlon, 2022). We ran three MCMC chains for each dataset and took a 25% burnin, as is the default setting for CLaDS2. Convergence was assessed by calculating the Gelman statistic (Gelman et al., 2014) every 1000^th^ generation and stopping the analysis once it achieved a Gelman statistic of 1.05, following the standard guidelines for CLaDS2.

### Convergence analysis

In cases where MCMC convergence was difficult, we aimed to determine the potential underlying cause. To assess whether the subset of trees where one or more method failed to converge was substantially different from the subset that did converge, we compared descriptive metrics including phylogeny size, phylogeny age, incomplete sampling fraction, branch length variance, and multidimensional scaling (MDS) via Robinson-Foulds (RF Robinson and Foulds, 1981) and Kuhner-Felsenstein (KF, Kuhner and Felsenstein, 1994) distances.

### Processing Model Output

We obtained estimates of the relevant diversification parameters (*e.g*., speciation rate, extinction rate, etc.) from each model. `BAMM` posterior estimates of speciation rate and extinction rate were extracted using the `getMarginalBranchRateMatrix()` function in the `BAMM`tools R package (Rabosky et al., 2014b).

We extracted LSBDS posterior distributions from the stochastic branch rate log file produced by the `mnStochasticBranchRate`() function in `LSBDS`. In the `PESTO` analyses, we computed the branch rates averaged across the branch. If *λ*_*k*_ and *μ*_*k*_ are the rate values in state *k*, and *P*_*k*_(*t*) is the posterior probability of being in state *k* at time *t*, then the average net-diversification rate along a branch is

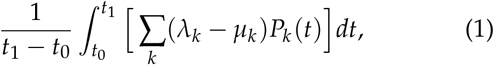

where *t*_0_ is the youngest and *t*_1_ is the oldest end point of the branch.

The posterior distributions of speciation and extinction rates of the MTBD model were obtained from the extended Newick file produced by BEAST2 using a modified read.beast() function from the treeio package (Wang et al., 2020). As CLaDS2 does not directly infer extinction rates, we calculated extinction rates per branch by multiplying the inferred global turnover value (*e*) by the branch-specific speciation rates (*μ*_*i*_ = *λ*_*i*_ *∗ e*). For all branches and models, we calculated net diversification by subtracting extinction rate from speciation rate (*λ*_*i*_ *−μ*_*i*_) per MCMC generation.

### Comparing Model Inferences

To compare inferences among the five models, we (1) visualized rate estimates on individual chronograms, (2) summarized inferences across all chronograms in the dataset to reveal systematic differences, (3) identified differences in the location and magnitude of inferred shifts among methods, and 4) tested for overlap in the 95% HPD interval of the posterior distributions.

#### Visualizing rates on trees

The canonical way of presenting the results of branch-specific diversification-rate analyses is by coloring the branches of the tree by the estimated rates. For each tree, we colored each branch by the posterior median estimate of speciation, extinction, and net diversification to visualize if the methods inferred similar shifts in similar locations on the tree.

#### Comparing rate estimates by method

To understand whether the methods displayed any consistent differences across the chronograms, we calculated six summary statistics for each tree. For each diversification rate (*i*.*e*., speciation rate, extinction rate, and net diversification rate) we calculated the posterior medians for each branch, and from those posterior medians we calculated the tree-wide mean and variance in branch rates for each phylogeny. For each of the six summary statistics (mean and variance for each of the three rates), we set up a linear mixed-effect model:

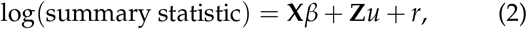

with inference method as a fixed-effect categorical predictor (effect sizes *β*), phylogeny as a random effect categorical predictor (*u*), and an error term *r*. **X** and **Z** are design matrices for the fixed and random effects. We visually checked that the residuals (*r*) were normally distributed and did not suffer from heteroscedasticity; phylogenies that violated these assumptions were excluded from this analysis. For each linear model, we tested if the least-square means of each pair of methods were statistically different using Tukey’s corrected p-value for multiple comparisons.

#### Location and magnitude of rate shifts

We additionally tested whether the methods inferred consistent locations and magnitudes of rate shifts, using the rooted version of the Kuhner-Felsenstein distance (Kuhner and Felsenstein, 1994). To do this, we first replaced branch lengths of each timetree with the posterior median rate estimate, from a given method, then scaled each branch by the total tree height. This produces a method-dependent tree with branch lengths that provide information regarding the magnitude and location of rate shifts but with identical topology. We calculated KF distances between the rescaled trees from each pair of methods; this distance is equivalent to the mean square error (MSE) given that the two trees being compared have the same topology, as they do in our analyses. For each tree and for each diversification parameter, we computed the mean square error among the different methods:

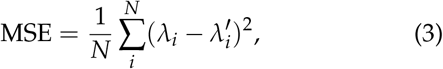

where *λ*_*i*_ (or similarly *μ*_*i*_, or (*λ*_*i*_ *−μ*_*i*_)) is the diversification rate parameter for branch *i*.

A large MSE tells us that the two methods being compared infer different rate magnitudes and/or rate shifts in different locations. A small MSE, however, indicates that the two methods give us similar results.

### Computation

We ran all diversification analyses either locally, on the Savio HPC at UC Berkeley, or using the CIPRES Science Gateway V. 3.3 (Miller et al., 2010).

We performed all comparison analyses in R version 3.6.0 (R Core Team, 2013). We performed data manipulation with the R `packages phytools`, (Revell, 2012), `tidyverse` (Wickham, 2017), `reshape2` (Wickham, 2012), readr (Wickham and Hester, 2020), `plyr`(Wickham, 2011b), and `coda` (Plummer et al., 2006). We generated plots with R `packages see` (Lü decke et al., 2021), `ggplot2` (Wickham, 2011a),`ggpubr` (Kassambara, 2018), `ggtree`(Yu et al., 2018), `ggsignif`(Ahlmann-Eltze, 2017), `ggExtra` (Attali and Baker, 2016), `cowplot`(Wilke, 2016) and `pdftools` (Ooms, 2020). We fit linear mixed models using the R package *lmer* (Bates et al., 2015) and obtain emeans estimates using the R package eemeans (Lenth, 2020). We additionally used `smacof` (Mair et al., 2022) and phangorn (Schliep, 2011) to perform MDS and to calculate RF and KF distances. Citations for R packages were generated with RefManageR (McLean, 2014).

## Results and Discussion

### Convergence

Our full dataset contains 76 chronograms from multicellular organisms, with 31 – 4161 extant tips, root ages of 4.9 – 349.8 MYA, and 0.014% – 100% of extant species sampled (Fig. S1A and Table S1). All methods converged for 43 trees (the “complete subset”; Fig. S1B). Of the methods tested, `LSBDS` had the most difficulty achieving MCMC convergence (it converged for 46 trees). All methods except `LSBDS` converged in 65 trees (the “partial subset”; Fig. S1C); PESTO directly computes the posterior mean and thus “convergence” does not apply. Trees that did not converge have poorer taxon sampling (i.e. the ratio of sampled species to total species richness; P-value = .039), older root ages (P-value = 0.0001), and greater branch length variance (P-value = .00006) than the converged trees, but sample size (number of tips) was not an important factor (P-value = .076; Fig. S2C–F). The branch length variance is consistent with the degree of spread between the KF and RF MDS analyses (Fig. S2A–B); the KF MDS—which accounts for branch lengths as well as topology—has a larger spread then the RF MDS. Overall these results fit with our intuitive understanding of the challenges in inferring shifts in diversification rates. We expect that older trees and trees with greater variation in branch lengths should undergo more rate shifts than younger trees and those with less variation in branch lengths. Thus inferring the diversification rates of these trees should be generally more challenging. These results suggest that users should be particularly attentive to MCMC convergence if their chronogram(s) are poorly sampled, old, or have a lot of branch-length variation, and especially so if they are using LSBDS. In these cases, even more so than usual, it is important to run each MCMC multiple times independently, to assess both stationarity and convergence.

### Comparison of Methods

#### Visualizing rates on trees

None of the 43 phylogenies in our “complete subset” had concordant estimates among all methods given our evaluation criteria (Fig. 1). For some phylogenies, the methods inferred similar shifts in net diversification (*e.g*., Fig. 1A–E), whereas for others the inferred shifts differed slightly (*e.g*., Fig.1F–J) or strongly (*e.g*., Fig. 1K–O).We would expect some differences when comparing different modeling approaches, as there are patterns to the differences in our results that can be attributed to the fundamental differences between the models. We illustrate these patterns using a few example phylogenies, which are representative of the patterns one will find when perusing the full set of trees in the supplemental materials (Supplemental Section S5).

**Figure 1:**
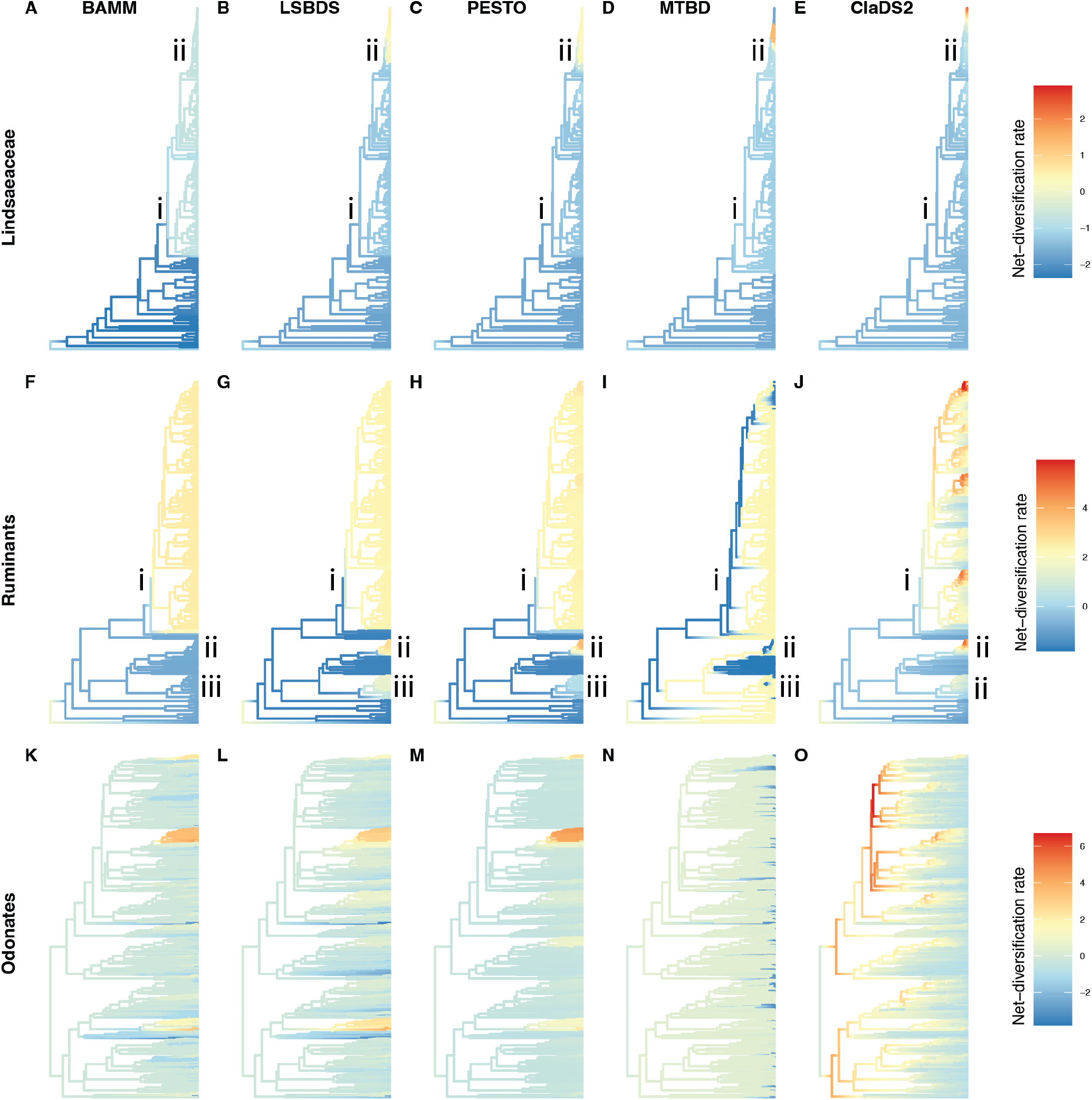
Three representative phylogenies with Z-transformed (mean centering and scaling to unit variance) posterior median estimate of net diversification painted on the branches. Columns show estimates from BAMM (A,F,K), LSBDS (B,G,L), PESTO (C,H,M), MTBD (D,I,N) and CLaDS2 (E,J,O). (A-E) Phylogeny of Lindsaeaceae (necklace ferns; Testo and Sundue, 2016), (F-J) Phylogeny of Ruminants (tetrapod; Toljagić et al., 2018), and (K-O) Phylogeny of Odonates (dragonflies and damselflies; Waller and Svensson, 2017). The rate values are in units of events per lineage per million years.

Occasionally two methods generally identified similar patterns. For example, `BAMM` and `LSBDS` identified a similar shift of about the same magnitude in speciation rates for some clades, *e.g*., the Lindsaeaceae (necklace ferns, clade i; Fig. 1A,B). Nonetheless, there are still differences between the two methods, *e.g*., a second nested rate shift in the LSBDS and PESTO estimates (clade ii).

In the ruminants (tetrapods) phylogeny (Fig. 1 F–J), we find that even for results that overall appear similar between methods, there are meaningful differences between their estimates. For example,`BAMM, LSBDS,` and `PESTO` inferred a shift around the ancestor of clade i, but `LSBDS` and `PESTO` also find approximately two more shifts (Fig. 1 G, clades ii and iii). Likewise, `MTBD` differs from the latter two as it infers several shifts in the largest clade and low net diversification rates on the backbone of that lineage (Fig. 1 I). Similarly, `CLaDS2` infers a slightly dif-ferent history from all of them, including multiple slow downs as well as an increase in net-diversification within clade i. `BAMM, LSBDS`, and `PESTO` identify a shift in approximately the same node (indicated by i) while MTBD infers many replicated increases in rate within clade i. LSBDS and PESTO infer the same shift, which is expected as they are based on the same underlying model and assumptions.

Multiple diversification shifts across a phylogeny is common to many of the `MTBD` trees (Fig. 1I,N; Barido-Sottani et al., 2020). This pattern is caused by the bimodal posterior distribution commonly inferred by this method (Barido-Sottani et al., 2020). Point-estimate summary statistics (*e.g*., posterior median) of these types of distributions are susceptible to small variation between the ancestor-descendant branches, which causes point estimates to switch between the two optima producing the rapid switching pattern (Fig. 1I,N).

Likewise, `CLaDS2` is the only time-dependent model in our analysis and thus is capable of detecting time-varying diversification patterns. Furthermore, the inherited speciation rate (*α*) only changes at cladogenic events, which results in many small changes at cladogenetic events, rather than the few large changes that characterize the other methods (Maliet et al., 2019; Maliet and Morlon, 2022). When *α* < 1 evolutionary slowdowns occur where the ancestral lineages have higher net diversification rates than the descendant lineages, a pattern observed in our data (Fig. 1O; Moen and Morlon, 2014).

On the other hand, BAMM, LSBDS, and PESTO are similar models and therefore we may expect them to infer similar diversification rates and shifts (Ronquist et al., 2021). While this is sometimes true (Figure 1 K,L), other times there are pronounced differences (Figure 1 F,G). This may be due to the well-known differences between these two models, namely assuming either rate shifts can (LSBDS and PESTO) or cannot (BAMM) occur on extinct lineages or unsampled lineages (Moore et al., 2016).

### Comparisons of rate estimates by method

To gain a global perspective of the differences between these models, we calculated two tree-wide summary statistics and distance metrics in order to compare these methods across the entire dataset (Fig. 2, S3).

**Figure 2:**
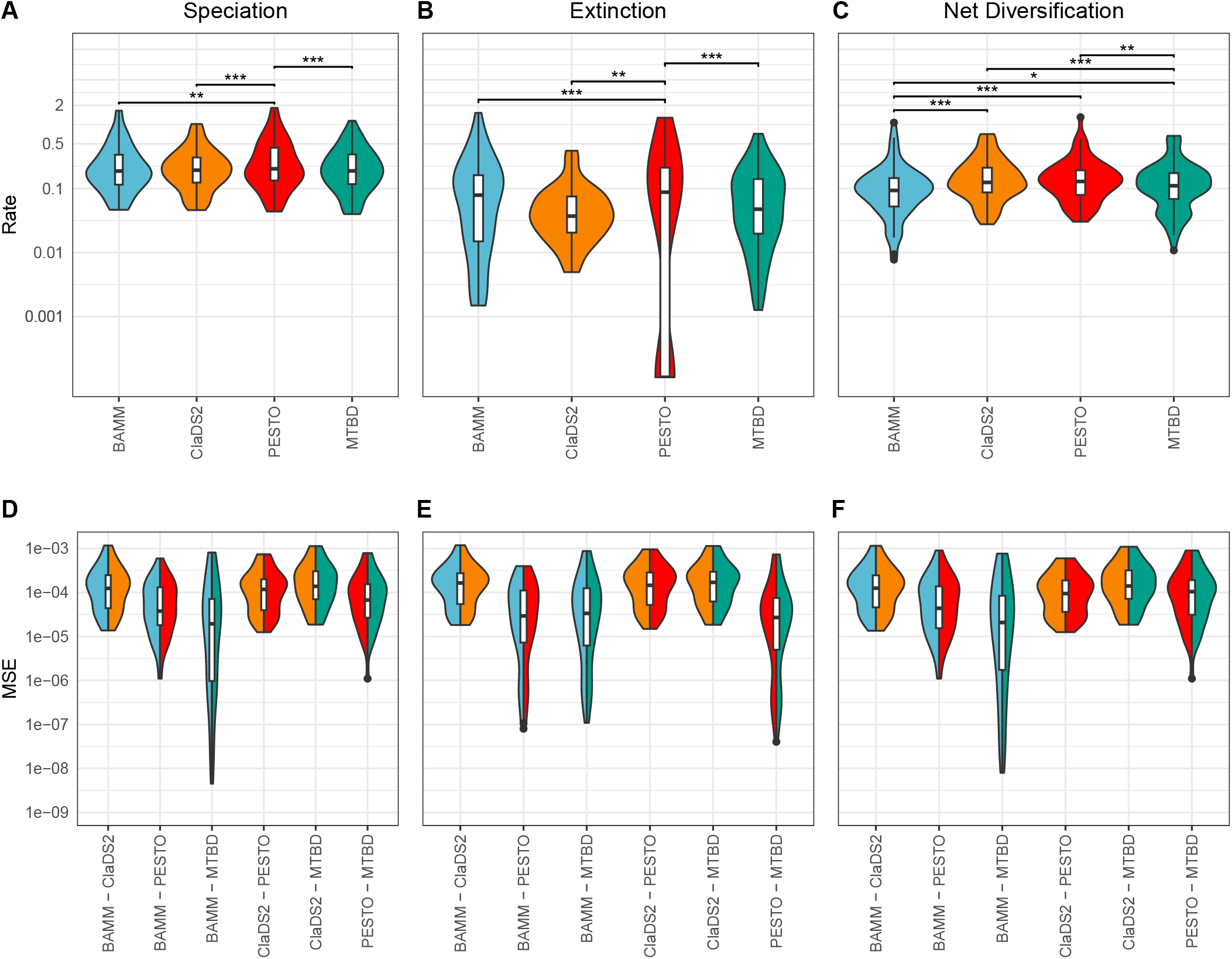
Comparison of tree-wide summary statistics across methods for the partial subset (n=65). (A–C) Tree-wide mean of posterior median estimates of the branch-specific rate parameters, plotted on a log scale. Asterisks correspond to the p-value of linear mixed model, calculated on the natural log of the rates (*: 0.05 *>* P-value *>* 0.01; **: 0.01 *>* P-value *>* 0.001; ***: 0.001 *>* P-value). (D–F) Pairwise mean squared error (MSE) between inference methods of phylogenies with branch lengths scaled by rates (speciation, extinction, and net diversification), plotted on a log scale. Split colors correspond to inference method color in A–C. Distributions closer to zero indicate that the inference methods produced more similar rate estimates, whereas higher values indicate greater dissimilarity. (D) MSE of speciation-scaled phylogenies; (E) MSE of extinction-scaled phylogenies; (F) MSE of net-diversification-scaled phylogenies.

We ran all comparisons on the complete subset (the 43 trees that converged for all methods) and on the partial subset (the 65 trees that converged for all methods except `LSBDS`). Comparisons between these two subsets reveal only one small difference (compare BAMM vs. MTBD -Fig. 2C vs. Supplemental Fig. S4C) and the most significant differences did not change. Given the large number of datasets that did not converge for `LSBDS` but converged for all other methods (unconverged datasets = 22) as well as the theoretical similarities between `PESTO` and `LSBDS`, we report the following results for the partial dataset (see Fig. S4A–F for summaries from the complete dataset).

We recover consistent differences in rate estimates among methods, particularly between `PESTO` (which is additionally standing-in for `LSBDS` in these comparisons, given that those two approaches share the same underlying model) and all other models; `CLaDS2` also was an outlier, albeit to a lesser extent (Table 1). In contrast, `BAMM` and `MTBD` tended to infer similar speciation and extinction rates. We find that tree-wide average speciation and extinction estimates of `PESTO` are statistically different from all other methods (Fig. 2A-B).

**Table 1:**
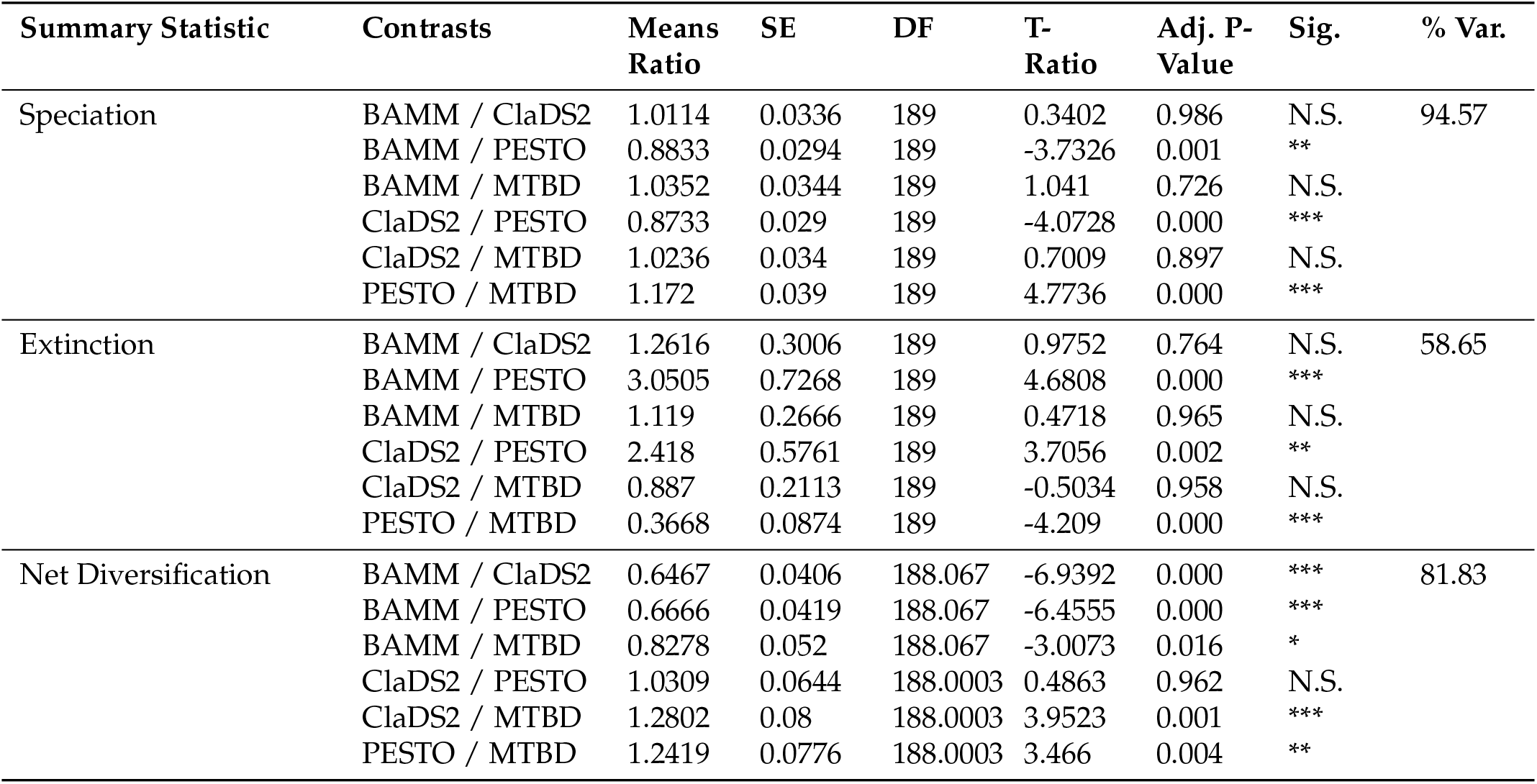
Post-hoc pairwise comparisons of inference methods using the tree-wide average of summary statistics: speciation, extinction, and net-diversification rates. Columns contain the summary statistics, contrasts of inference methods, the ratios of geometric means, standard errors, degrees of freedom, t-ratios, Tukey-adjusted p-values, significances, and the percent variances explained by the random effect.

While `PESTO` inferred higher tree-wide average speciation values, the magnitude of the differences is small (ratio of means < 1.2 for all significant contrasts; Table 1). Conversely, `PESTO` inferred lower tree-wide averages of extinction rates with larger magnitude changes (ratio of geometic means > 1.2; Table 1). The significant difference between `PESTO` and other methods holds for tree-wide average net-diversification as well, except for the comparison between `PESTO` and `CLaDS2` (Fig. 2C), which is not significant.

Additionally, `CLaDS2` tree-wide average net-diversification estimates are significantly different from `BAMM` and `MTBD` (Fig. 2C). A significant difference in net-diversification could be driven by the `CLaDS2` parameterization of extinction: extinction is not directly estimated in `CLaDS2`. Therefore the net diversification rates of `CLaDS2` are scaled speciation rates. Alternatively, the differences between methods could be due to the wider variance of netication estimates that both `BAMM` and `MTBD` have compared to `CLaDS2` (Fig. S3A–C). However, similar to tree-wide average speciation, the magnitude of difference between the contrast is not large (Table 1). There is also a weakly significant difference between speciation rates of `BAMM` and `MTBD` in our partial subset that was not found in the smaller complete subset.

All methods generally had comparable tree-wide average extinction-rate estimates with the exception of `PESTO`, which may infer much lower extinction rates for some trees than the other methods (though, on average, it infers higher extinction rates). The inference of extinction rate has been the subject of substantial debate, particularly in how failures to account for diversification shifts along extinct branches can impact the likelihood function (Moore et al., 2016; Rabosky et al., 2017). Regardless of the theoretical importance of correctly inferring extinction rates, we demonstrate that differences between extinction and speciation rates manifest in statistically different estimates of net diversification in empirical studies. Therefore, our results indicate that method-dependent tree-wide bias in diversification parameter inference may influence the interpretation of evolutionary shifts in diversification rates.

We find discrepancies between results derived from tree-wide summary statistics and our visual inspection of trees (see section “Visualizing rates on trees”). For example we find that `CLaDS2` and `PESTO` show no statistical difference in average net diversification (Fig. 2C). However, visual inspection of many trees suggests that `CLaDS2` and `PESTO` often differ greatly in the number and position of inferred rate shifts (*e.g*., Fig. 1). Conversely, `BAMM` and `LSBDS` often look very similar when we assess individual phylogenies and yet significantly differ when we compare speciation, extinction, and diversification averages S3A–C). This discrepancy reveals the difficulty of summarizing diversification rate estimates across phylogenies to reveal general patterns, and motivates the topology-informed rate comparisons, discussed in the following section.

### Location and magnitude of rate shifts

We also test whether the models recover similar locations and magnitudes of rate shifts by comparing the mean squared error (MSE) of branch rates; this metric bridges the discrepancies between the global metrics and the observed patterns across the trees (both described above; Fig. 2).

When quantifying differences in the location and magnitude of shifts in speciation and net diversification rates, `CLaDS2` differs the most (larger MSE), compared with the other methods (Fig. 2D,F) and it is by far the biggest outlier across the models when visually inspecting the trees (Fig. 1E,J,O). This result indicates that `CLaDS2` estimates differ strongly from those of the other methods in the degree of the shifts in infers, and in their location. This result is in contrast to the tree-wide averages presented above (see also Fig. 2A–C), where `CLaDS2` is unexceptional.

These results are also corroborated by analyses that take into account uncertainty in rate estimates (see Supplemental Section S4).

## Tools for Assessing Methods

Inferred rates generally differ depending on the analysis method; how then should an empirical biologist choose which method to use?

Our advice for empirical users is to take one of two paths. The first path is to carefully select a method based on the model assumptions. The methods presented in this analysis have theoretical differences in their approach, which appear to produce corresponding differences in results. For example, methods differ in whether shifts in diversification rates are allowed on extinct or unsampled lineages (`LSBDS, PESTO`, and the “exact `MTBD”` not tested here), whether diversification rates of each regime are drawn from a continuous distribution (`BAMM, MTBD`, and `CLaDS2`) or from a set of discrete rate categories (`LSBDS` and `PESTO`), and if shifts occur at cladogenetic events (`CLaDS2`) or along lineages (`BAMM, MTBD, LSBDS`, and `PESTO`). The models make additional assumptions, such as whether shifts in diversification rates affect the process-intrinsic parameters (the speciation and extinction rates) or transformations thereof (*e.g*., the net diversification or turnover rate) and whether shifts affect single parameters or combinations of parameters. These assumptions lead to notably different interpretations of how values change through time. Choice of method can be supported by taxon-specific data such as species distribution, fossil record, or phenotypic data (Morlon, 2014). Thus, users should also familiarize themselves with how these models parameterize and estimate diversification rates and ensure that these modeling choices reflect the user’s assumptions about biological processes.

The second path is to critically compare multiple methods when performing diversification analyses. We have shown that—despite the difference in models—in some cases multiple methods produce results with similar biological interpretations. To facilitate the adoption of this practice, we provide R code to easily visualize the results of multiple diversification-rate models across the same phylogeny: https://github.com/Jesusthebotanist/ CompDiv_processing_and_plotting.

## The Future of Diversification Analyses

The rise of methods aiming to identify shifts in diversification speaks to the importance of these analyses for understanding the drivers and impacts of important evolutionary events. However, we advocate for caution, for two reasons described below.

First, taking a cautious approach is especially important in light of the many potential problems with these methods, including the controversy surrounding the identifiability of birth-death models (Louca and Pennell, 2020, but see also Helmstetter et al. 2021; Legried and Terhorst 2022; Morlon et al. 2022; Kopperud et al. 2023b, among others).

Louca and Pennell (2020) presented a class of birthdeath models that are unidentifiable if the rate functions are time-varying (but homogeneous across lineages) and allowed to take any continuous shape. Nonetheless, hypothesis-driven approaches are not allowed to take any arbitrary shape. Since the rate shapes are designed to test diversification scenarios, defined *a priori*, it has been argued that this approach is less prone to the identifiability issue (Morlon et al., 2022). Even time-varying models that are more agnostic about prior hypotheses are not typically allowed to take any continuous rate shape. Among the “agnostic” models, the piecewise-constant model is the most eminent (Stadler, 2011; Magee et al., 2020), and this model has been proven to be asymptotically identifiable provided there are not too many pieces (Legried and Terhorst, 2022).

However, in spite of the non-identifiability, inferences of rapidly changing speciation and extinction rates are still typically robust (Kopperud et al., 2023b). The issue of non-identifiability remains to be investigated thoroughly in lineage-heterogeneous models. These models are more parameter-rich than their homogeneous cousins, and so we do not expect the issue of non-identifiability to be any simpler here.

Second, we caution against relying too heavily on the estimates from a single method without justifying the assumptions encoded into the model’s choices regarding parameterization and estimation, as we describe in detail in “Tools for Assessing Methods”.

The methods investigated in this paper vary in their underlying model and assumptions, but are theoretically related (Ronquist et al., 2021). Due to these model differences, we expect differences in inferences which, in turn, could translate into different biological interpretations. Using a set of empirically derived phylogenies, we show that this is true (Fig. 1): no two methods inferred the same shifts for any phylogeny. In some cases, methods generally agreed upon the location and timing of inferred shifts, but in other cases methods strongly disagreed. Method-dependent differences of individual trees were corroborated by tree-wide summary statistics, which indicated small but significant differences between methods (Fig. 2; Table 1). While these results hold up when we take into account the uncertainty in rate estimates, we also urge caution in relying too heavily on summary statistics and encourage users to carefully examine their posterior distributions, as 95% HPD intervals vary among methods and distributions may be bimodal, which may mislead common summary statistics (Fig. S6, see also Barido-Sottani et al., 2020).

Regardless, it is clear there will be a continued interest in using diversification analysis with a renewed appreciation for the complexities of these methods and the details of how rates are parameterized and estimated.

## Author Contributions

JM-G, MJS, and CMT designed and performed the analyses, JM-G, MJS, CMT, SH, BTK, WAF, CDS, and CJR wrote the manuscript.

## Acknowledgements

We thank Dr. Francisco Henao Diaz and Dr. Matthew Pennell for providing materials necessary to replicate their analyses. We thank Dr. Michael R. May, members of the Rothfels Lab *sensu lato*, and the Specht lab for comments on the manuscript, and we thank the Bonnie Lane Brain Trust for rapid brainstorming. Cornell University Statistical consulting unit provided valuable advice on linear mixed models. This research used the Savio computational cluster resource provided by the Berkeley Research Computing program at the University of California, Berkeley (supported by the UC Berkeley Chancellor, Vice Chancellor for Research, and Chief Information Officer). This material is based upon work supported by the National Science Foundation Graduate Research Fellowship Program under Grant No. DGE-1650441 for JMG and No. DGE-1752814 for CMT. Any opinions, findings, and conclusions or recommendations expressed in this material are those of the authors and do not necessarily reflect the views of the National Science Foundation. This material is based upon work supported by the NSF Postdoctoral Research Fellowships in Biology Program under Grant No. 2209159 to JM-G and No. 2109835 to CMT. JM-G was additionally supported by a Cornell Provost Diversity Fellowship. This work was supported by the Deutsche Forschungsgemeinschaft (DFG) Emmy Noether-Program (Award HO 6201/1-1 to S.H.) and by the European Union (ERC, MacDrive, GA 101043187). Views and opinions expressed are, however, those of the authors only and do not necessarily reflect those of the European Union or the European Research Council Executive Agency. Neither the European Union nor the granting authority can be held responsible for them. The authors declare no conflicts of interest.

## Data accessibility

All scripts, data and outputs can be found at on Dryad at (doi:10.6078/D18Q68) upon publication. A set of R functions to help user analyze outputs of studied Baysian methods can be found at (https://github.com/Jesusthebotanist/CompDiv_ processing_and_plotting)

## Notes

### Competing Interest Statement

The authors have declared no competing interest.

