## Supplemental Material for "Commonly used Bayesian diversification-rate models produce biologically meaningful differences on empirical phylogenies"

### Supplemental Materials

#### S1 Empirical datatsets

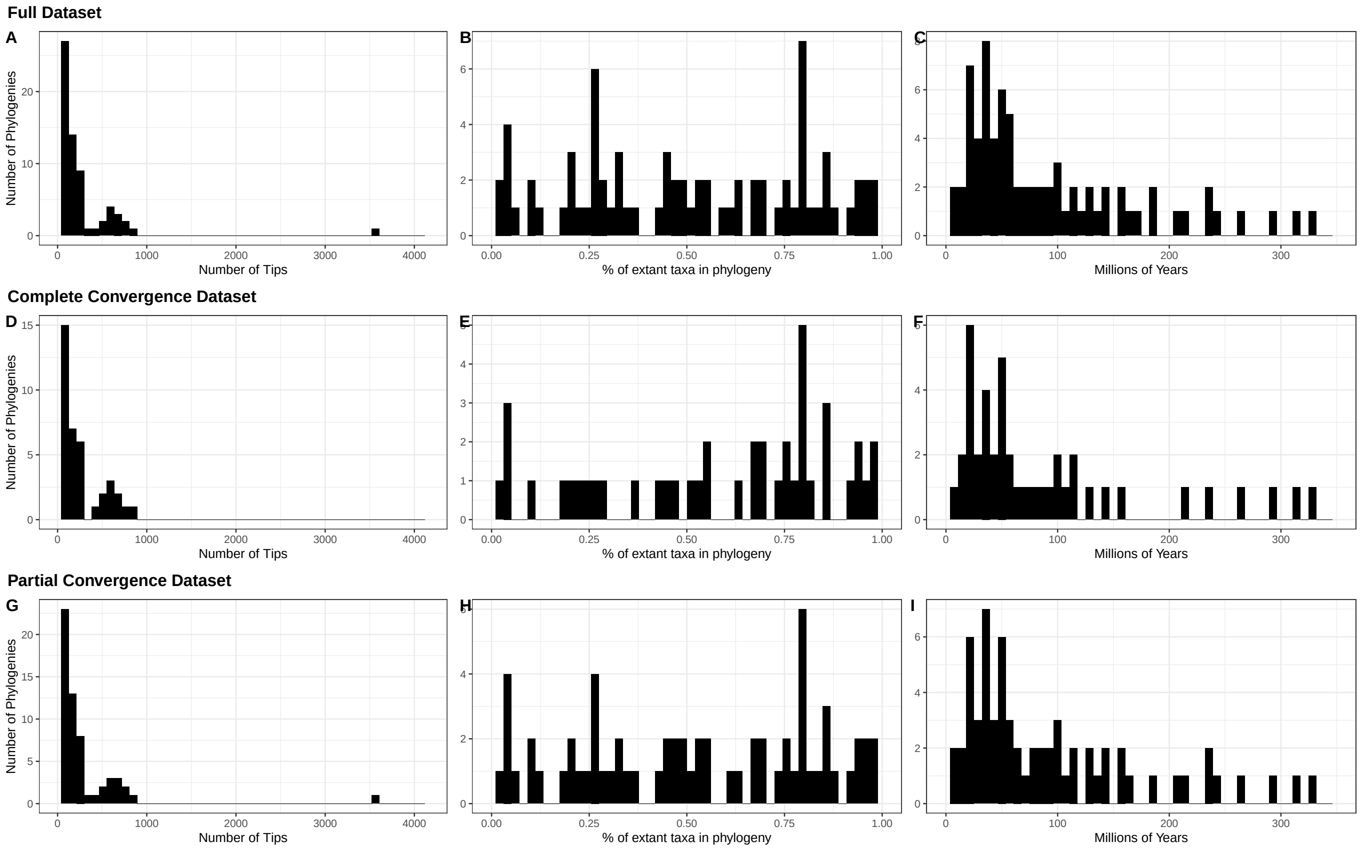


**Figure S1:** Summary statistics on the chronograms used in analyses from (Henao Diaz et al., 2019). A,D,G) Distribution of the number of tips per phylogeny. B,E,H) Distribution of incomplete-sampling percentage per phylogeny. C,F,I) Distribution of the age of the most recent comment ancestor. A-C) Summary statistics for the full dataset in this study. D-F) Summary statistics for the 'complete converged dataset' (only includes trees that reached convergence for all methods, n=43). G-I) Summary statistics for the 'Complete convergence dataset' (only includes tree that reached convergence for all methods except LSBDS, n = 65)

#### S2 Convergence

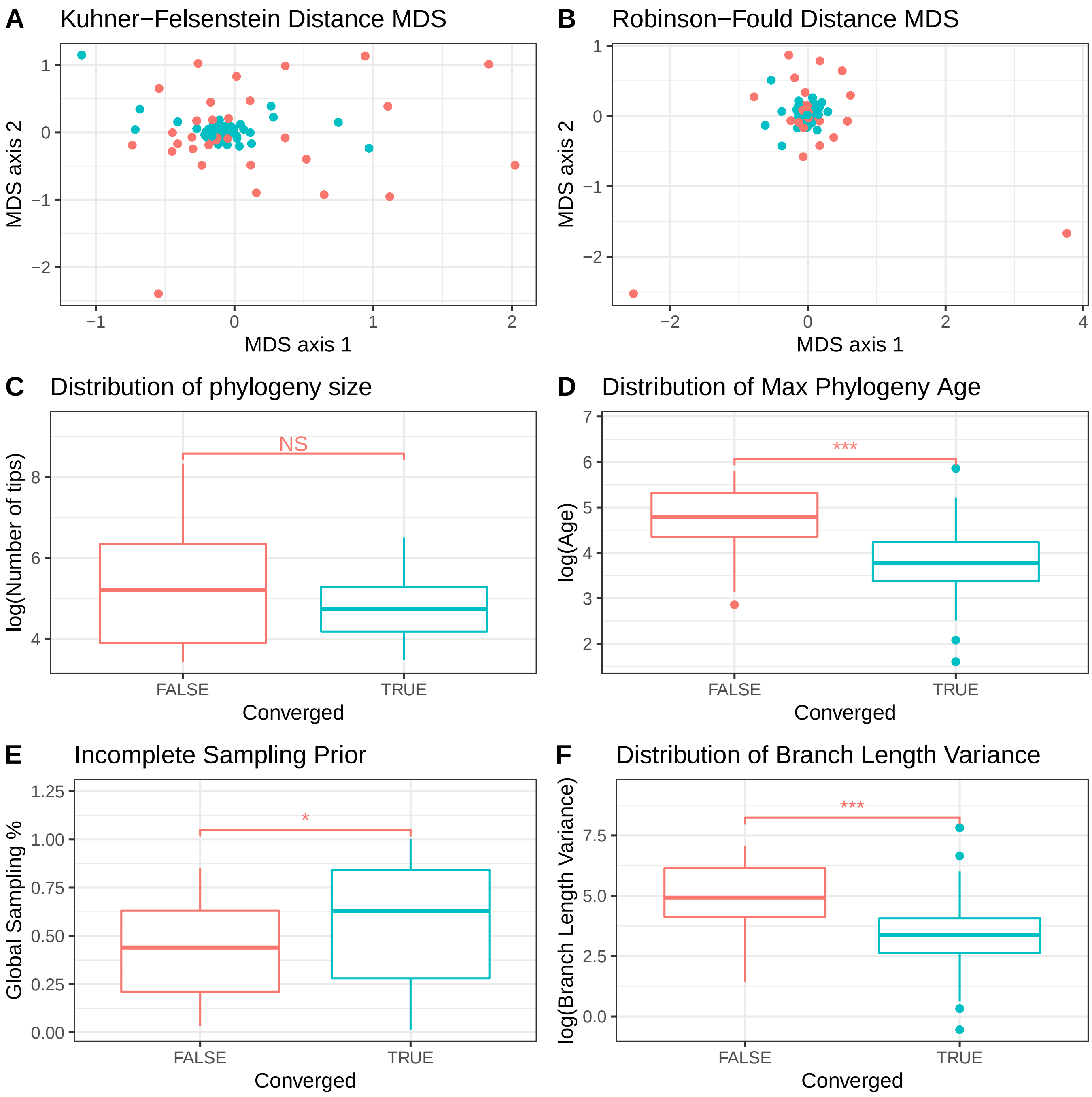


**Figure S2:** Comparison of convergence. Phylogeneies where converged was reached across all five methods n = 43 (Blue); Phylogeneies where converge was not reached for one or more methods n=33 (Red). A-B) Multidimensional scaling of phylogeny by, A) Kuhner-Felsenestin distance matrix and by, B) Robinson-Fould distance matrix. C) Size of the phylogeny in terms of number of tips. D) Age of the oldest nodes in the phylogeny. E) Percent incomplete sampling incorporated into diversification analysis. F) Variance of branch lengths of phylogenies. C-F) T-test performed between converged and unconverged phylogenies. P-value of T-test displayed (*: 0.05>P-value>0.01; **: 0.01>P-value>0.001; ***: 0.001>P-value).

**Table S1:** The chronograms that form part of the complete and partial convergence data subsets. The complete convergence subset includes trees that converged across all methods: LSBDS, BAMM, ClaDS2, and MTBD. The partial convergence subset includes trees that converged across all methods *but* LSBDS. The abbreviations are borrowed from the Henao Diaz et al. (2019) paper.

| Abbreviation | Clade | Complete Convergence Subset | Partial Convergence Subset | Citation |
|---|---|---|---|---|
| A2016 | Viperidae | TRUE | TRUE | Alencar et al. (2016) |
| A2017 | Costaceae | TRUE | TRUE | André et al. (2016) |
| B2016 | Onthophagus | TRUE | TRUE | Breeschoten et al. (2016) |
| BR2017 | Actinopterygii | FALSE | FALSE | Betancur-R et al. (2017) |
| C2015 | Ovalentaria | TRUE | TRUE | Campanella et al. (2015) |
| C2017 | Coronellini | TRUE | TRUE | Chen et al. (2017) |
| D2018 | Dytiscidae | FALSE | TRUE | Désamoré et al. (2018) |
| F2017 | Cichlidae | TRUE | TRUE | Missing in Henao Diaz et al. (2019) |
| G2011 | Ceanothus | TRUE | TRUE | Goldberg et al. (2011) |
| G2017 | Scolytinae | FALSE | FALSE | Gohli et al. (2017) |
| H2007 | Pinnipedia | TRUE | TRUE | Higdon et al. (2007) |
| H2016 | Cracidae | TRUE | TRUE | Hosner et al. (2016) |
| H2017 | Ctenitis | TRUE | TRUE | Hennequin et al. (2017) |
| HI2017 | Quercus | TRUE | TRUE | Hipp et al. (2018) |
| I2011 | Sebastes | TRUE | TRUE | Ingram (2011) |
| I2016 | Anolis | FALSE | TRUE | Ingram et al. (2016) |
| IL2017 | Heliconia | TRUE | TRUE | Iles et al. (2017) |
| JO2016 | Corvides | TRUE | TRUE | Jønsson et al. (2016) |
| L2012 | Coniferophyta; Pinidae | FALSE | FALSE | Leslie et al. (2012) |
| L2013 | Rhododendron section Vireya | TRUE | TRUE | Neupane et al. (2017) |
| L2014 | Agama | TRUE | TRUE | Leaché et al. (2014) |
| L2016 | Odonata | FALSE | TRUE | Letsch et al. (2016) |
| L2017 | Lobelioidae | TRUE | TRUE | Lagomarsino et al. (2017) |
| MC2016a | Balistidaea | TRUE | TRUE | McCord and Westneat (2016) |
| MC2016b | Monacanthidae | FALSE | TRUE | McCord and Westneat (2016) |
| N2017a | Spermacoceae | TRUE | TRUE | Neupane et al. (2017) |
| P2016 | Cephalotes | TRUE | TRUE | Price et al. (2016) |
| P2017 | Testudinata | FALSE | FALSE | Pereira et al. (2017) |
| PB2014 | Squamata | FALSE | FALSE | Pyron and Burbrink (2014) |
| pg1646 | Pinnipedia | FALSE | TRUE | Higdon et al. (2007) |
| pg1953 | Furnariidae | TRUE | TRUE | Derryberry et al. (2011) |
| pg2575 | Passeriformes | TRUE | TRUE | Barker et al. (2013) |
| pg2576 | Actinopterygii | FALSE | FALSE | Betancur-R et al. (2017) |
| pg2659 | Otophysi | TRUE | TRUE | Chen et al. (2013) |
| pg2689 | Lupinus | TRUE | TRUE | Drummond et al. (2012) |
| pg2850 | Columbidae | TRUE | TRUE | Cibois et al. (2014) |
| pg2853 | Trochilidae | TRUE | TRUE | Désamoré et al. (2018) |
| PRE2017 | Temnothrorax | TRUE | TRUE | Prebus (2017) |
| R2013 | Mormoopidae & Phyllostomidae | TRUE | TRUE | Rojas et al. (2013) |
| R2018 | Phymaturus | FALSE | TRUE | Reaney et al. (2018) |
| S2012 | Carnivora | FALSE | TRUE | Slater et al. (2012) |
| S2014 | Poaceae | FALSE | FALSE | Spriggs et al. (2014) |
| S2015 | Galliformes | TRUE | TRUE | Stein et al. (2015) |
| S2018.01 | Elasmobranchii | FALSE | FALSE | Stein et al. (2018) |
| SA2016 | Columbiformes | TRUE | TRUE | Soares et al. (2016) |
| SamplePrimates | Primates | TRUE | TRUE | Missing in Henao Diaz et al. (2019) |
| SampleWhale | Ballenidae | TRUE | TRUE | Steeman et al. (2009) |
| SH2015 | Chiroptera | FALSE | TRUE | Shi and Rabosky (2015) |
| SH2016 | Pectinidae | FALSE | TRUE | Sherratt et al. (2016) |
| SO2017 | Aedes | TRUE | TRUE | Soghigian et al. (2017) |
| ST2017 | Astacoidea & Parastacoidea | FALSE | TRUE | Stern et al. (2017) |
| SW2014 | Pergidae | TRUE | TRUE | Schmidt and Walter (2014) |
| T2015 | Myrtaceae | TRUE | TRUE | Thornhill et al. (2015) |
| T2018 | Cetartiodactyla | TRUE | TRUE | Toljagić et al. (2018) |
| TE201602 | Aspleniaceae | FALSE | TRUE | Testo and Sundue (2016) |
| TE201603 | Athyriaceae | TRUE | TRUE | Testo and Sundue (2016) |
| TE201604 | Blechnaceae | TRUE | TRUE | Testo and Sundue (2016) |
| TE201607 | Cyatheaceae | FALSE | TRUE | Testo and Sundue (2016) |
| TE201608 | Cystopteridaceae | FALSE | TRUE | Testo and Sundue (2016) |
| TE201610 | Dennstaedtiaceae | FALSE | TRUE | Testo and Sundue (2016) |
| TE201614 | Dryopteridaceae | FALSE | FALSE | Testo and Sundue (2016) |
| TE201616 | Gleicheniaceae | FALSE | TRUE | Testo and Sundue (2016) |
| TE201617 | Hymenophyllaceae | FALSE | TRUE | Testo and Sundue (2016) |
| TE201619 | Lindsaeaceae | TRUE | TRUE | Testo and Sundue (2016) |
| TE201620 | Lomariopsidaceae | FALSE | TRUE | Testo and Sundue (2016) |
| TE201624 | Marattiaceae | FALSE | TRUE | Testo and Sundue (2016) |
| TE201625 | Marsileaceae | FALSE | TRUE | Testo and Sundue (2016) |
| TE201631 | Ophioglossaceae | FALSE | TRUE | Testo and Sundue (2016) |
| TE201634 | Polypodiaceae | FALSE | FALSE | Testo and Sundue (2016) |
| TE201636 | Pteridaceae | TRUE | TRUE | Testo and Sundue (2016) |
| TE201641 | Tectariaceae | TRUE | TRUE | Testo and Sundue (2016) |
| TE201642 | Thelypteridaceae | FALSE | TRUE | Testo and Sundue (2016) |
| UC2015 | Rhinantheae | FALSE | TRUE | Uribe-Convers and Tank (2015) |
| VA2017b | Myrteae | FALSE | FALSE | Vasconcelos et al. (2017) |
| W2013 | Vitis | TRUE | TRUE | Wan et al. (2013) |
| WS2017.01 | Anisoptera | TRUE | TRUE | Waller and Svensson (2017) |
| WS2017.02 | Zygoptera | FALSE | FALSE | Waller and Svensson (2017) |

### S3 Methods Comparison

**Table S2:** Post-hoc pairwise comparisons of variance summary statistics

| Summary Statistic | Contrasts | Ratio of Geometric Means | Standard error | Degrees of freedom | t-ratio | Tukey adjusted p-value | Significance | % Variance Explained by Random Effect |
|---|---|---|---|---|---|---|---|---|
| Speciation Variance | BAMM / ClaDS2 | 7.00E-04 | 4.00E-04 | 189 | -11.8382 | 0 | *** | 60.1183 |
| | BAMM / PESTO | 0.0033 | 0.0021 | 189 | -9.2785 | 0 | *** | |
| | BAMM / MTBD | 0.4084 | 0.2509 | 189 | -1.4577 | 0.465212569 | N.S. | |
| | ClaDS2 / PESTO | 4.8196 | 2.9612 | 189 | 2.5597 | 0.054320203 | N.S. | |
| | ClaDS2 / MTBD | 588.653 | 361.6703 | 189 | 10.3805 | 0 | *** | |
| | PESTO / MTBD | 122.1364 | 75.041 | 189 | 7.8208 | 2.13E-12 | *** | 60.1183 |
| Extinction Variance | BAMM / ClaDS2 | 0.0041 | 0.0041 | 189 | -5.5293 | 6.32E-07 | *** | |
| | BAMM / PESTO | 15.2876 | 15.1724 | 189 | 2.7478 | 0.033046884 | * | |
| | BAMM / MTBD | 1.2546 | 1.2451 | 189 | 0.2285 | 0.99578196 | N.S. | |
| | ClaDS2 / PESTO | 3694.5857 | 3666.7289 | 189 | 8.277 | 1.17E-13 | *** | |
| | ClaDS2 / MTBD | 303.192 | 300.9059 | 189 | 5.7578 | 2.03E-07 | *** | |
| | PESTO / MTBD | 0.0821 | 0.0814 | 189 | -2.5193 | 0.060168773 | N.S. | |
| Diversification Variance | BAMM / ClaDS2 | 5.00E-04 | 3.00E-04 | 189 | -11.6464 | 0 | *** | 54.9987 |
| | BAMM / PESTO | 0.0014 | 9.00E-04 | 189 | -9.9236 | 0 | *** | |
| | BAMM / MTBD | 0.401 | 0.2649 | 189 | -1.3832 | 0.511419639 | N.S. | |
| | ClaDS2 / PESTO | 3.1214 | 2.0623 | 189 | 1.7228 | 0.314734182 | N.S. | |
| | ClaDS2 / MTBD | 880.9356 | 582.0436 | 189 | 10.2632 | 0 | *** | |
| | PESTO / MTBD | 282.2281 | 186.4711 | 189 | 8.5404 | 7.44E-15 | *** | |

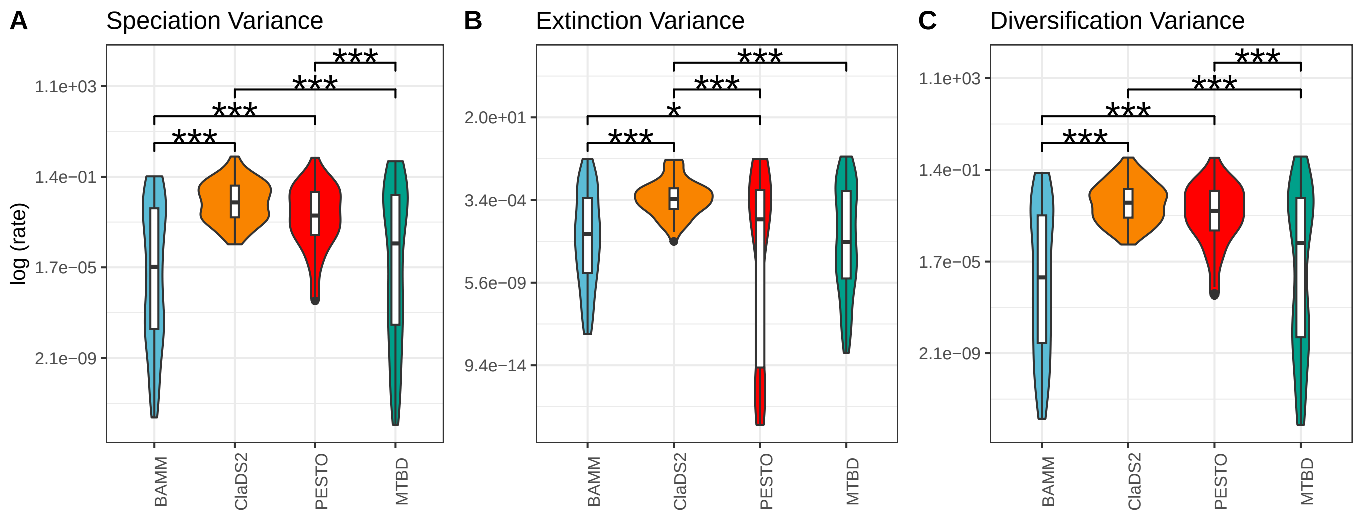


**Figure S3:** Comparison of summary statistics across the Partial Convergence dataset which excludes LSBDS. The distributions of averages is shown in Fig. 2 A–C in the main text. (A) Variances of speciation rates, (B) Variance of extinction rate, (C) Variance of net diversification rate. P-value of linear mixed model displayed (*: 0.05>P-value>0.01; **: 0.01>P-value>0.001; ***: 0.001>P-value)

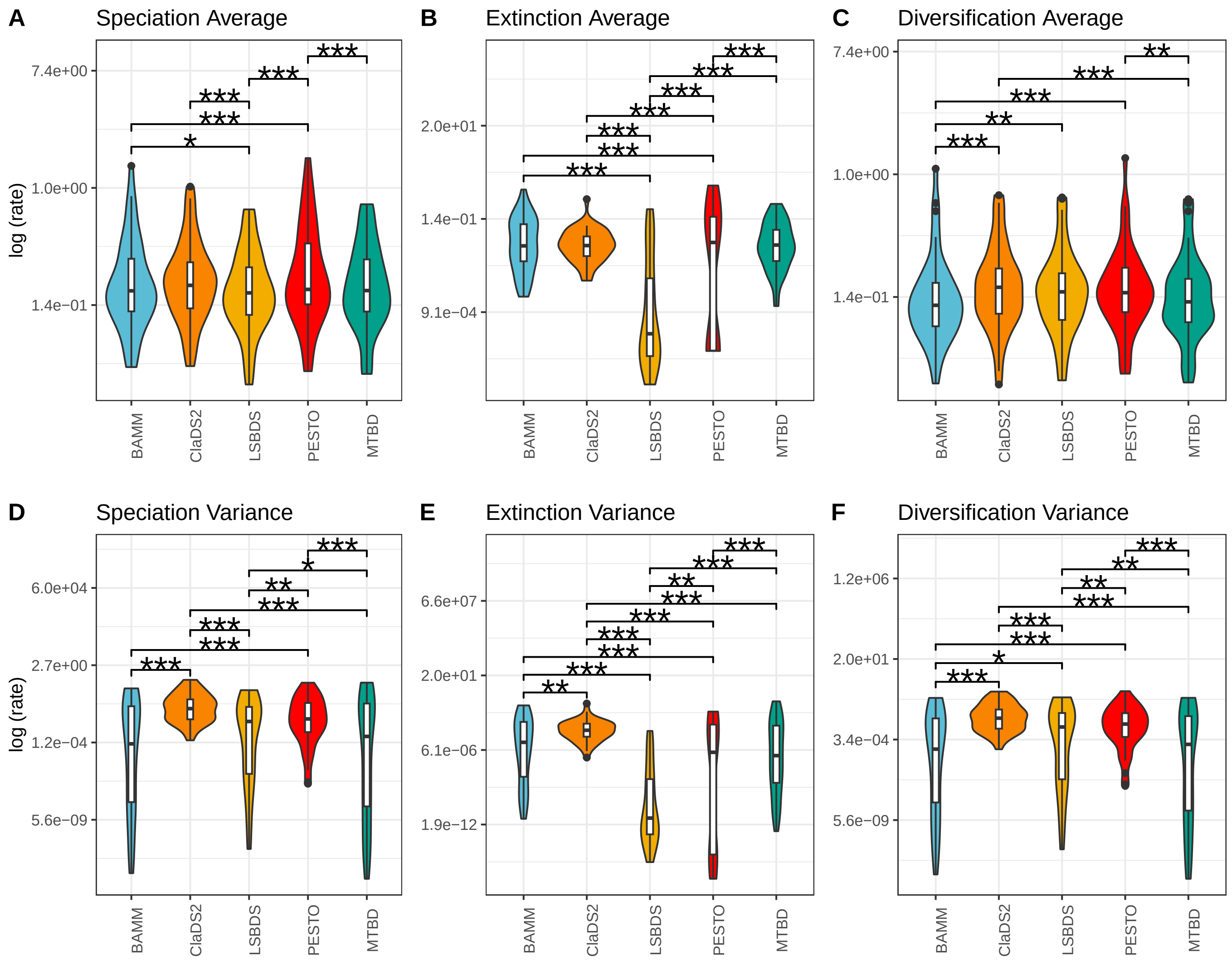


**Figure S4:** Comparison of summary statistics across methods for the complete convergence subset which includes all methods. (A) Average speciation rate, (B) Average extinction rate, (C) Average net diversification rate, (D) Variances of speciation rates, (E) Variance of extinction rate, (F) Variance of net diversification rate. P-value of linear mixed model displayed (*: 0.05>P-value>0.01; **: 0.01>P-value>0.001; ***: 0.001>P-value).

### S4 Quantifying Uncertainty in Rate Estimates

#### S4.1 Uncertainty Methods

To quantify how methods differ in terms of the uncertainty of their branch rate estimates, we calculated the 95% HPD interval for all net diversification branch estimates for all trees. We used the complete convergence subset in order to include LSBDS, however our results hold for the partial subset as well. This analysis excludes PESTO, as PESTO directly infers the posterior mean and thus does not estimate a posterior distribution from which an HPD interval could be inferred. Using the HPD intervals, we calculated two metrics. First, we calculated the HPD interval overlap ratio (HPD IOR) between pairs of methods by taking the length of overlap between two HPD intervals and dividing it by the length of the union of the two HPD intervals, where the union is the total "distance" spanned by the two intervals. An HPD IOR of 1 is signifies complete overlap while 0 signifies no overlap. This measure quantifies the similarity of HPD intervals across all branches and methods. Second, we quantified the average precision of the uncertainty—in other words, the breadth of the 95% HPD interval for estimates from `LSBDS`, `CLaDS2`, `BAMM`, and `MTBD`. We used a linear mixed model to test for statistical differences between these interval breadths:

$$\log(95\%\text{ HPD interval breadth}) = \mathbf{X}\beta + \mathbf{Z}i + r, \tag{S1}$$

with inference method as a fixed-effect categorical predictor (effect sizes $\beta$), branch as a random effect categorical predictor ($i$), and an error term $r$, we tested if the least-squares means of each pair of methods were statistically different using Tukey's corrected p-value for multiple comparisons.

#### S4.2 Uncertainty results

When comparing the HPD IOR between methods we observed the same general trends that we observed in our MSE result (Fig. S5, Fig. 2B). `CLaDS2` differed the most, on average a lower overlap ration when compared to all other methods comparisons (S5. `BAMM-LSBDS` and `BAMM-MTBD` show high overlap, and `MTBD-LSBDS` shows higher overlap than any `CLaDS2` comparison but is lower than `BAMM-LSBDS` and `BAMM-MTBD`.

On average HPD IOR for all comparison was less than 20% (S5). This indicates that on average most of the methods are inferring different rates for a particular branch. However, for some branches—excluding `CLaDS2-MTBD` and `CLaDS2-MTBD`—there was nearly complete overlap in HPD intervals (*i.e.*, HPD IOR near 1) suggesting that in these cases the methods infer nearly the same rate and uncertainty range. Branch-dependent agreement in method is generally consistent with our qualitative assessment of painted trees (Fig. 1 in the main text). There were certain trees where methods inferred the same values, but generally for most phylogenies the methods inferred different rates and rate shifts.

When considering HPD interval length in net-diversification estimates, a post-hoc pairwise comparison of methods found that significant difference amongst all contrast except for `LSBDS` and `MTBD` (Table S3). Interestingly, `CLaDS2` has 95% HPD intervals are at least 1.97 standard deviation longer than all other methods (ClaDS2 vs. BAMM: d = 2.3240, SD = 0.00994; ClaDS2 vs. MTBD: d = 2.0002, SD = 0.00994; ClaDS2 vs.LSBDS: d = 1.9778, SD = 0.00994) This was surprising given that comparisons that included `CLaDS2` had the lowest overlap. Collectively, these corroborate our results based on summary statistics of method dependent difference. Moreover, our results highlight that often times methods are inferring partially overlapping distribution and therefore the entire posterior distribution should be examined when drawing conclusion.

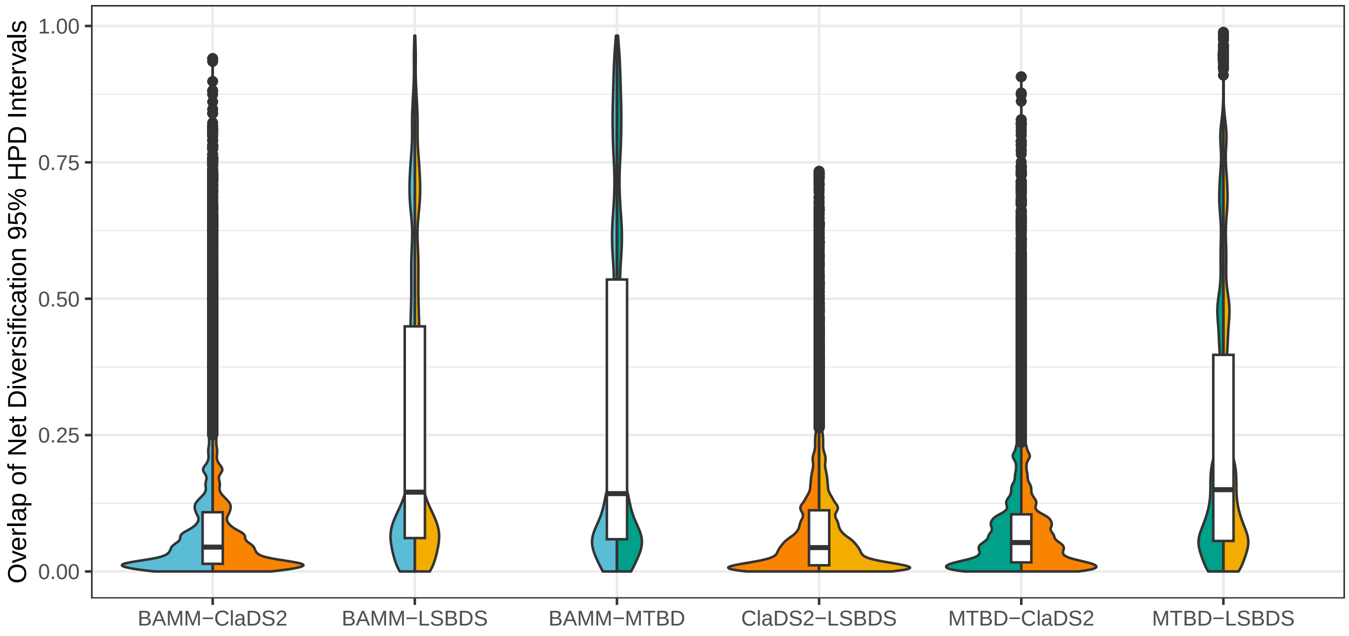


**Figure S5:** The extent to which the 95 % HPD intervals for net-diversification estimates overlap of the partial convergence subset. Violin plots correspond to pairs of methods (*eg.* BAMM and LSBDS).

**Table S3:** Post-hoc pairwise comparisons of inference methods on HPD interval length. Peformed on the ”complete convergence dataset” Columns contain the, contrasts of inference methods, the ratios of geometric means, standard errors, degrees of freedom, t-ratios, Tukey-adjusted p-values, significances.

| Contrasts | Means Ratio | SE | DF | Z-Ratio | Adj. P-Value | Sig. |
|---|---|---|---|---|---|---|
| ClaDS2 / BAMM | 4.431 | 0.02822 | Inf | 233.787 | <.0001 | *** |
| ClaDS2 / MTBD | 3.601 | 0.02293 | Inf | 1 201.219 | <.0001 | *** |
| ClaDS2 / LSBDS | 3.550 | 0.02260 | Inf | 198.960 | <.0001 | *** |
| BAMM / MTBD | 0.813 | 0.00518 | Inf | -32.568 | <.0001 | *** |
| BAMM / LSBDS | 0.801 | 0.00510 | Inf | -34 827 | <.0001 | *** |
| MTBD / LSBDS | 0.986 | 0.00628 | Inf | -2.258 | 0.1080 | *** |

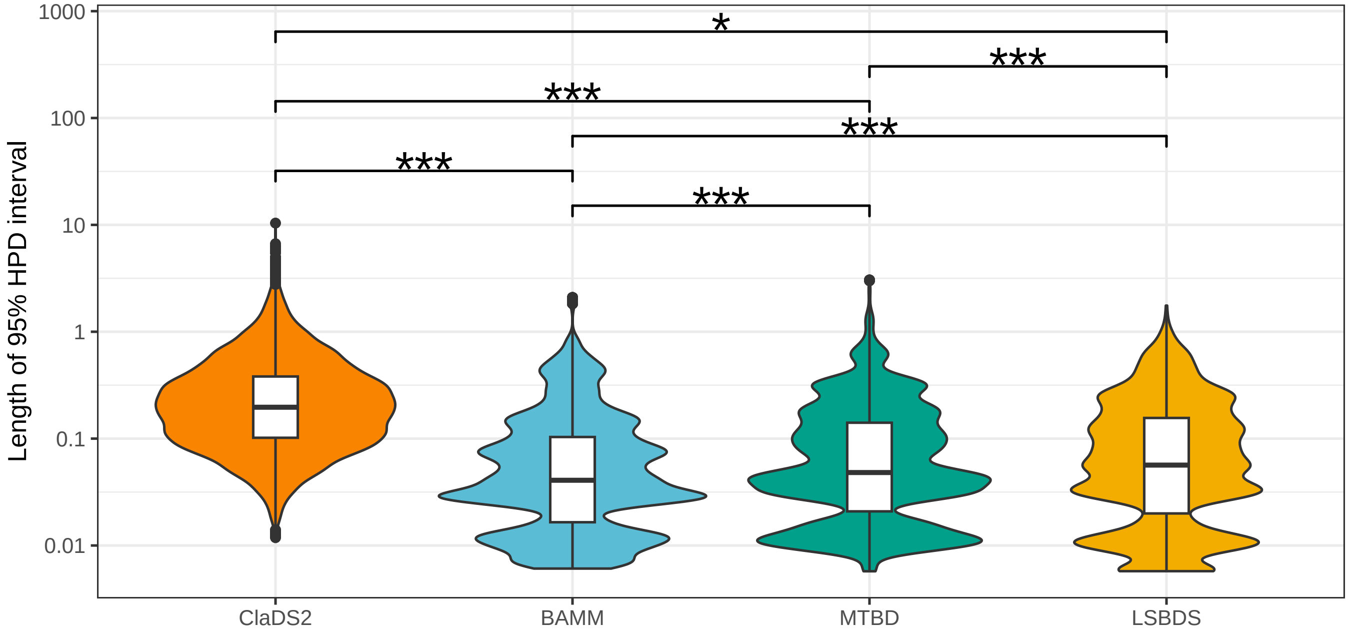


**Figure S6:** Comparison of HPD length of net-diversification across all branches of all trees in the partial convergence subset. P-value of linear mixed model with branch as random effect displayed (*: 0.05>P-value>0.01; **: 0.01>P-value>0.001; ***: 0.001>P-value).

#### S5 Results for Individual Chronograms

Below, we include a figure for each chronogram in the partial convergence subset of the speciation, extinction, and net-diversification rates for each inference method. If the figure does not include all inference methods, that chronogram did not converge using the missing method.

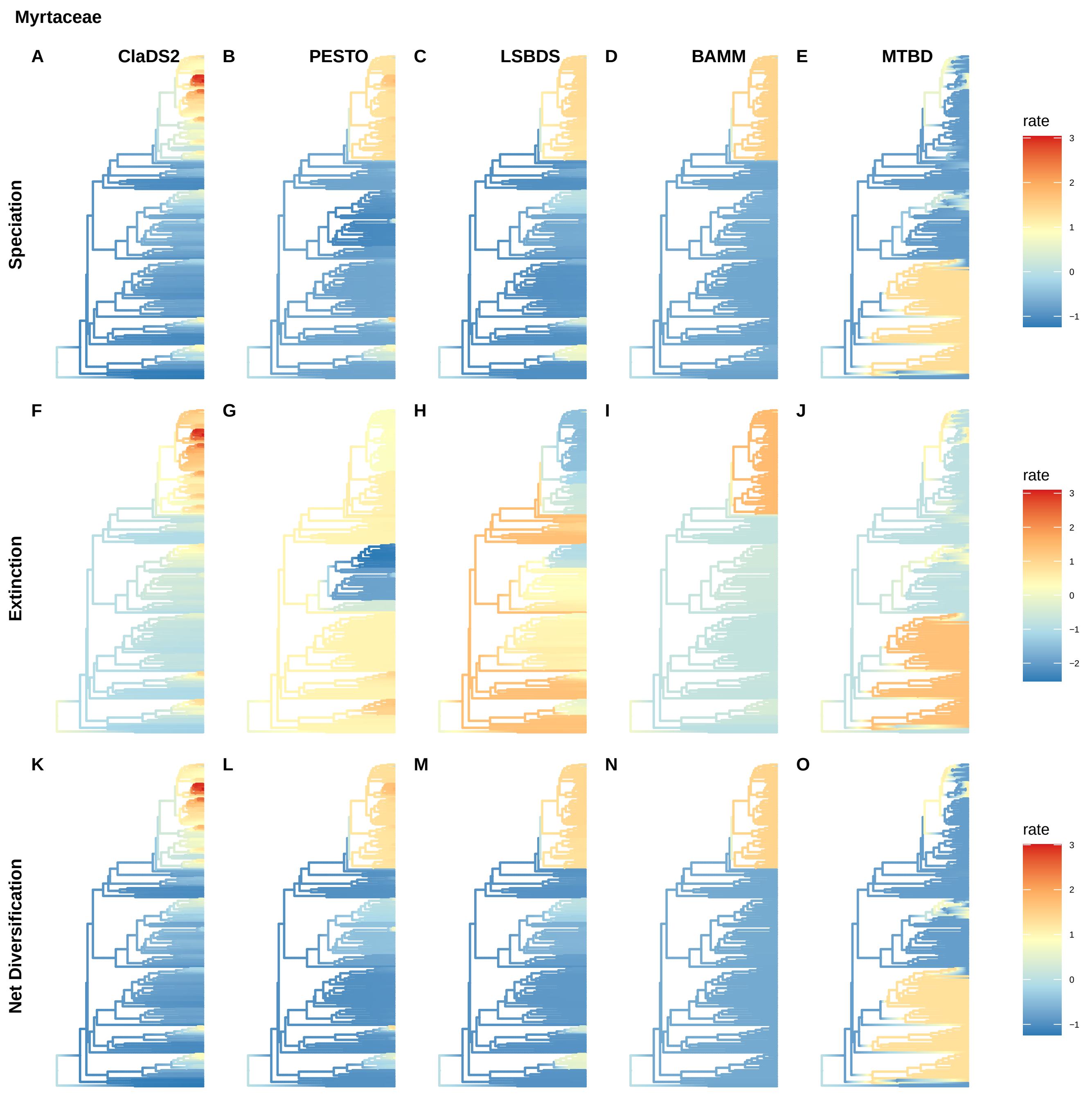
Myrtaceae
A
ClaDS2
B
PESTO
C
LSBDS
D
BAMM
E
MTBD
F
G
H
I
J
K
L
M
N
O
Speciation
Extinction
Net Diversification
rate
3
2
1
0
−1
rate
3
2
1
0
−1
−2
rate
3
2
1
0
−1

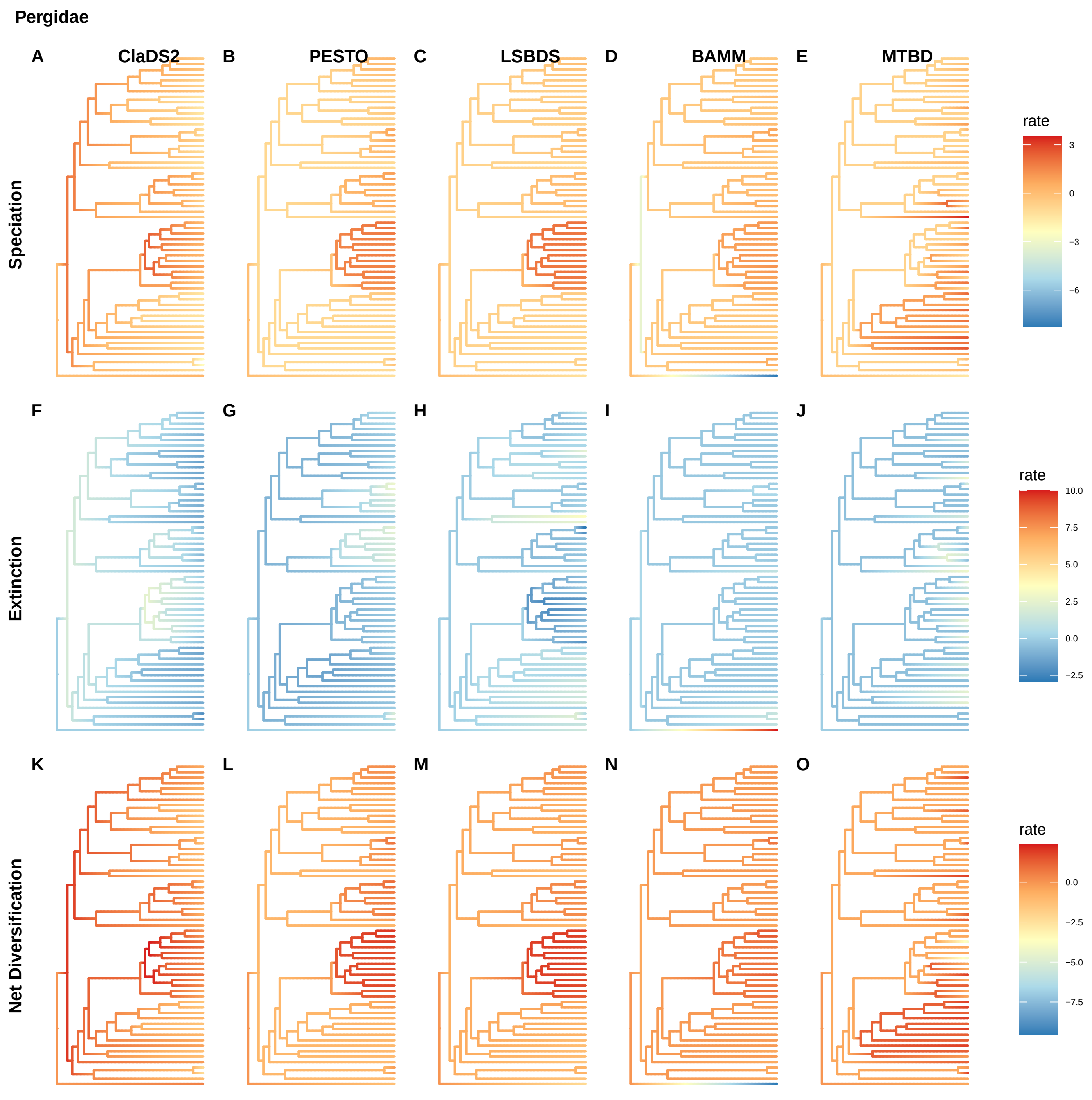
Pergidae
A
ClaDS2
B
PESTO
C
LSBDS
D
BAMM
E
MTBD
Speciation
rate
3
0
−3
−6
F
G
H
I
J
Extinction
rate
10.0
7.5
5.0
2.5
0.0
−2.5
K
L
M
N
O
Net Diversification
rate
0.0
−2.5
−5.0
−7.5

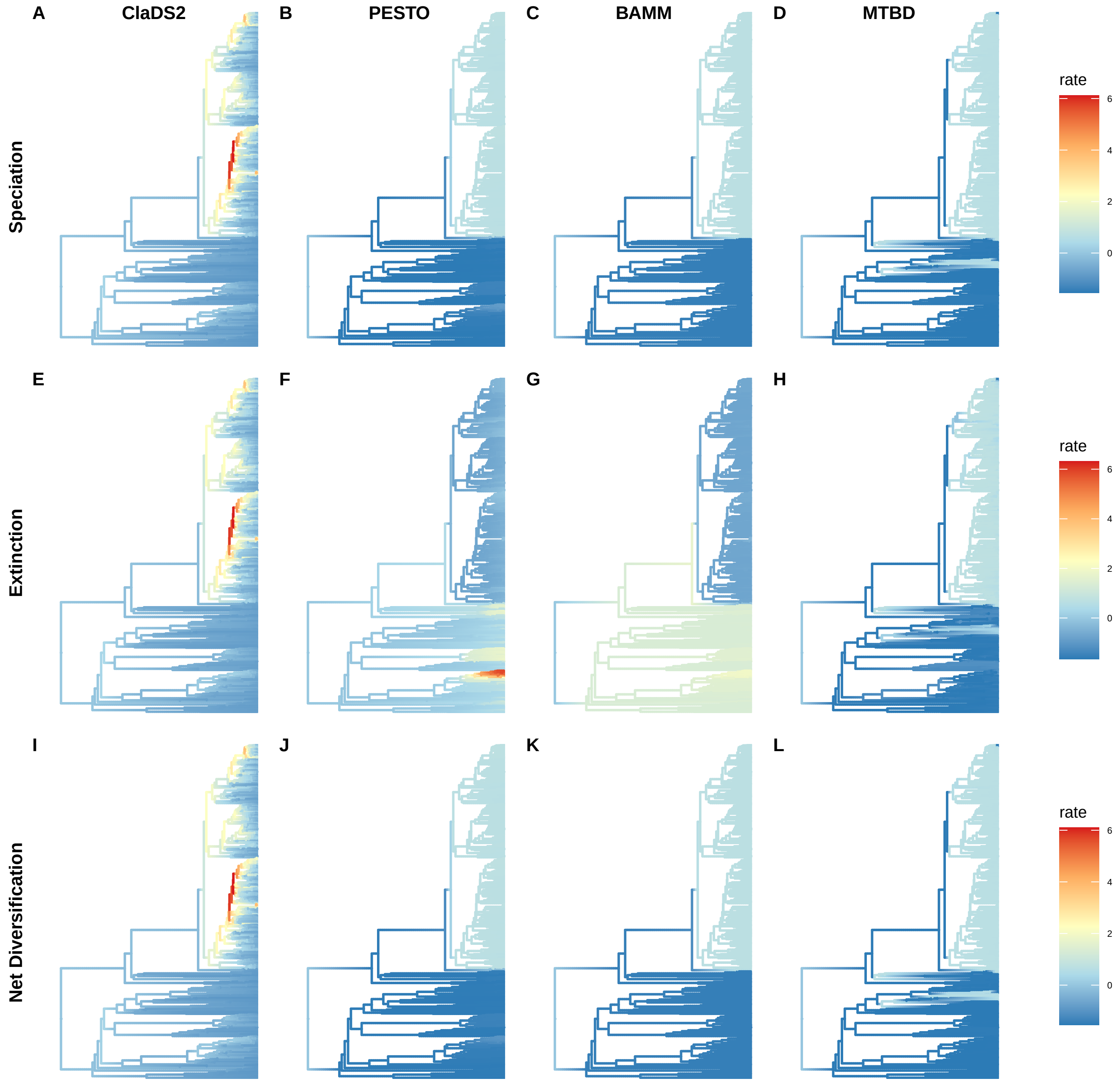
A
ClaDS2
B
PESTO
C
BAMM
D
MTBD
Speciation
rate
6
4
2
0
E
F
G
H
Extinction
rate
6
4
2
0
I
J
K
L
Net Diversification
rate
6
4
2
0

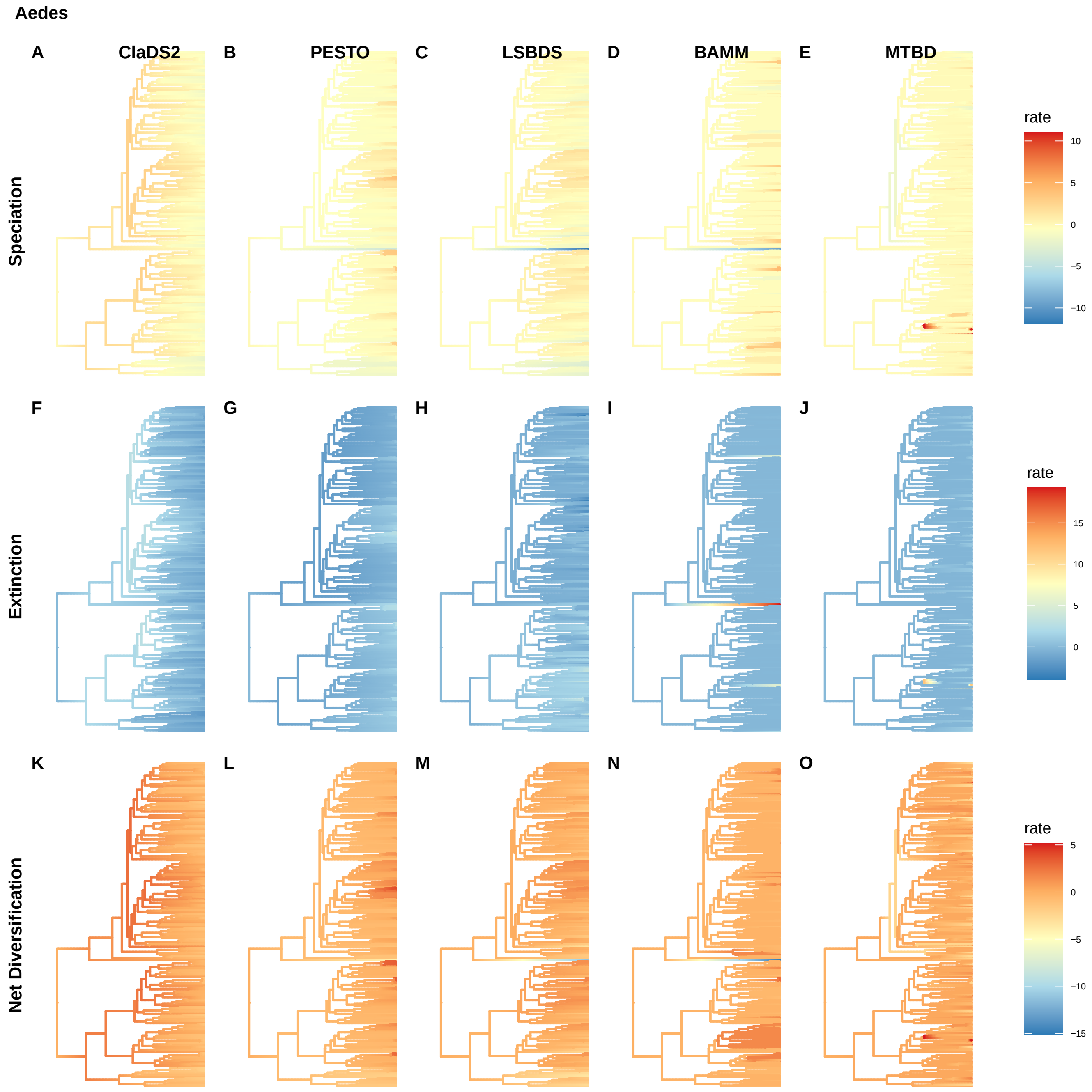
Aedes
A
ClaDS2
B
PESTO
C
LSBDS
D
BAMM
E
MTBD
F
G
H
I
J
K
L
M
N
O
Speciation
Extinction
Net Diversification
rate
10
5
0
−5
−10
rate
15
10
5
0
rate
5
0
−5
−10
−15

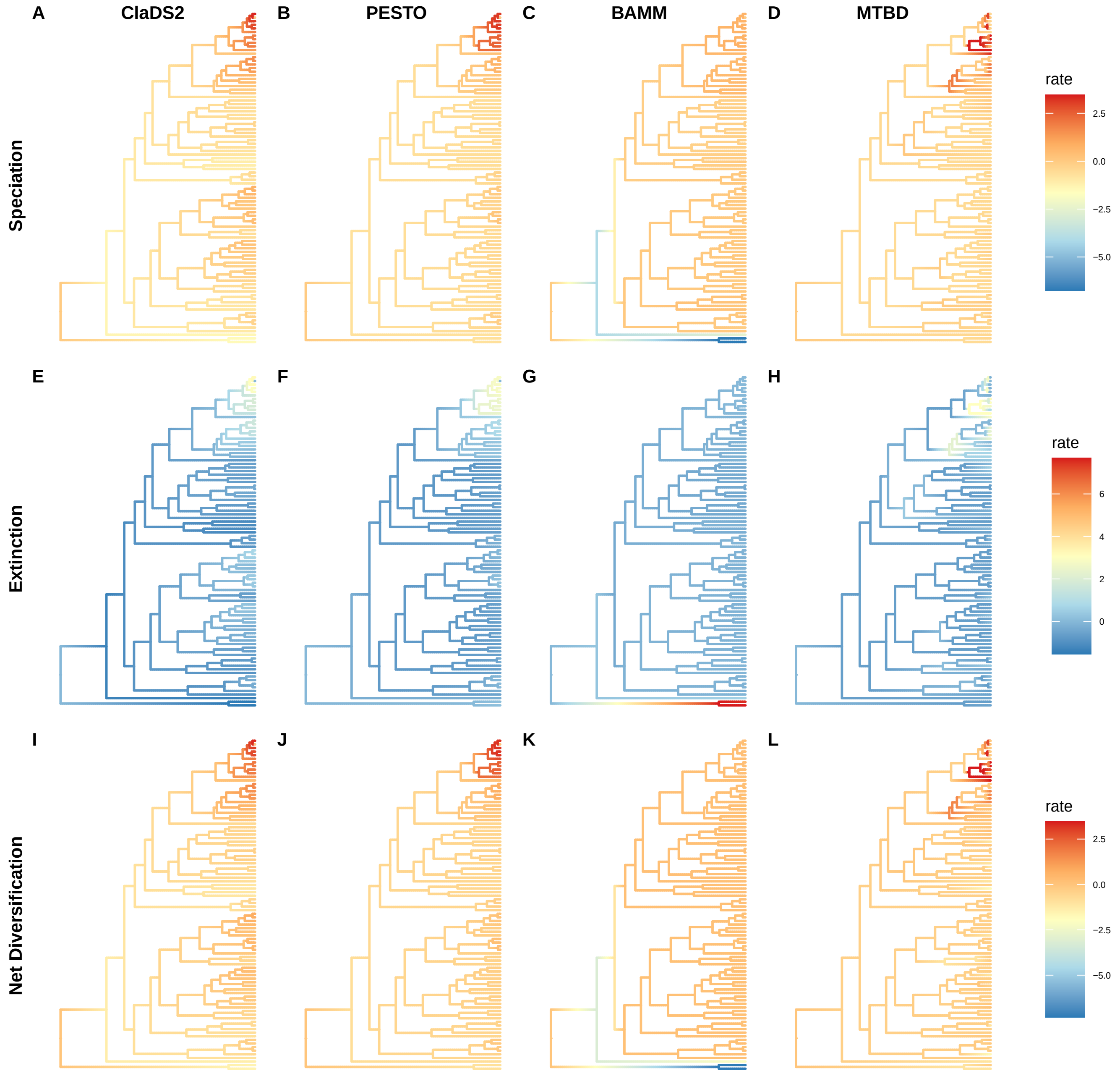
A
ClaDS2
B
PESTO
C
BAMM
D
MTBD
Speciation
rate
2.5
0.0
−2.5
−5.0
E
F
G
H
Extinction
rate
6
4
2
0
I
J
K
L
Net Diversification
rate
2.5
0.0
−2.5
−5.0

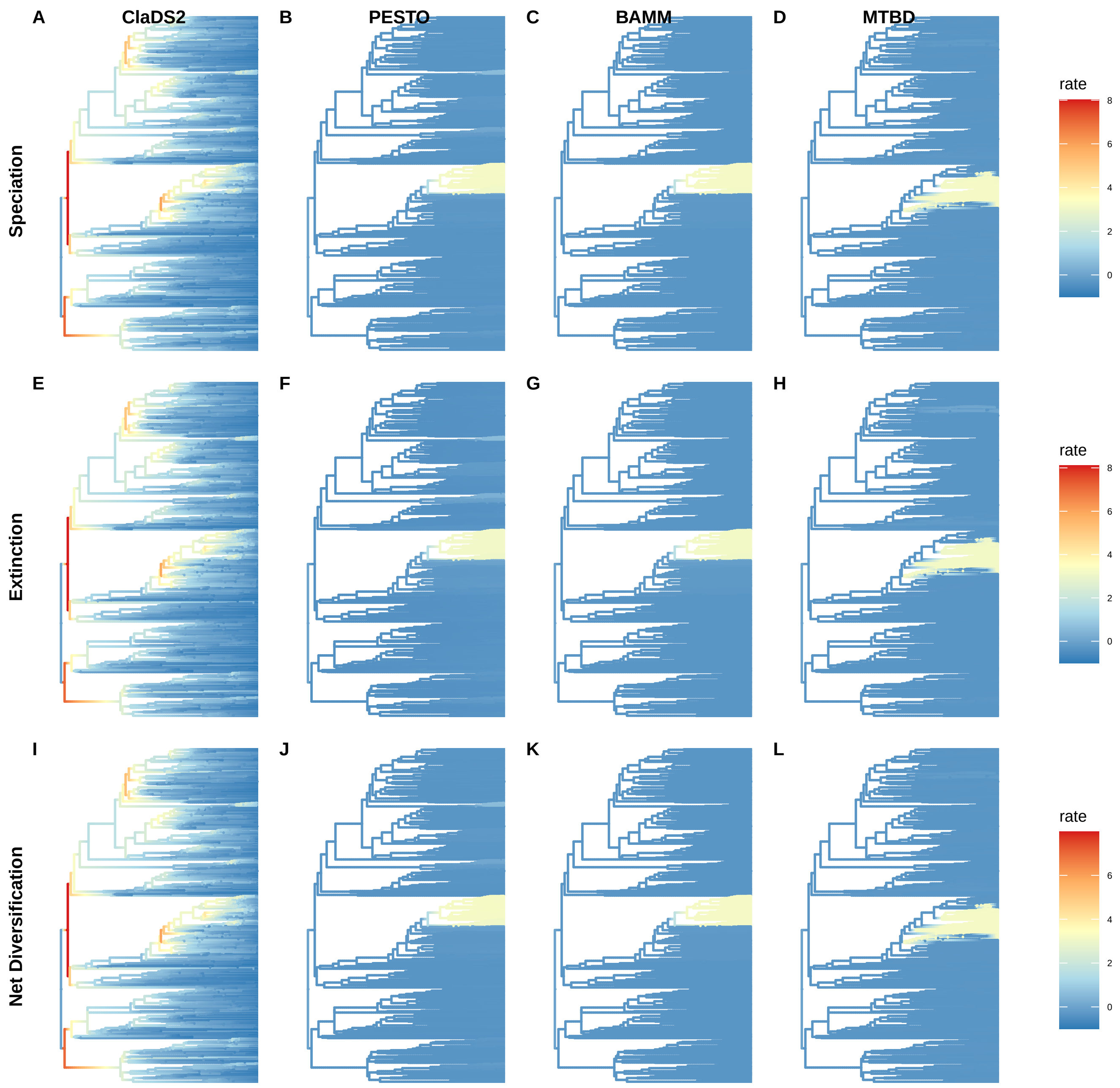
A
ClaDS2
B
PESTO
C
BAMM
D
MTBD
E
F
G
H
I
J
K
L
Speciation
Extinction
Net Diversification
rate
8
6
4
2
0

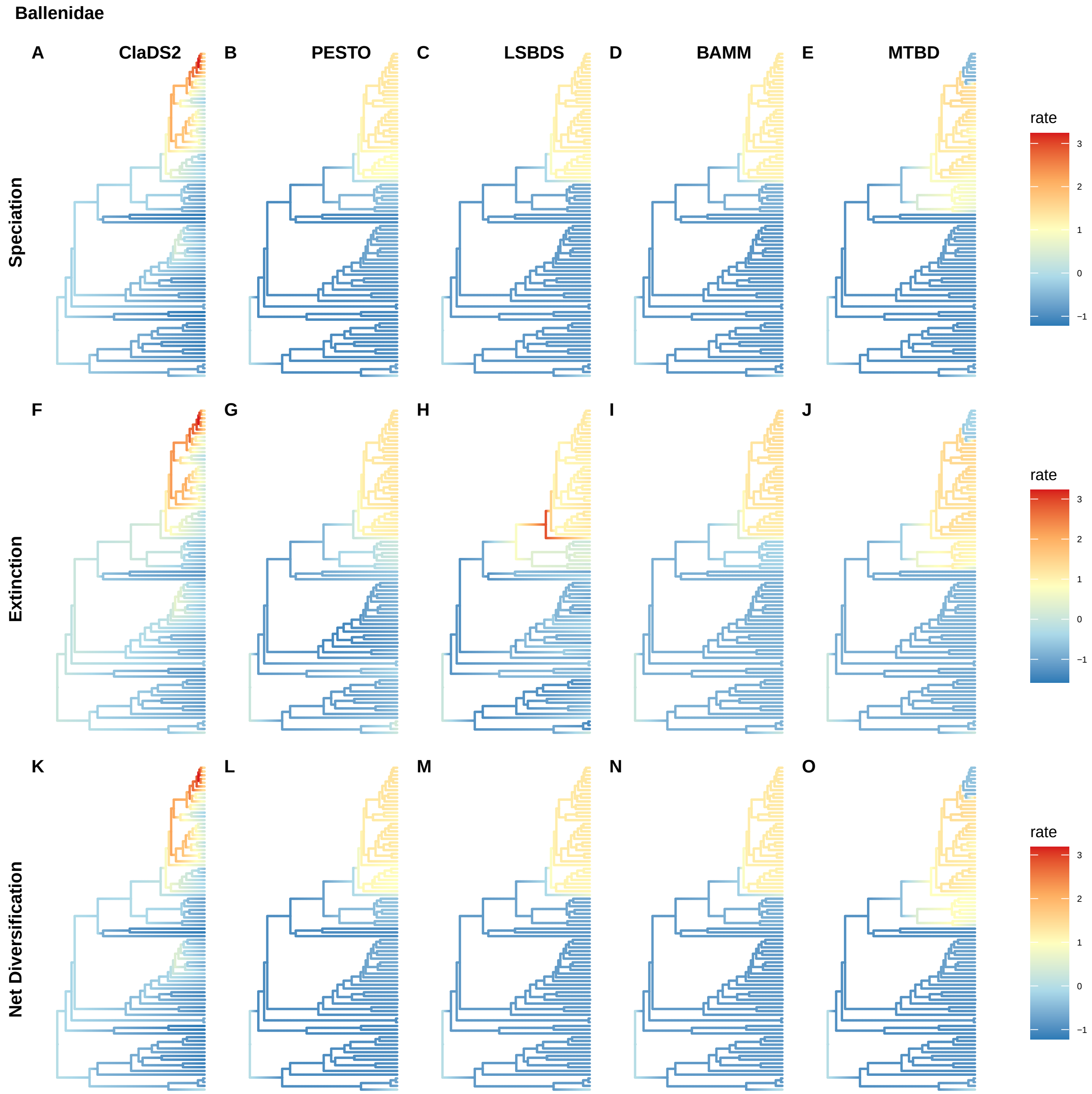
Ballenidae
A
ClaDS2
B
PESTO
C
LSBDS
D
BAMM
E
MTBD
Speciation
F
G
H
I
J
Extinction
K
L
M
N
O
Net Diversification
rate
3
2
1
0
−1
rate
3
2
1
0
−1
rate
3
2
1
0
−1

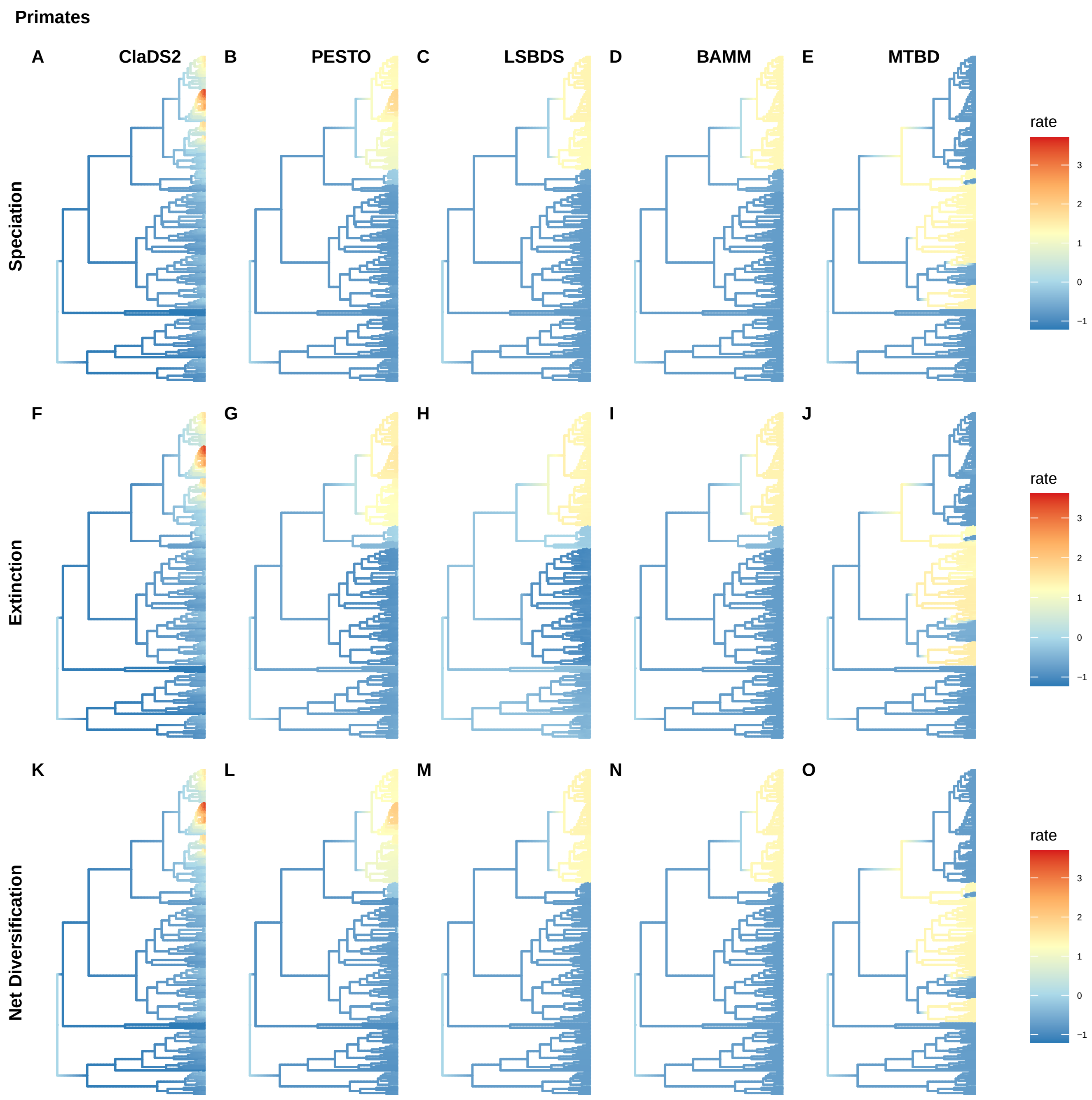
Primates
A
ClaDS2
B
PESTO
C
LSBDS
D
BAMM
E
MTBD
Speciation
F
G
H
I
J
Extinction
K
L
M
N
O
Net Diversification
rate
3
2
1
0
−1

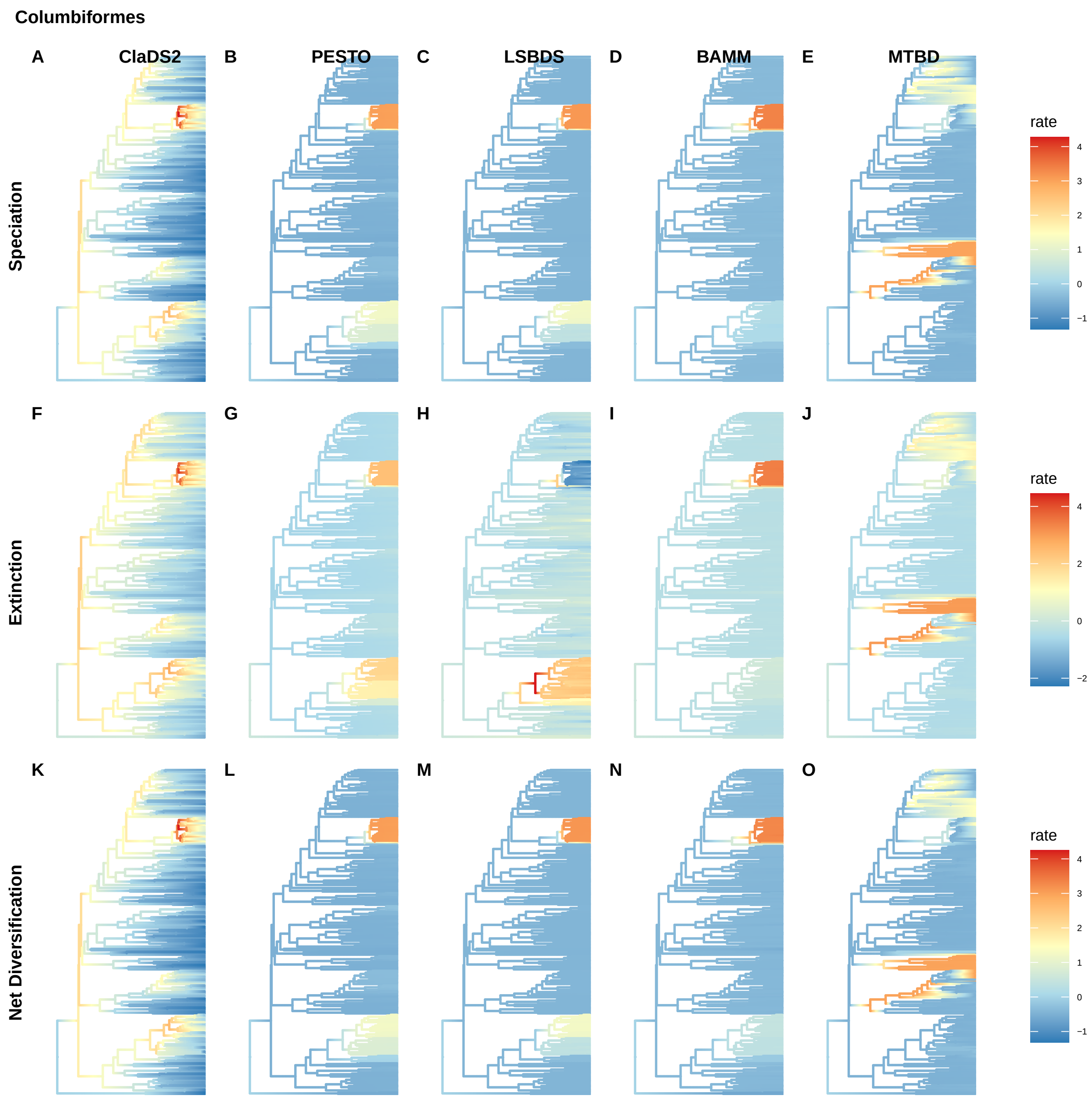
Columbiformes
A
ClaDS2
B
PESTO
C
LSBDS
D
BAMM
E
MTBD
Speciation
rate
4
3
2
1
0
−1
F
G
H
I
J
Extinction
rate
4
2
0
−2
K
L
M
N
O
Net Diversification
rate
4
3
2
1
0
−1

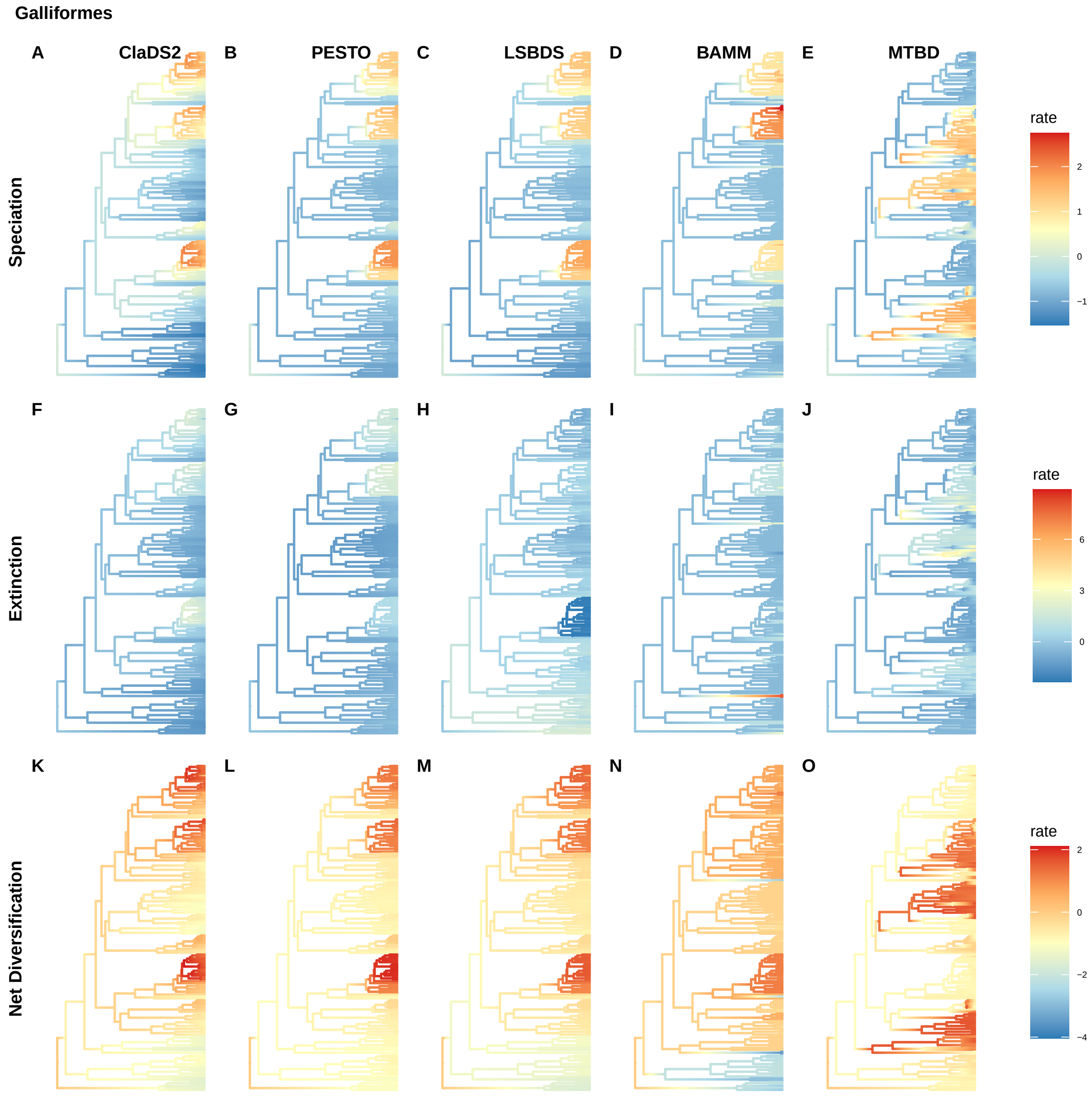
Galliformes
A
ClaDS2
B
PESTO
C
LSBDS
D
BAMM
E
MTBD
Speciation
rate
2
1
0
−1
F
G
H
I
J
Extinction
rate
6
3
0
K
L
M
N
O
Net Diversification
rate
2
0
−2
−4

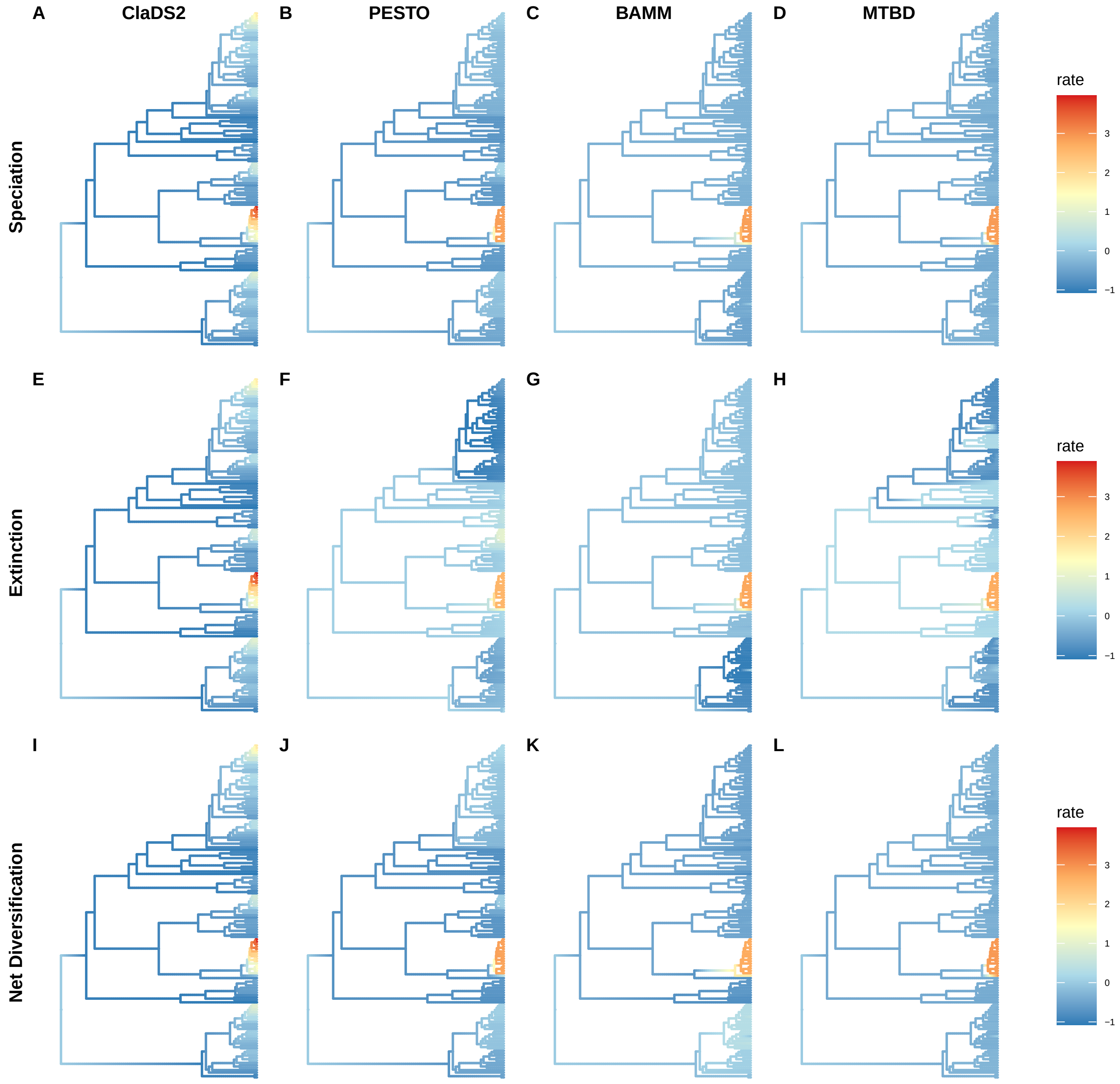
A
ClaDS2
B
PESTO
C
BAMM
D
MTBD
Speciation
rate
3
2
1
0
−1
E
F
G
H
Extinction
rate
3
2
1
0
−1
I
J
K
L
Net Diversification
rate
3
2
1
0
−1

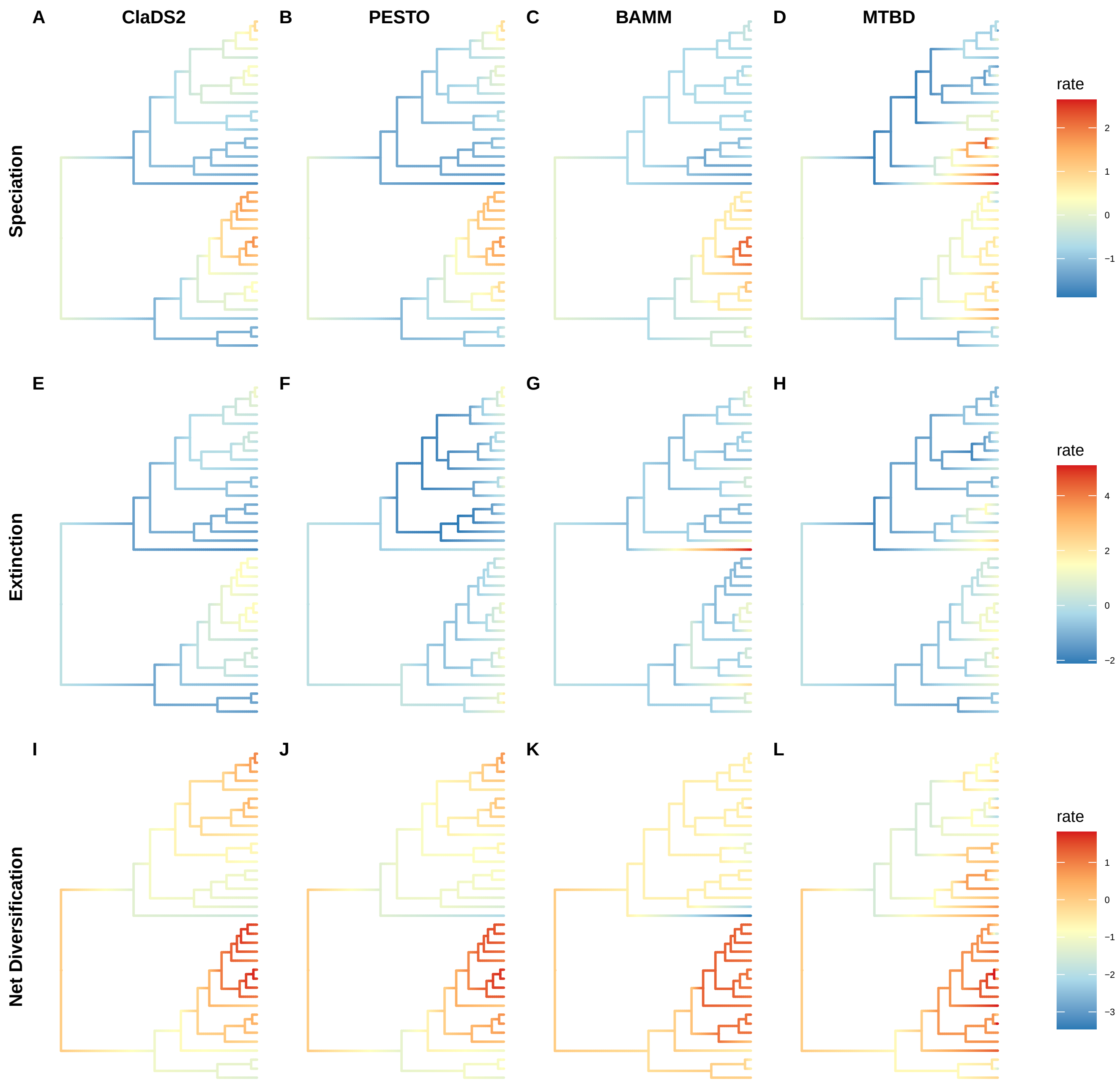
A
ClaDS2
B
PESTO
C
BAMM
D
MTBD
Speciation
rate
2
1
0
−1
E
F
G
H
Extinction
rate
4
2
0
−2
I
J
K
L
Net Diversification
rate
1
0
−1
−2
−3

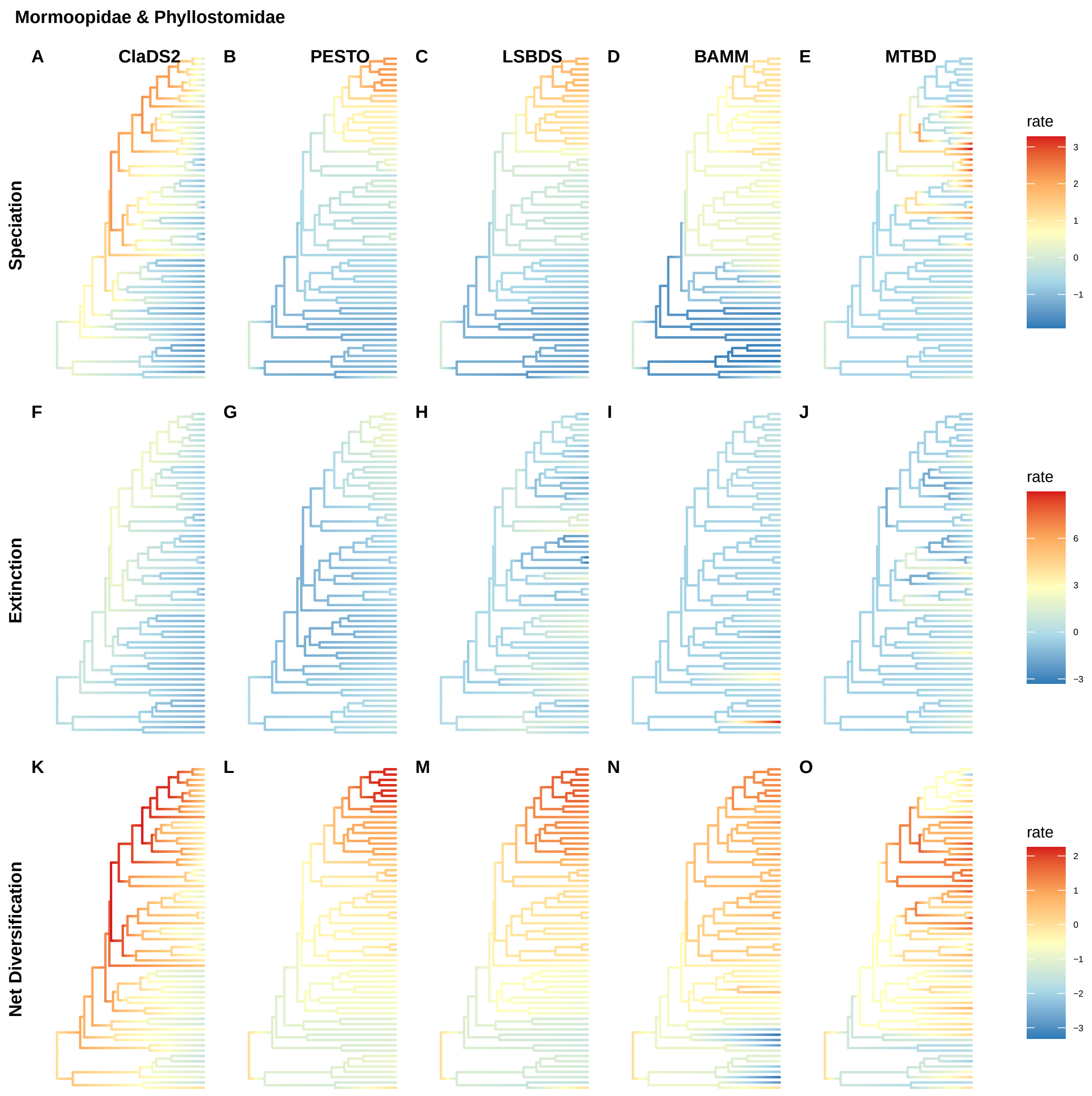
Mormoopidae & Phyllostomidae
A
ClaDS2
B
PESTO
C
LSBDS
D
BAMM
E
MTBD
Speciation
rate
3
2
1
0
−1
F
G
H
I
J
Extinction
rate
6
3
0
−3
K
L
M
N
O
Net Diversification
rate
2
1
0
−1
−2
−3

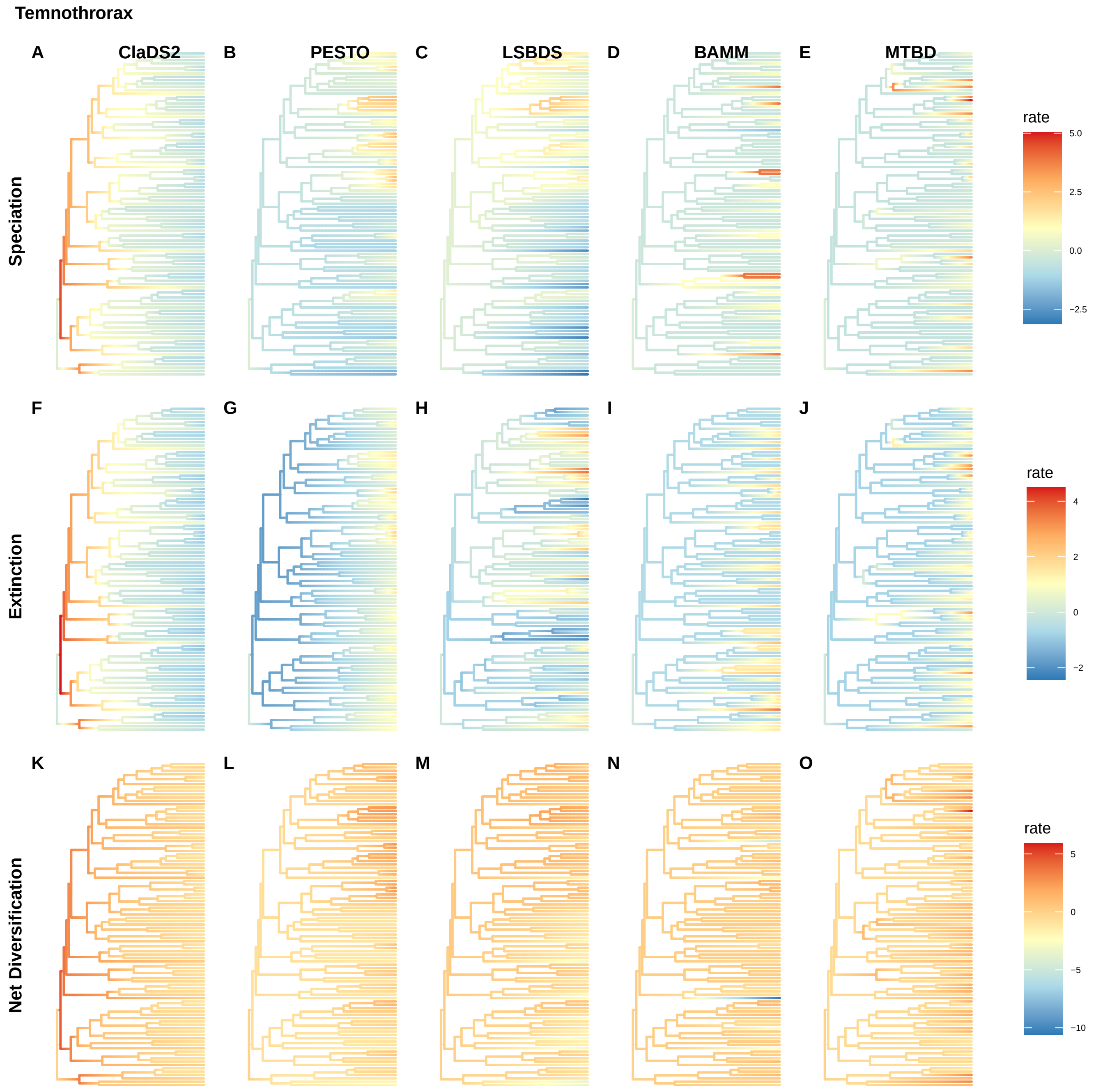
Temnothrorax
A
ClaDS2
B
PESTO
C
LSBDS
D
BAMM
E
MTBD
Speciation
rate
5.0
2.5
0.0
−2.5
F
G
H
I
J
Extinction
rate
4
2
0
−2
K
L
M
N
O
Net Diversification
rate
5
0
−5
−10

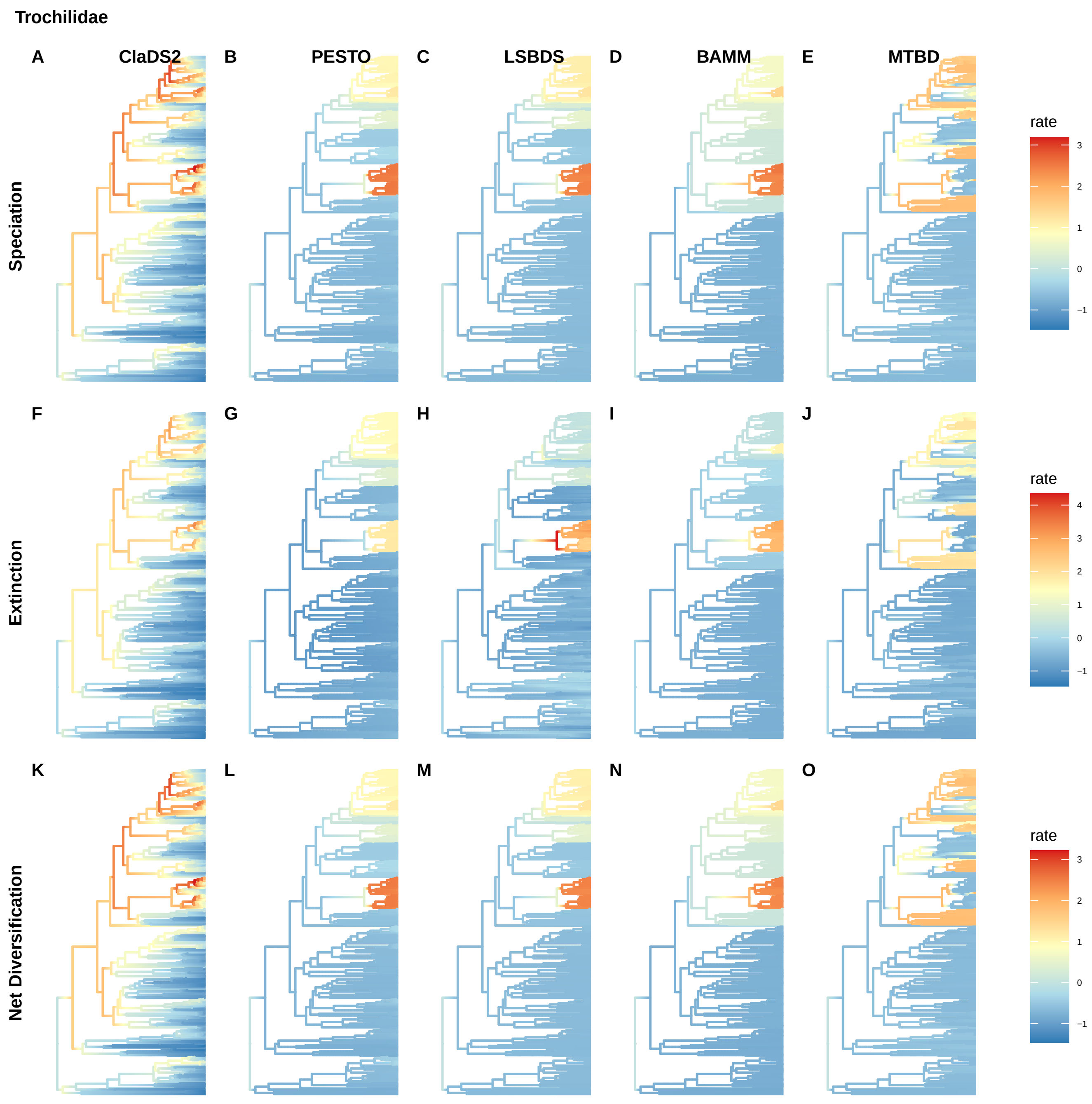
Trochilidae
A
ClaDS2
B
PESTO
C
LSBDS
D
BAMM
E
MTBD
Speciation
rate
3
2
1
0
−1
F
G
H
I
J
Extinction
rate
4
3
2
1
0
−1
K
L
M
N
O
Net Diversification
rate
3
2
1
0
−1

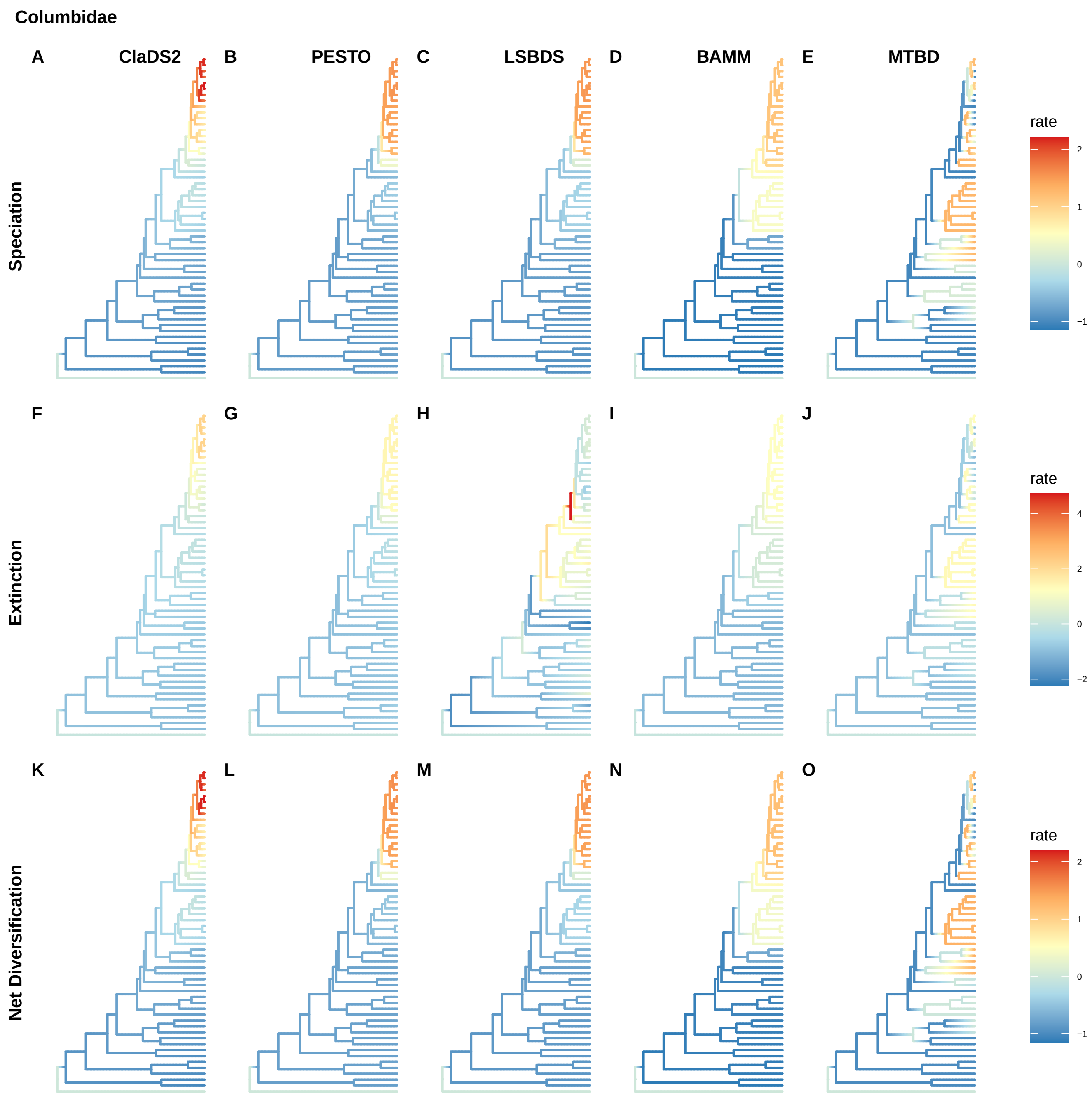
Columbidae
A
ClaDS2
B
PESTO
C
LSBDS
D
BAMM
E
MTBD
F
G
H
I
J
K
L
M
N
O
Speciation
Extinction
Net Diversification
rate
2
1
0
−1
rate
4
2
0
−2
rate
2
1
0
−1

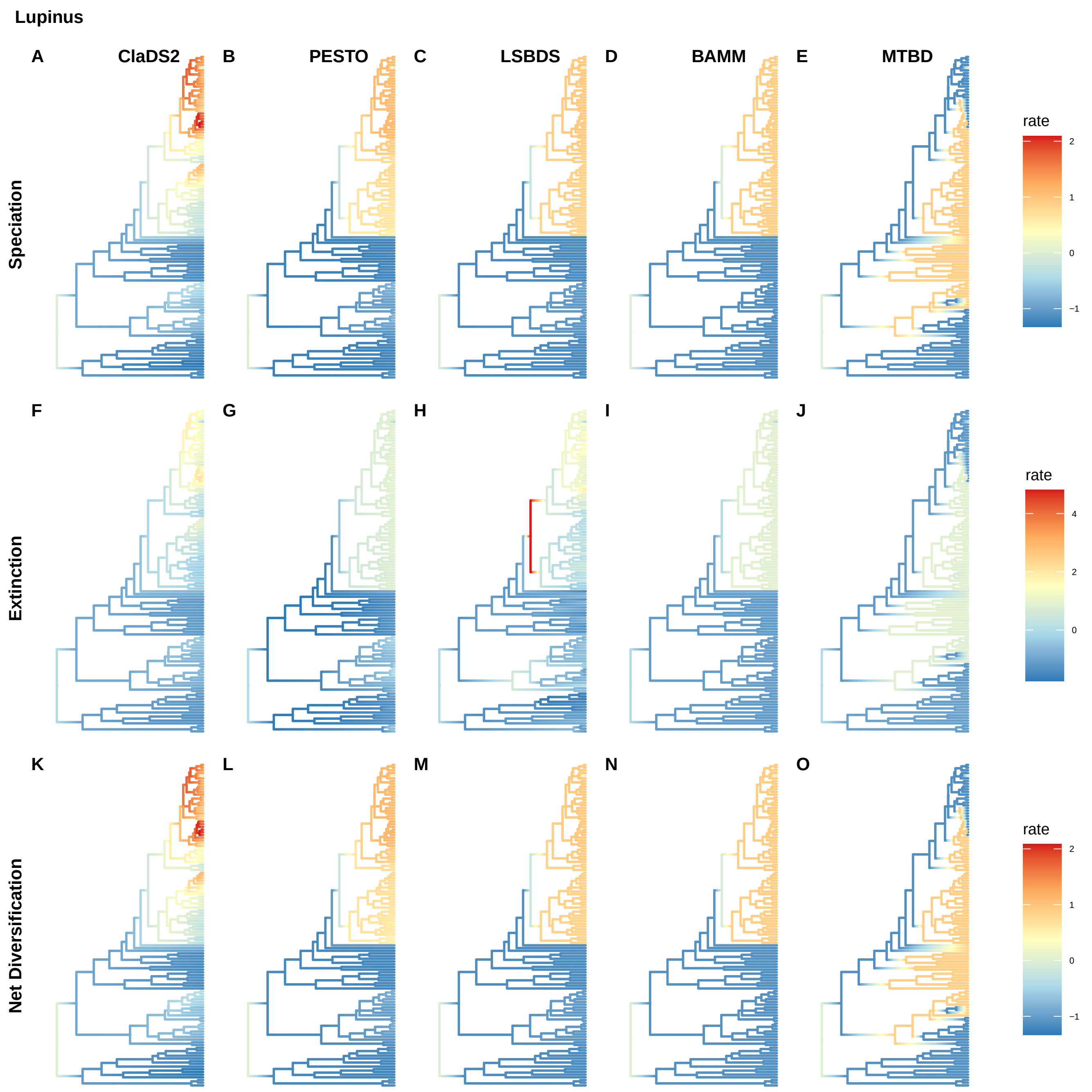
Lupinus
A
ClaDS2
B
PESTO
C
LSBDS
D
BAMM
E
MTBD
Speciation
rate
2
1
0
−1
F
G
H
I
J
Extinction
rate
4
2
0
K
L
M
N
O
Net Diversification
rate
2
1
0
−1

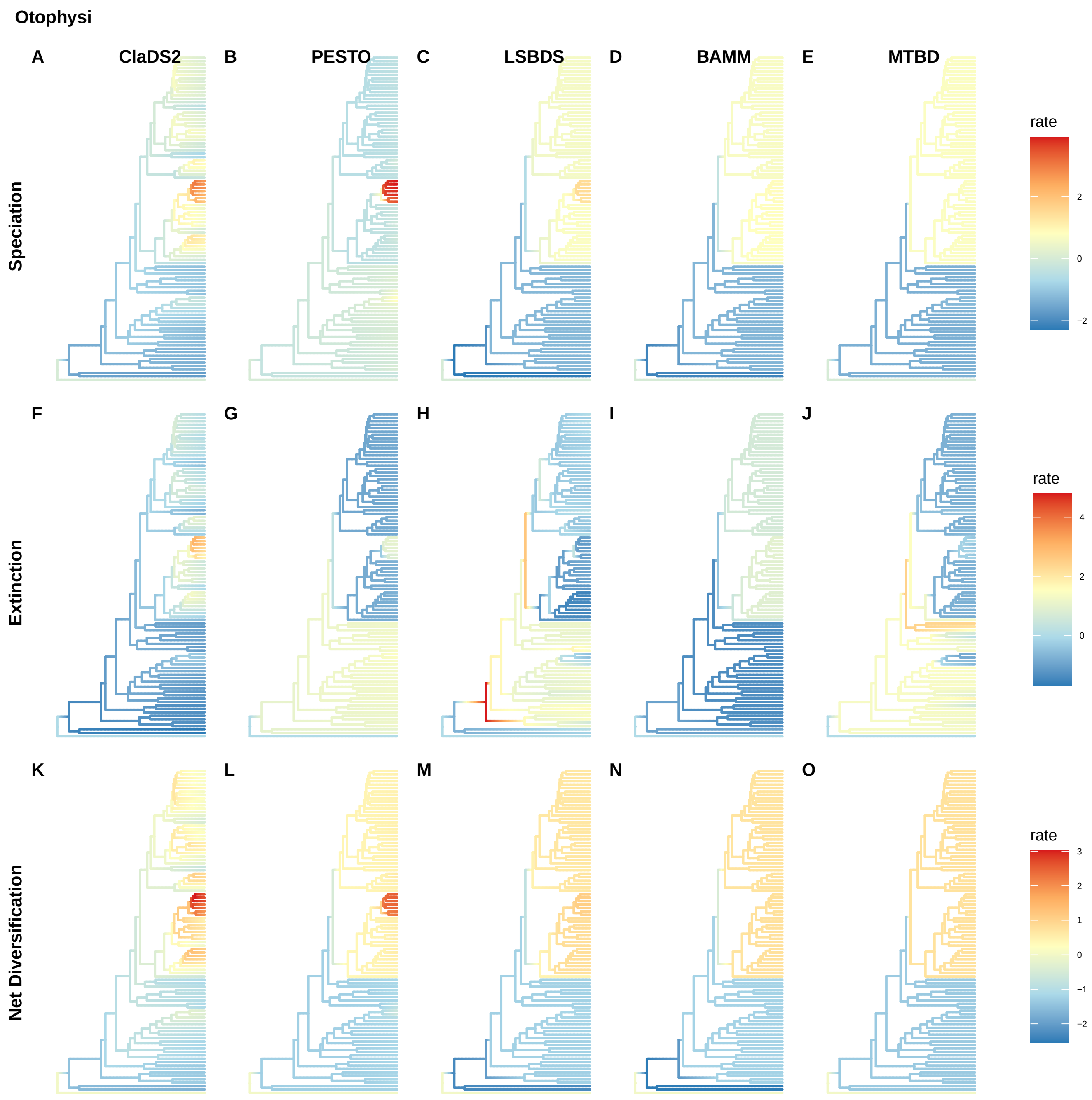

Otophysi
A
ClaDS2
B
PESTO
C
LSBDS
D
BAMM
E
MTBD
Speciation
rate
2
0
−2
F
G
H
I
J
Extinction
rate
4
2
0
K
L
M
N
O
Net Diversification
rate
3
2
1
0
−1
−2

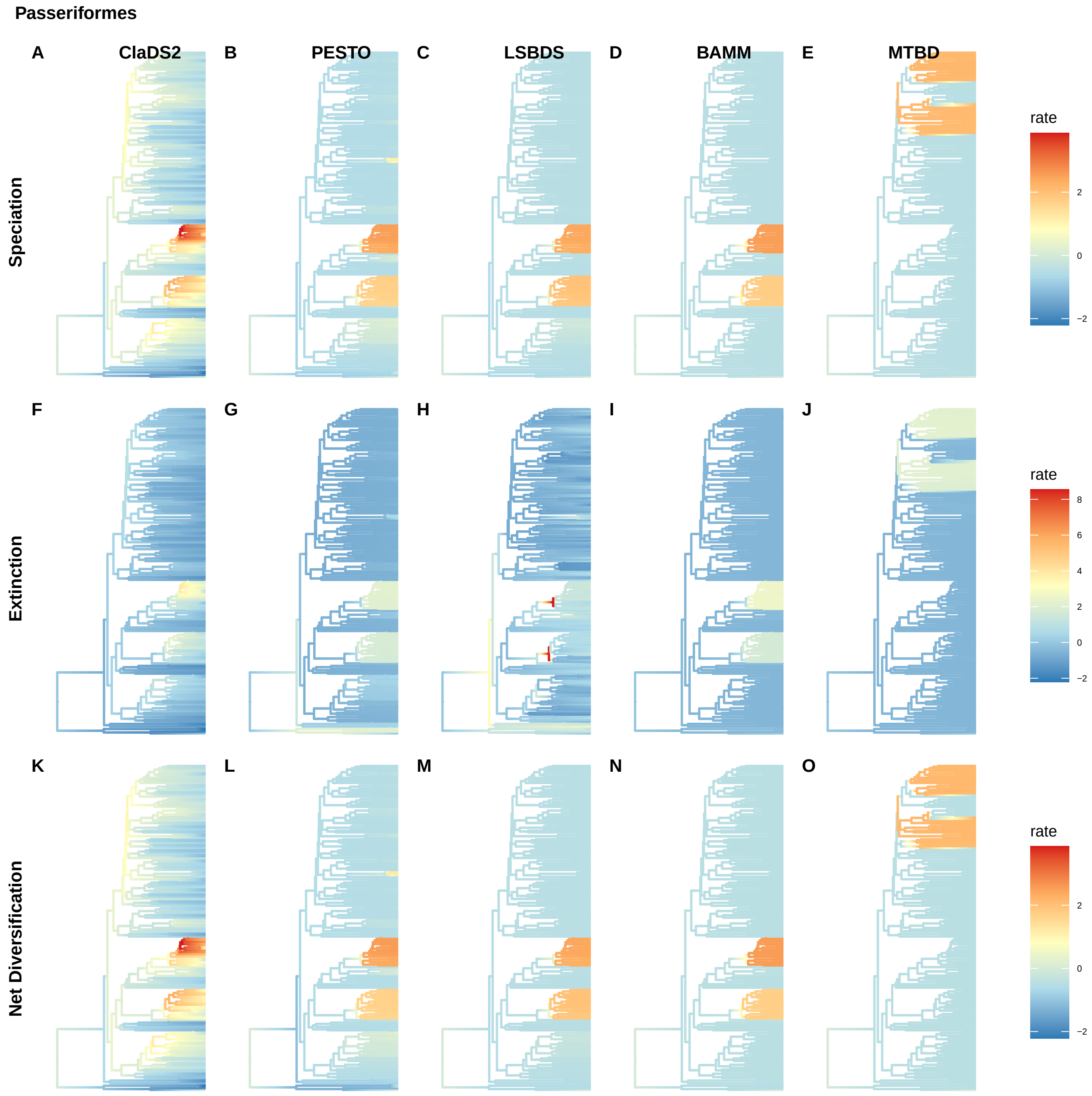
Passeriformes
A
ClaDS2
B
PESTO
C
LSBDS
D
BAMM
E
MTBD
Speciation
rate
2
0
−2
F
G
H
I
J
Extinction
rate
8
6
4
2
0
−2
K
L
M
N
O
Net Diversification
rate
2
0
−2

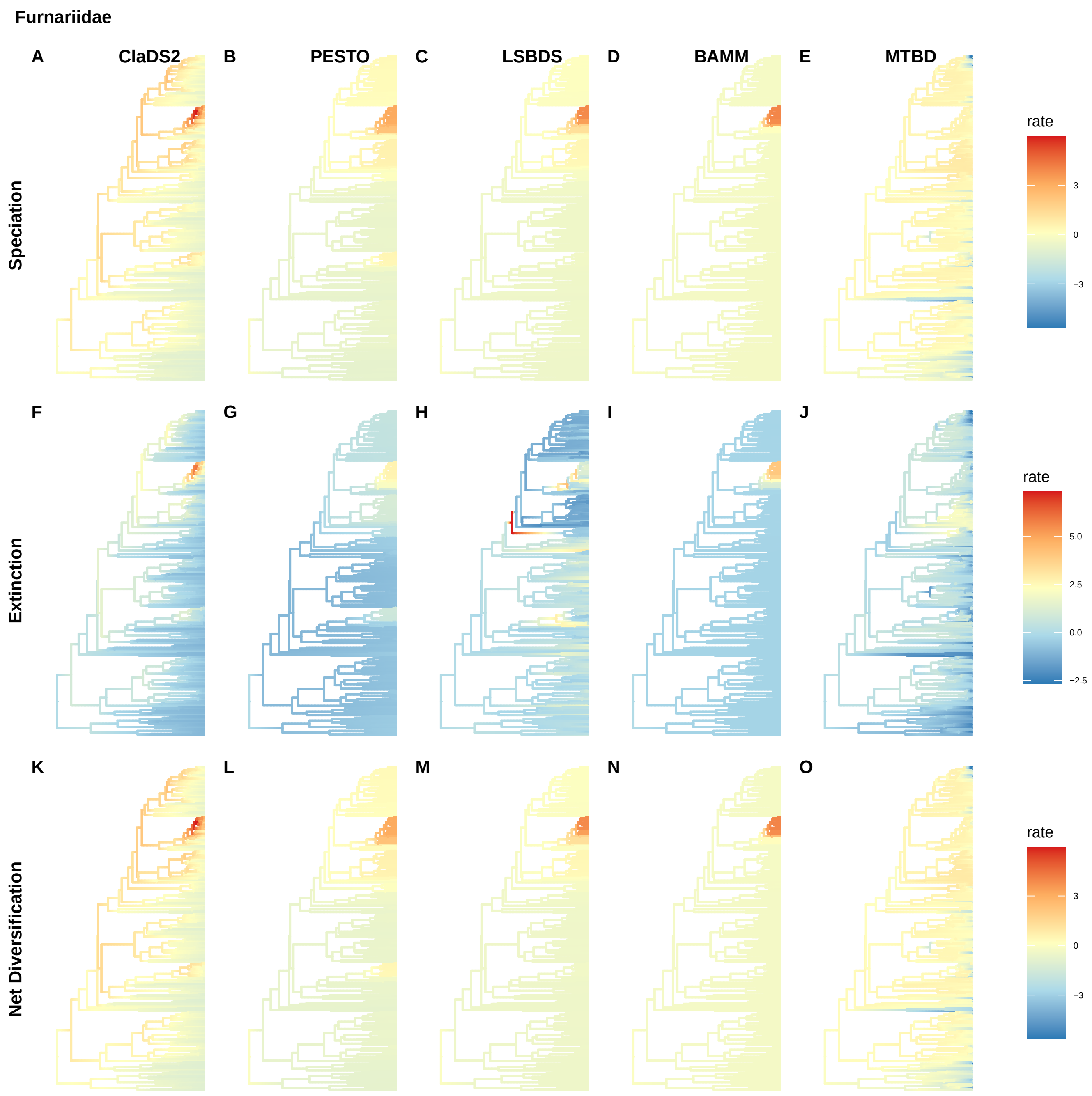
Furnariidae
A
ClaDS2
B
PESTO
C
LSBDS
D
BAMM
E
MTBD
Speciation
rate
3
0
−3
F
G
H
I
J
Extinction
rate
5.0
2.5
0.0
−2.5
K
L
M
N
O
Net Diversification
rate
3
0
−3

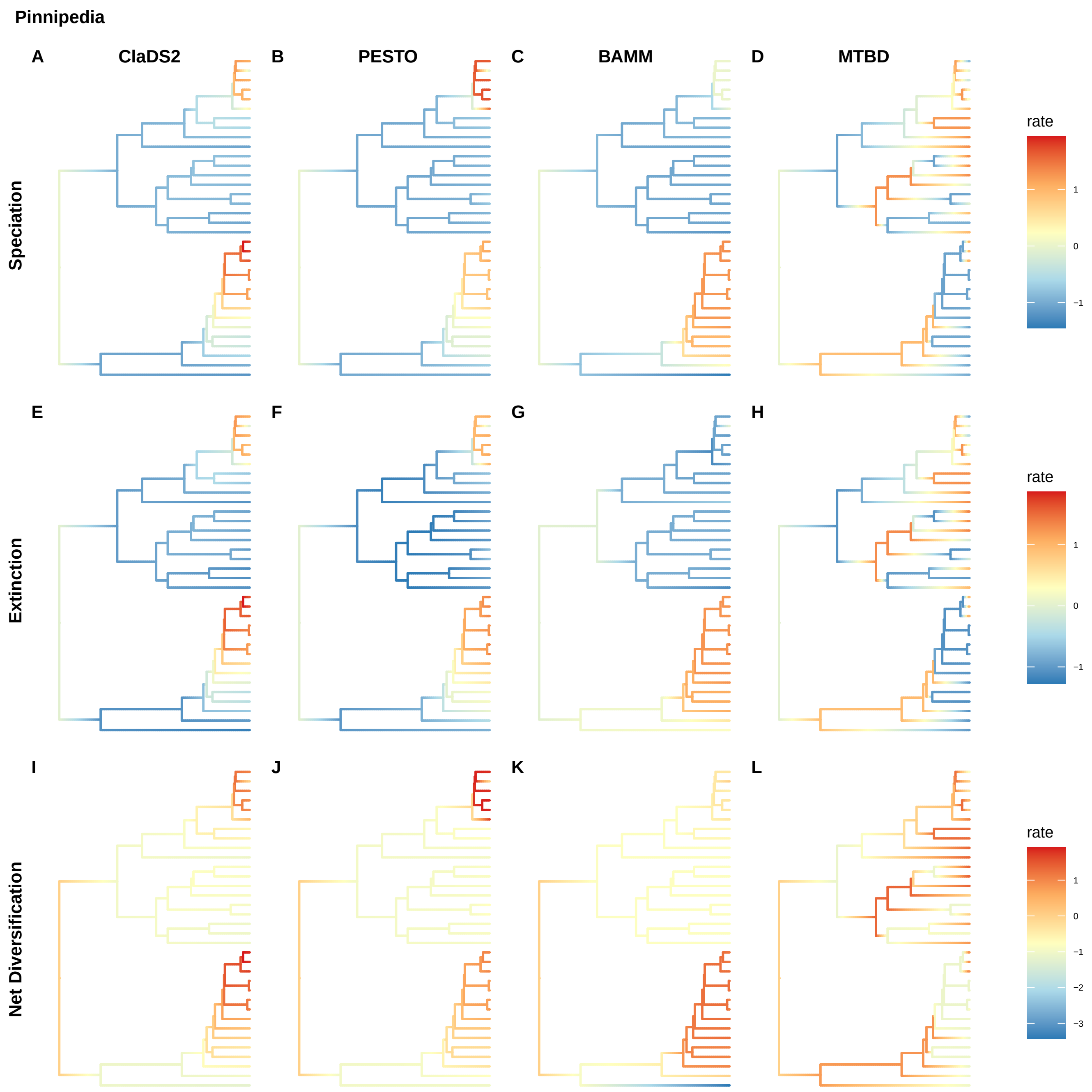
Pinnipedia
A
ClaDS2
B
PESTO
C
BAMM
D
MTBD
Speciation
rate
1
0
−1
E
F
G
H
Extinction
rate
1
0
−1
I
J
K
L
Net Diversification
rate
1
0
−1
−2
−3

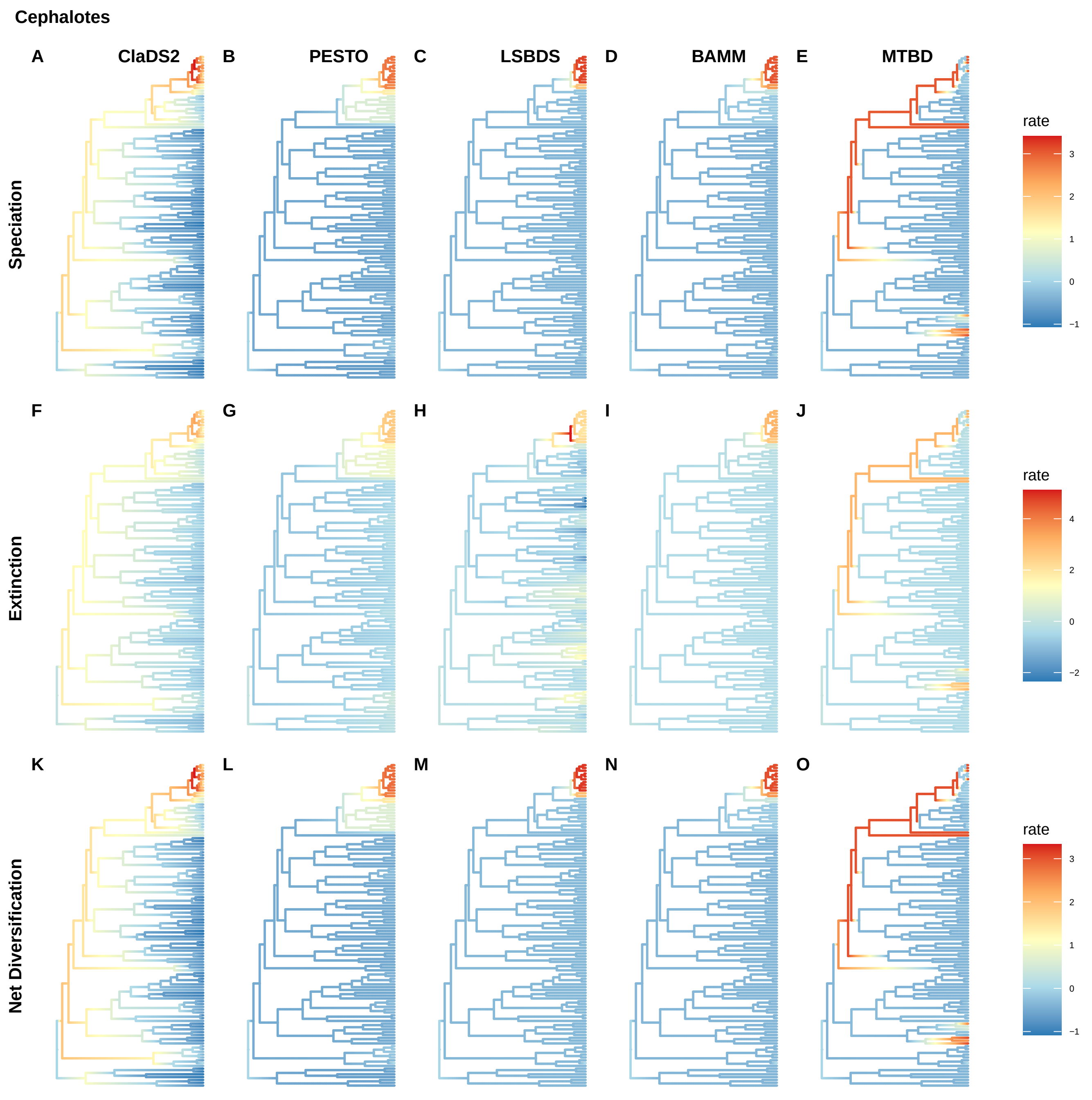
Cephalotes
A
ClaDS2
B
PESTO
C
LSBDS
D
BAMM
E
MTBD
Speciation
rate
3
2
1
0
−1
F
G
H
I
J
Extinction
rate
4
2
0
−2
K
L
M
N
O
Net Diversification
rate
3
2
1
0
−1

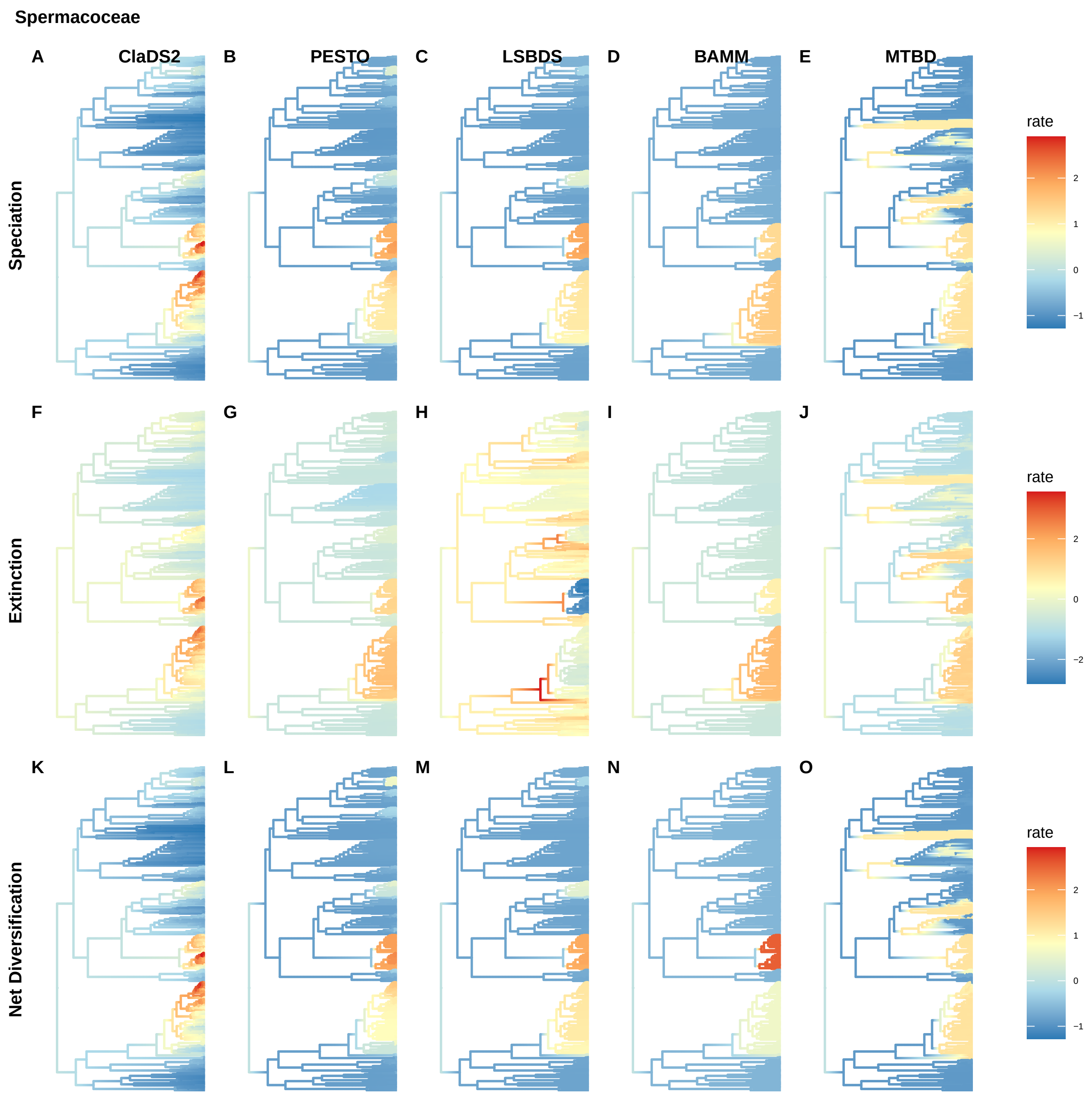
Spermacoceae
A
ClaDS2
B
PESTO
C
LSBDS
D
BAMM
E
MTBD
Speciation
rate
2
1
0
−1
F
G
H
I
J
Extinction
rate
2
0
−2
K
L
M
N
O
Net Diversification
rate
2
1
0
−1

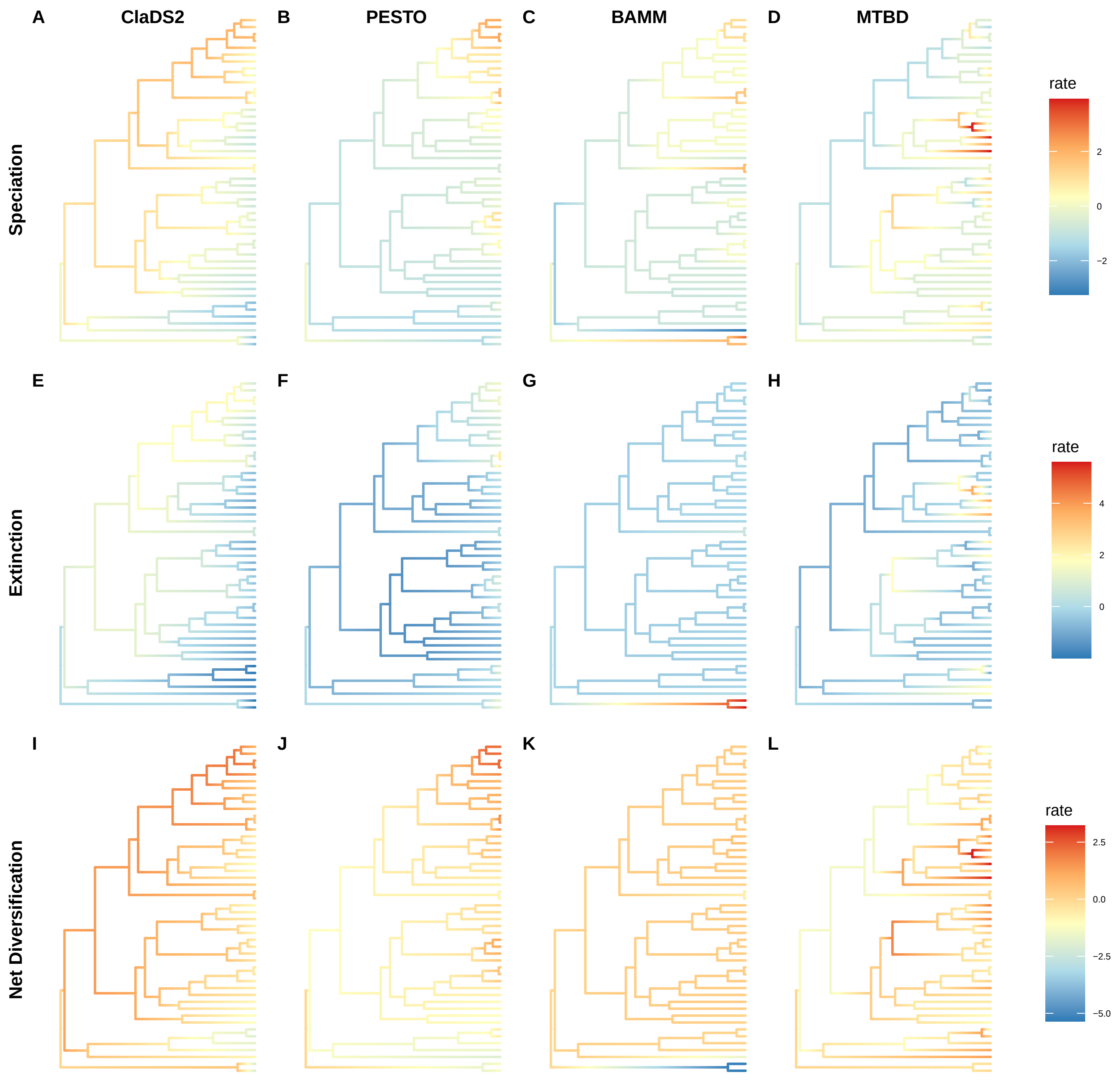
A
ClaDS2
B
PESTO
C
BAMM
D
MTBD
Speciation
rate
2
0
−2
E
F
G
H
Extinction
rate
4
2
0
I
J
K
L
Net Diversification
rate
2.5
0.0
−2.5
−5.0

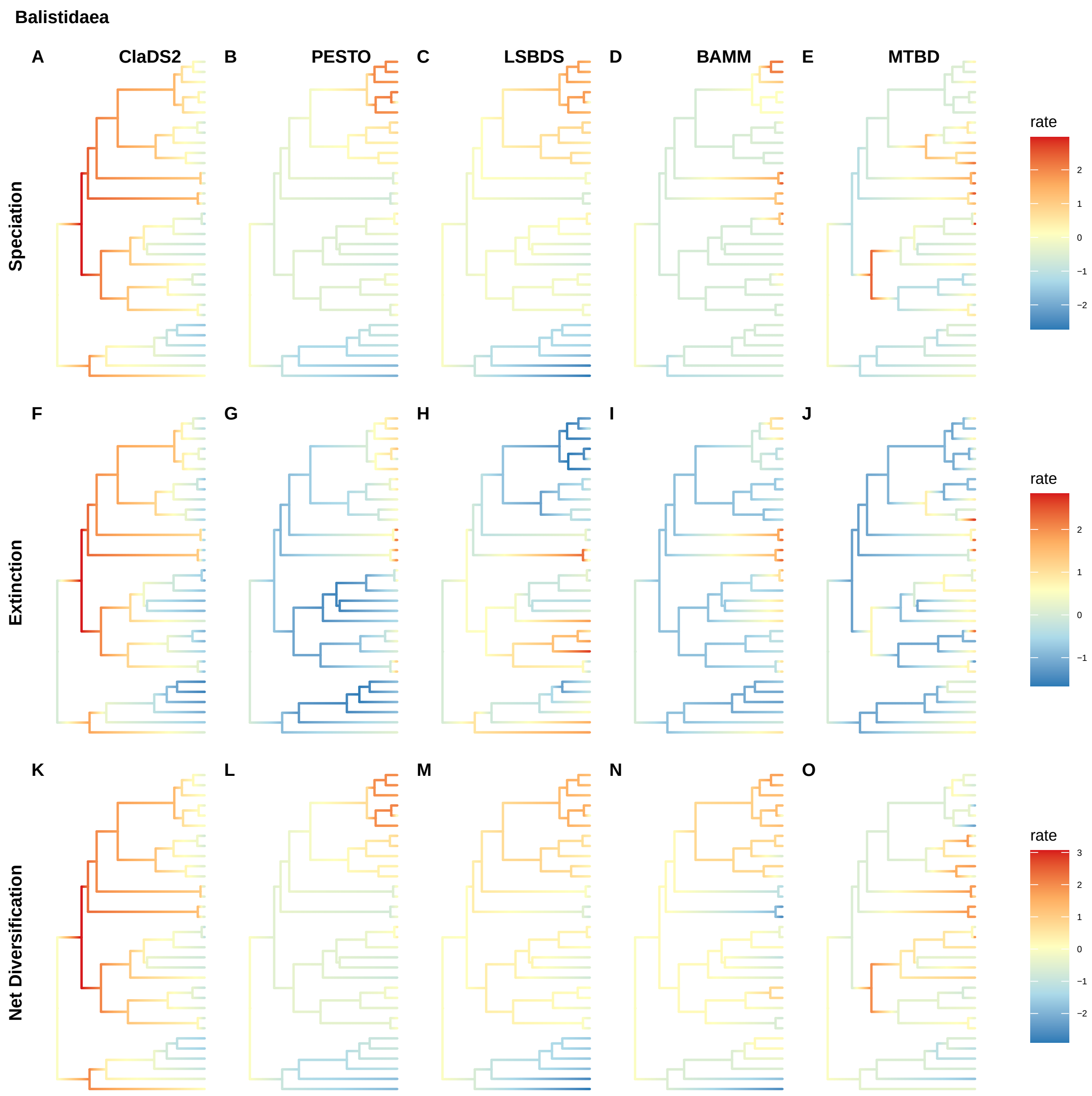
Balistidaea
A
ClaDS2
B
PESTO
C
LSBDS
D
BAMM
E
MTBD
Speciation
rate
2
1
0
−1
−2
F
G
H
I
J
Extinction
rate
2
1
0
−1
K
L
M
N
O
Net Diversification
rate
3
2
1
0
−1
−2

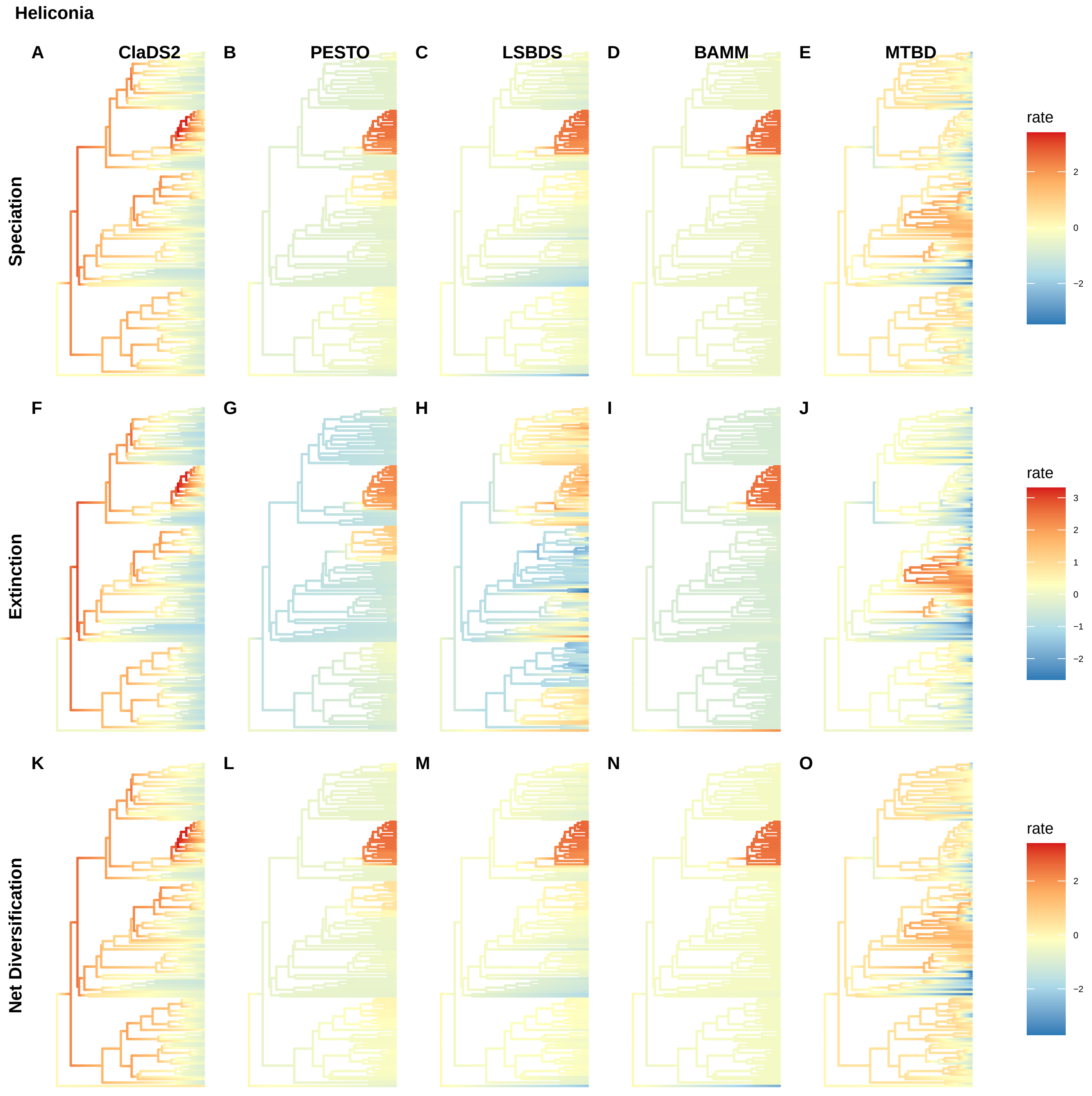

Heliconia
A
ClaDS2
B
PESTO
C
LSBDS
D
BAMM
E
MTBD
Speciation
rate
2
0
−2
F
G
H
I
J
Extinction
rate
3
2
1
0
−1
−2
K
L
M
N
O
Net Diversification
rate
2
0
−2

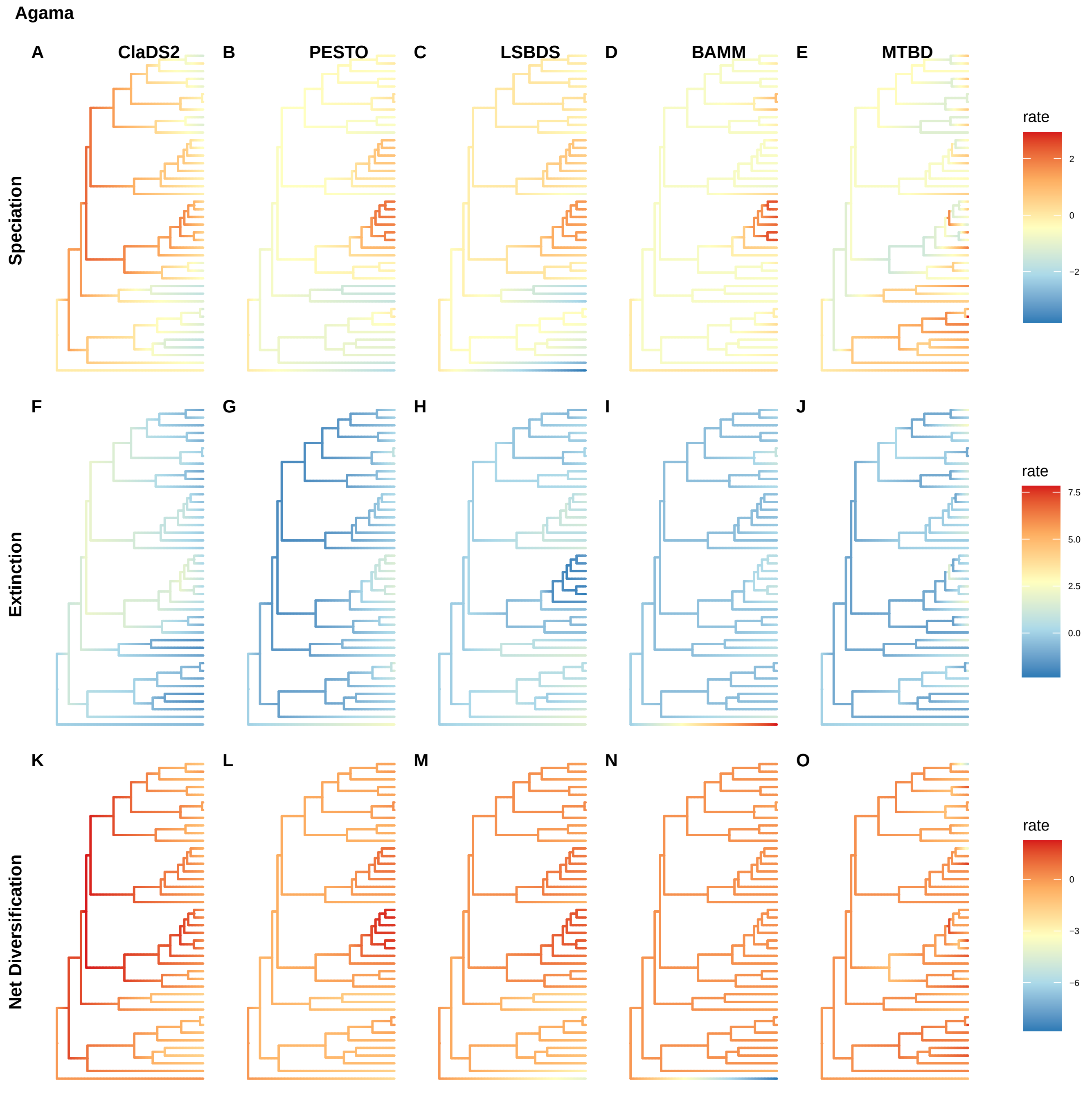

Agama
A
ClaDS2
B
PESTO
C
LSBDS
D
BAMM
E
MTBD
Speciation
rate
2
0
−2
F
G
H
I
J
Extinction
rate
7.5
5.0
2.5
0.0
K
L
M
N
O
Net Diversification
rate
0
−3
−6

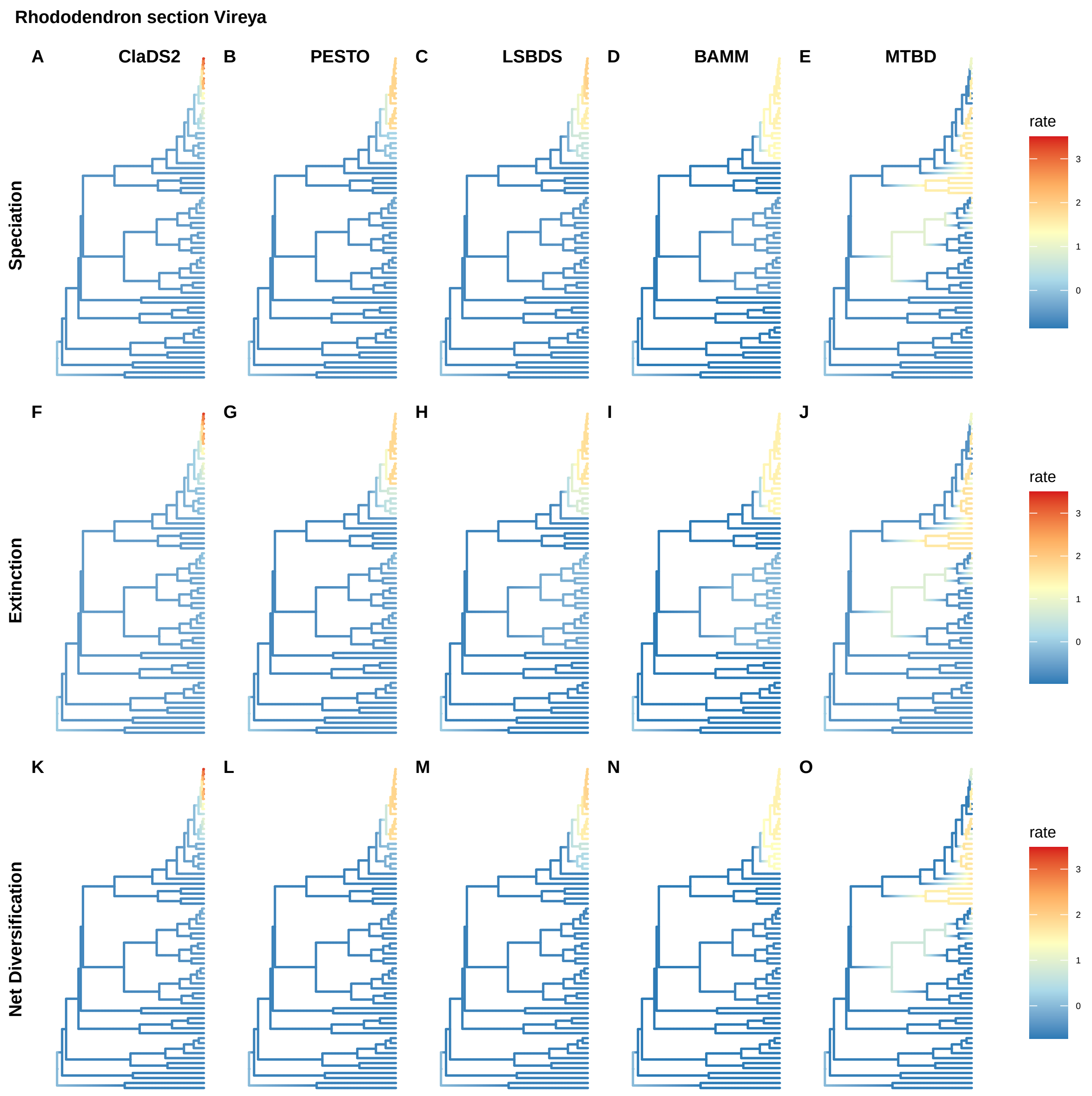
Rhododendron section Vireya
A
ClaDS2
B
PESTO
C
LSBDS
D
BAMM
E
MTBD
F
G
H
I
J
K
L
M
N
O
Speciation
Extinction
Net Diversification
rate
3
2
1
0
rate
3
2
1
0
rate
3
2
1
0

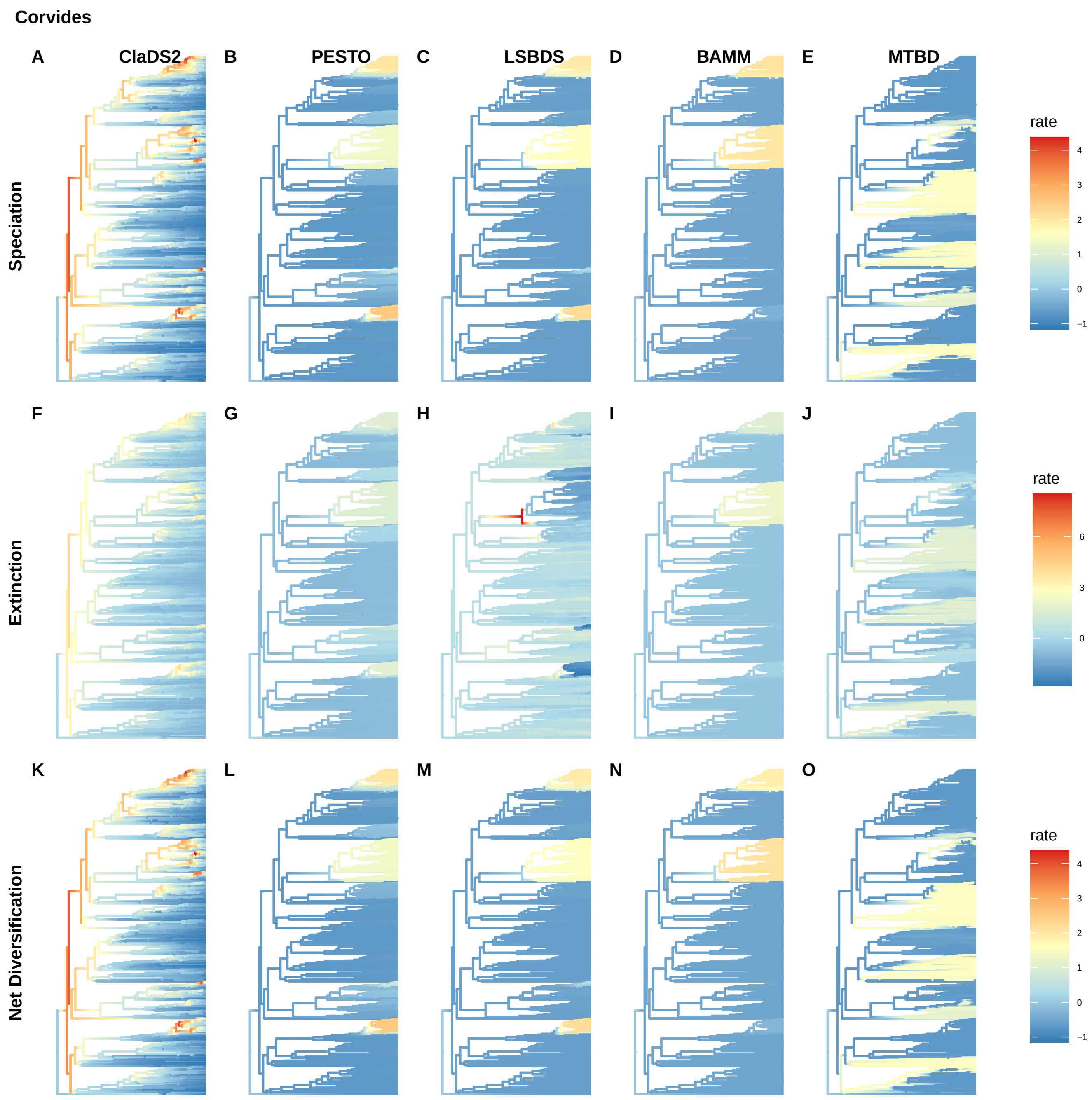
Corvides
A
ClaDS2
B
PESTO
C
LSBDS
D
BAMM
E
MTBD
Speciation
rate
4
3
2
1
0
−1
F
G
H
I
J
Extinction
rate
6
3
0
K
L
M
N
O
Net Diversification
rate
4
3
2
1
0
−1

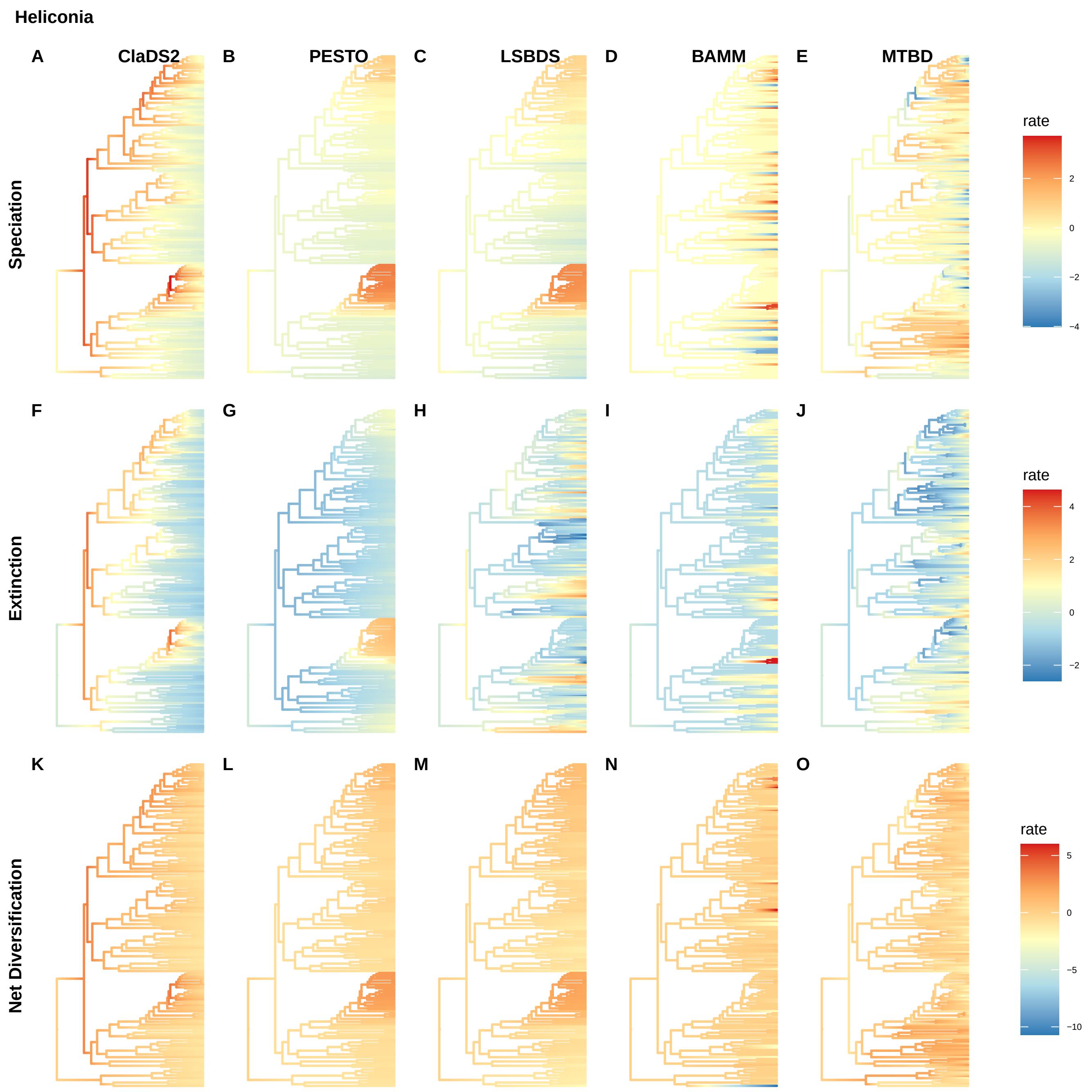
Heliconia
A
ClaDS2
B
PESTO
C
LSBDS
D
BAMM
E
MTBD
Speciation
rate
2
0
−2
−4
F
G
H
I
J
Extinction
rate
4
2
0
−2
K
L
M
N
O
Net Diversification
rate
5
0
−5
−10

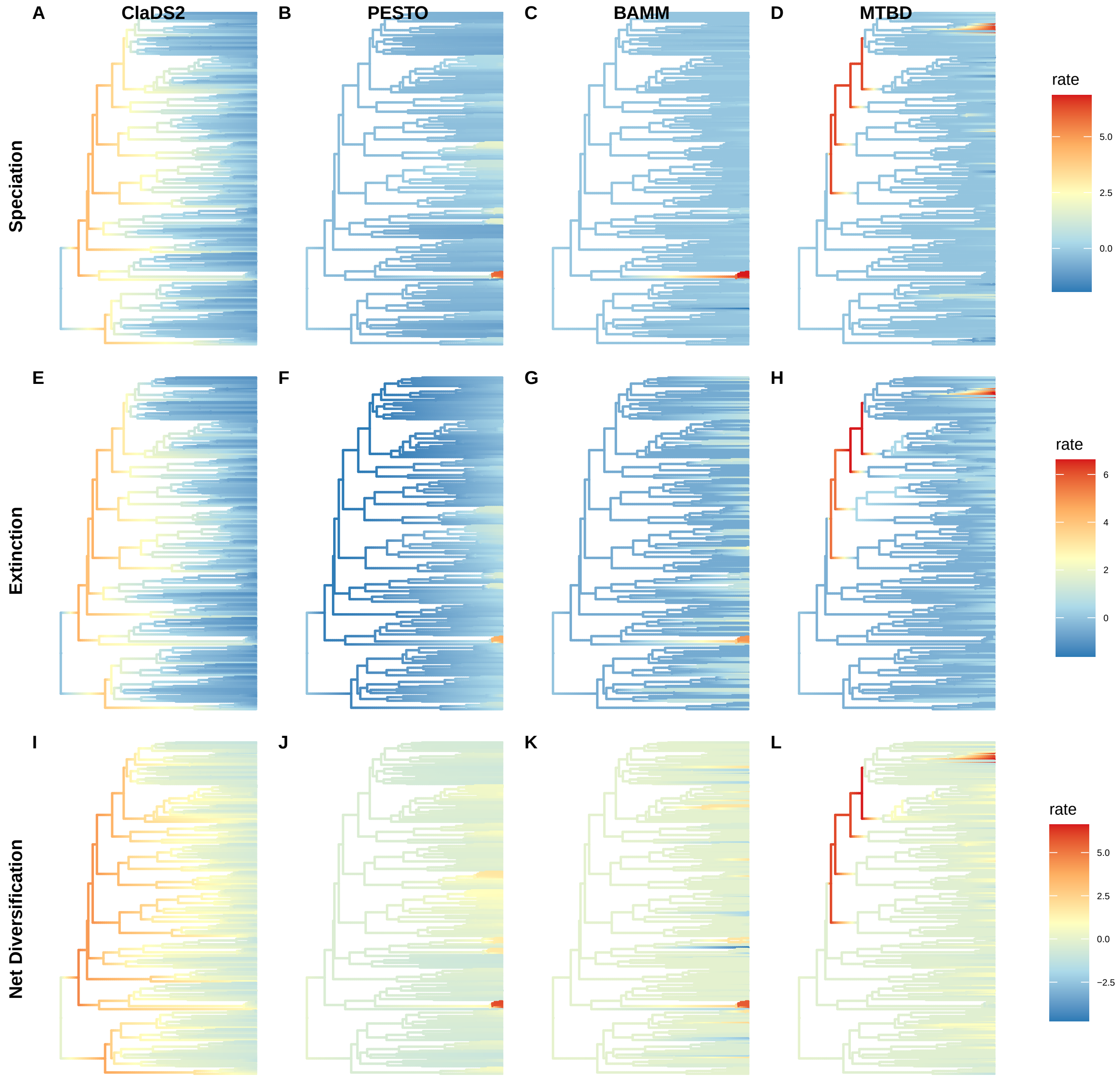
A
ClaDS2
B
PESTO
C
BAMM
D
MTBD
Speciation
rate
5.0
2.5
0.0
E
F
G
H
Extinction
rate
6
4
2
0
I
J
K
L
Net Diversification
rate
5.0
2.5
0.0
−2.5

Sebastes

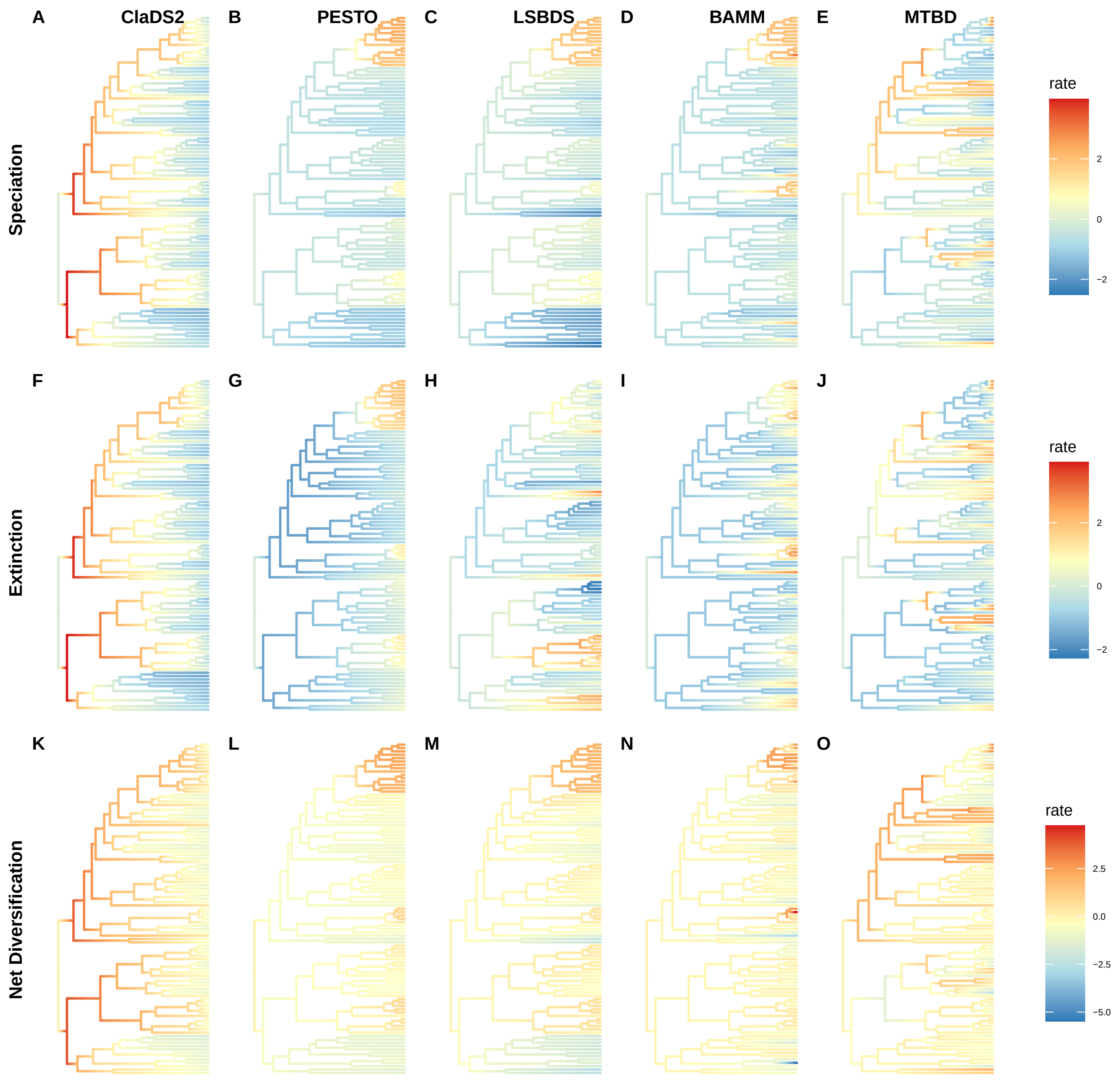
A
ClaDS2
B
PESTO
C
LSBDS
D
BAMM
E
MTBD
Speciation
rate
2
0
−2
F
G
H
I
J
Extinction
rate
2
0
−2
K
L
M
N
O
Net Diversification
rate
2.5
0.0
−2.5
−5.0

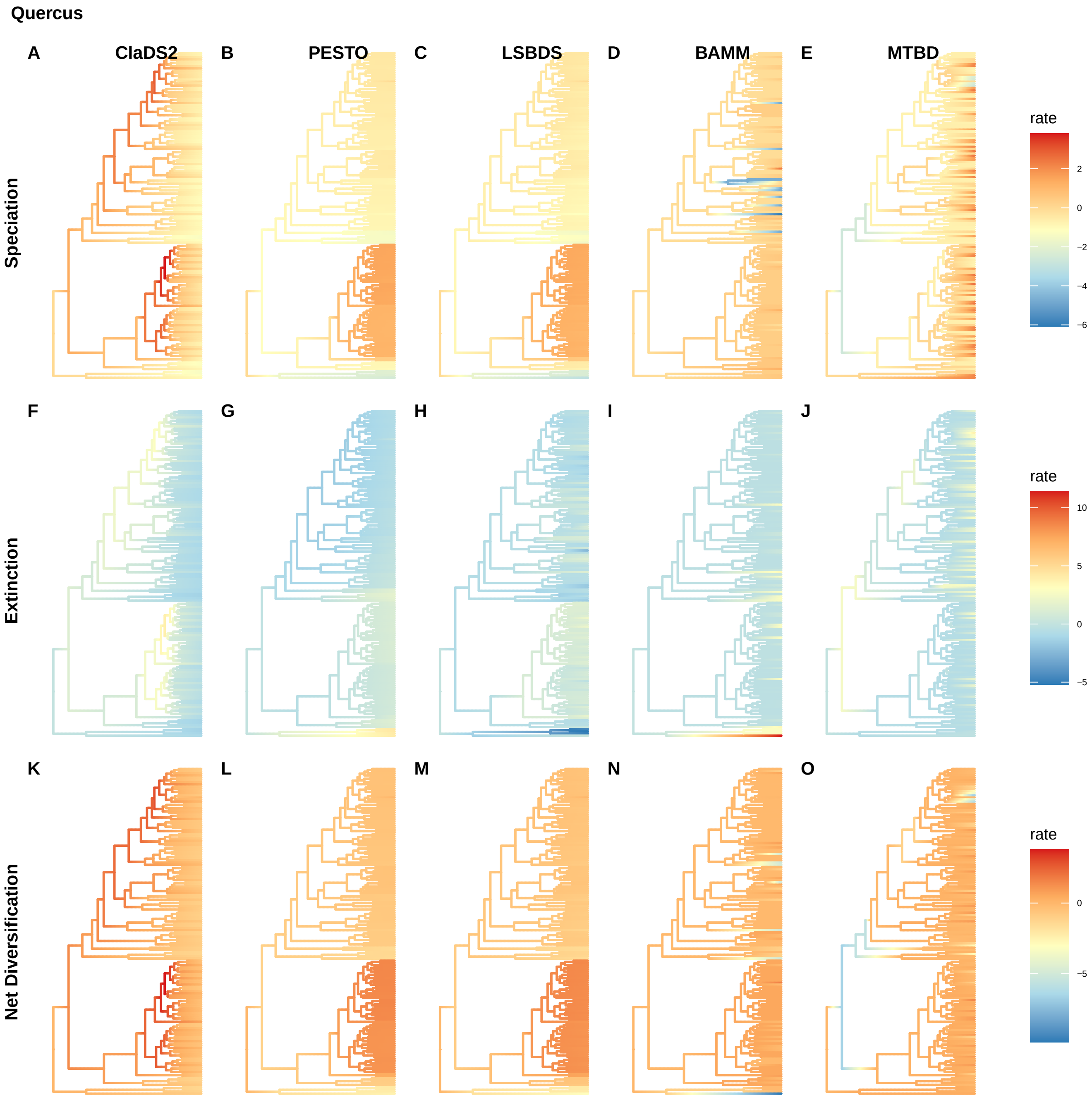
Quercus
A
ClaDS2
B
PESTO
C
LSBDS
D
BAMM
E
MTBD
Speciation
rate
2
0
−2
−4
−6
F
G
H
I
J
Extinction
rate
10
5
0
−5
K
L
M
N
O
Net Diversification
rate
0
−5

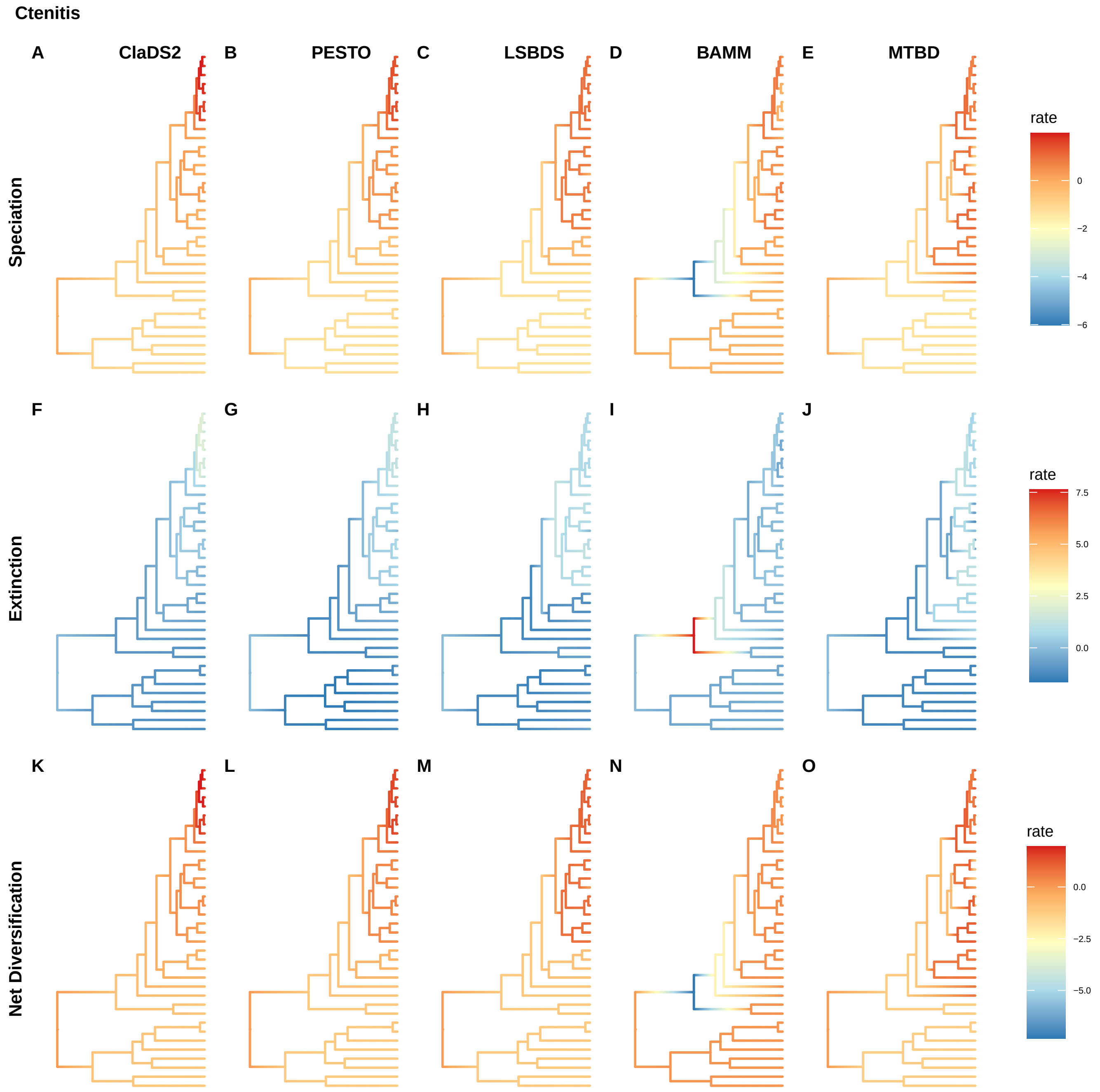

Ctenitis
A
ClaDS2
B
PESTO
C
LSBDS
D
BAMM
E
MTBD
Speciation
rate
0
−2
−4
−6
F
G
H
I
J
Extinction
rate
7.5
5.0
2.5
0.0
K
L
M
N
O
Net Diversification
rate
0.0
−2.5
−5.0

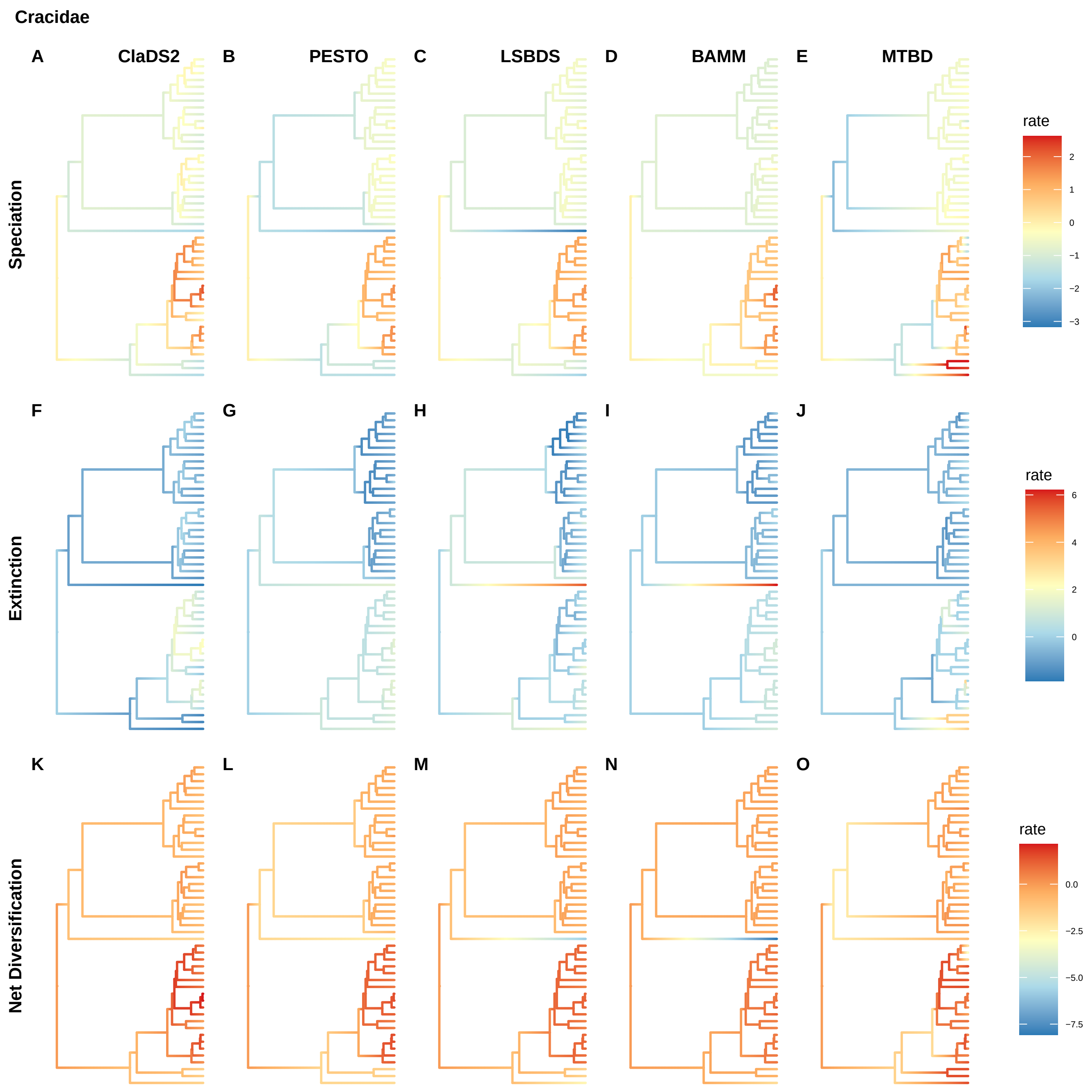
Cracidae
A
ClaDS2
B
PESTO
C
LSBDS
D
BAMM
E
MTBD
Speciation
rate
2
1
0
−1
−2
−3
F
G
H
I
J
Extinction
rate
6
4
2
0
K
L
M
N
O
Net Diversification
rate
0.0
−2.5
−5.0
−7.5

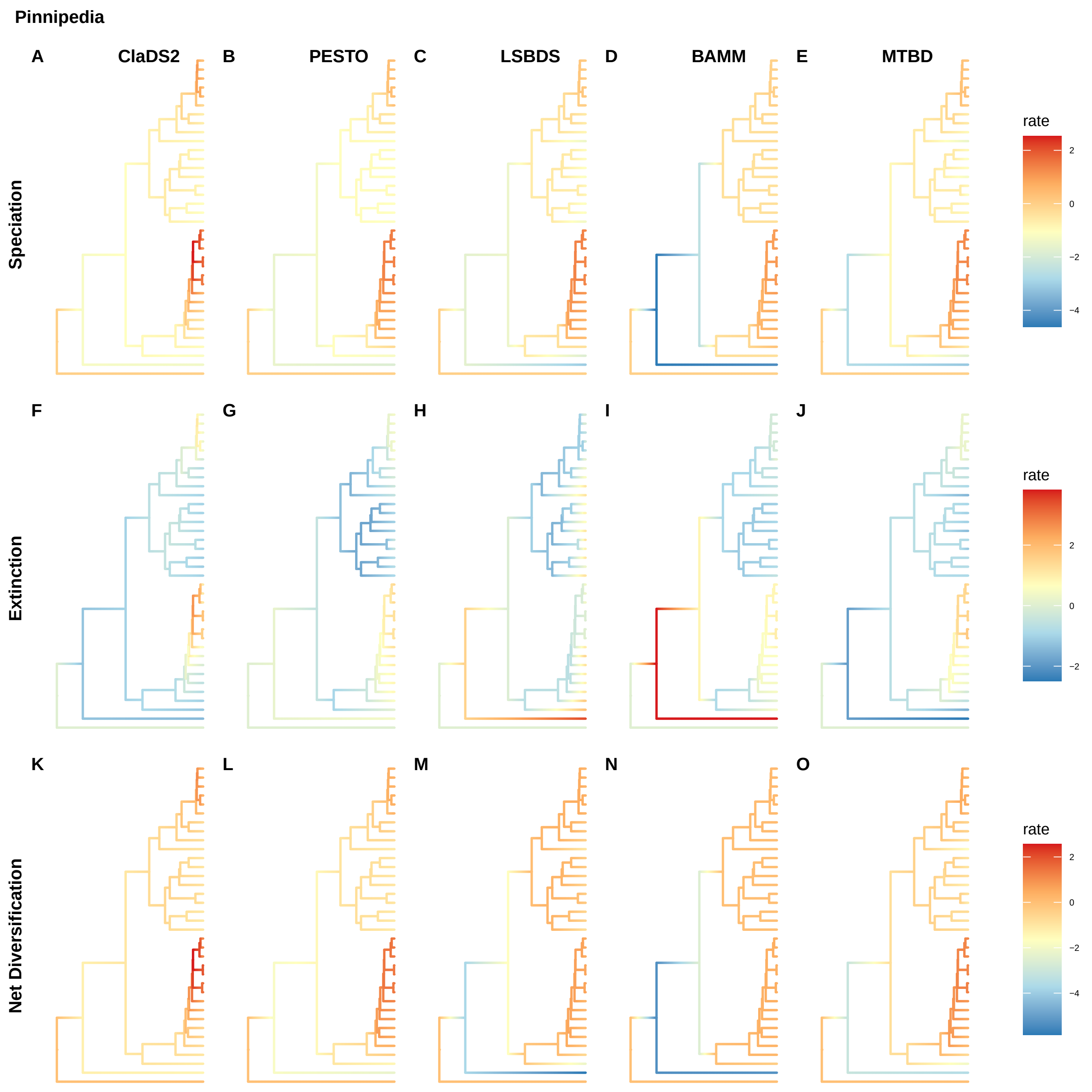
Pinnipedia
A
ClaDS2
B
PESTO
C
LSBDS
D
BAMM
E
MTBD
Speciation
rate
2
0
−2
−4
F
G
H
I
J
Extinction
rate
2
0
−2
K
L
M
N
O
Net Diversification
rate
2
0
−2
−4

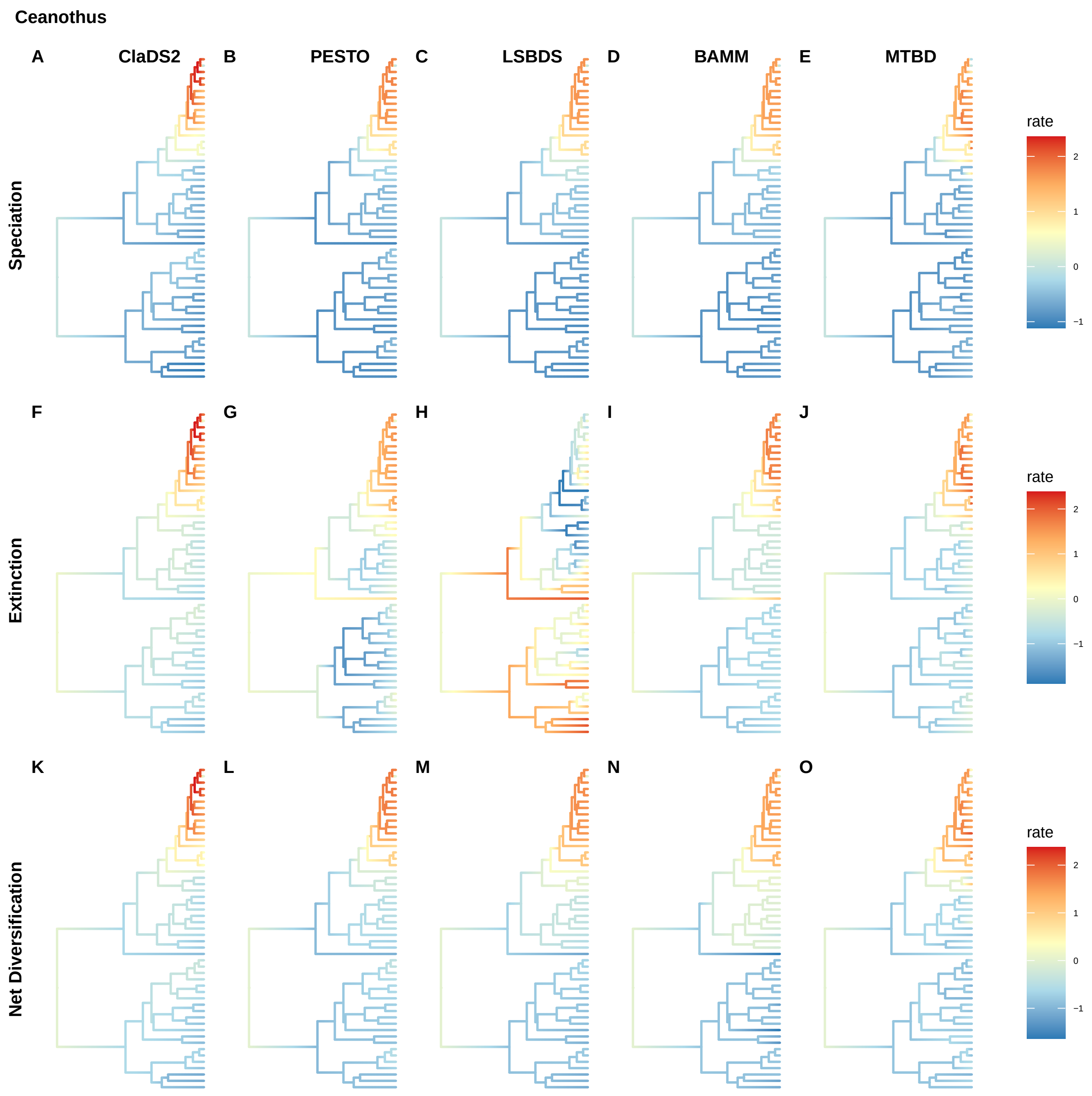

Ceanothus
A
ClaDS2
B
PESTO
C
LSBDS
D
BAMM
E
MTBD
Speciation
rate
2
1
0
−1
F
G
H
I
J
Extinction
rate
2
1
0
−1
K
L
M
N
O
Net Diversification
rate
2
1
0
−1

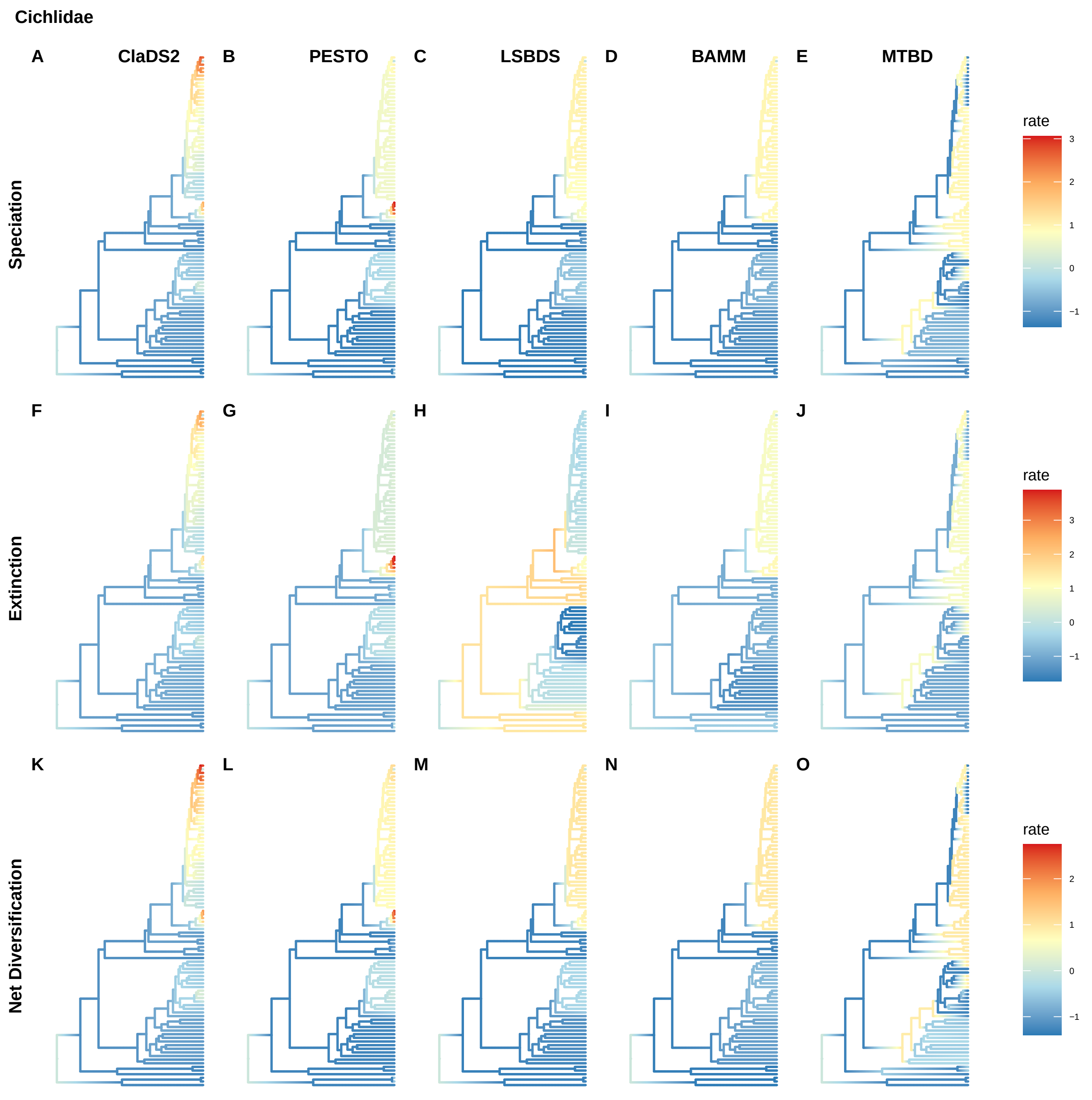
Cichlidae
A
ClaDS2
B
PESTO
C
LSBDS
D
BAMM
E
MTBD
F
G
H
I
J
K
L
M
N
O
Speciation
Extinction
Net Diversification
rate
3
2
1
0
−1
rate
3
2
1
0
−1
rate
2
1
0
−1

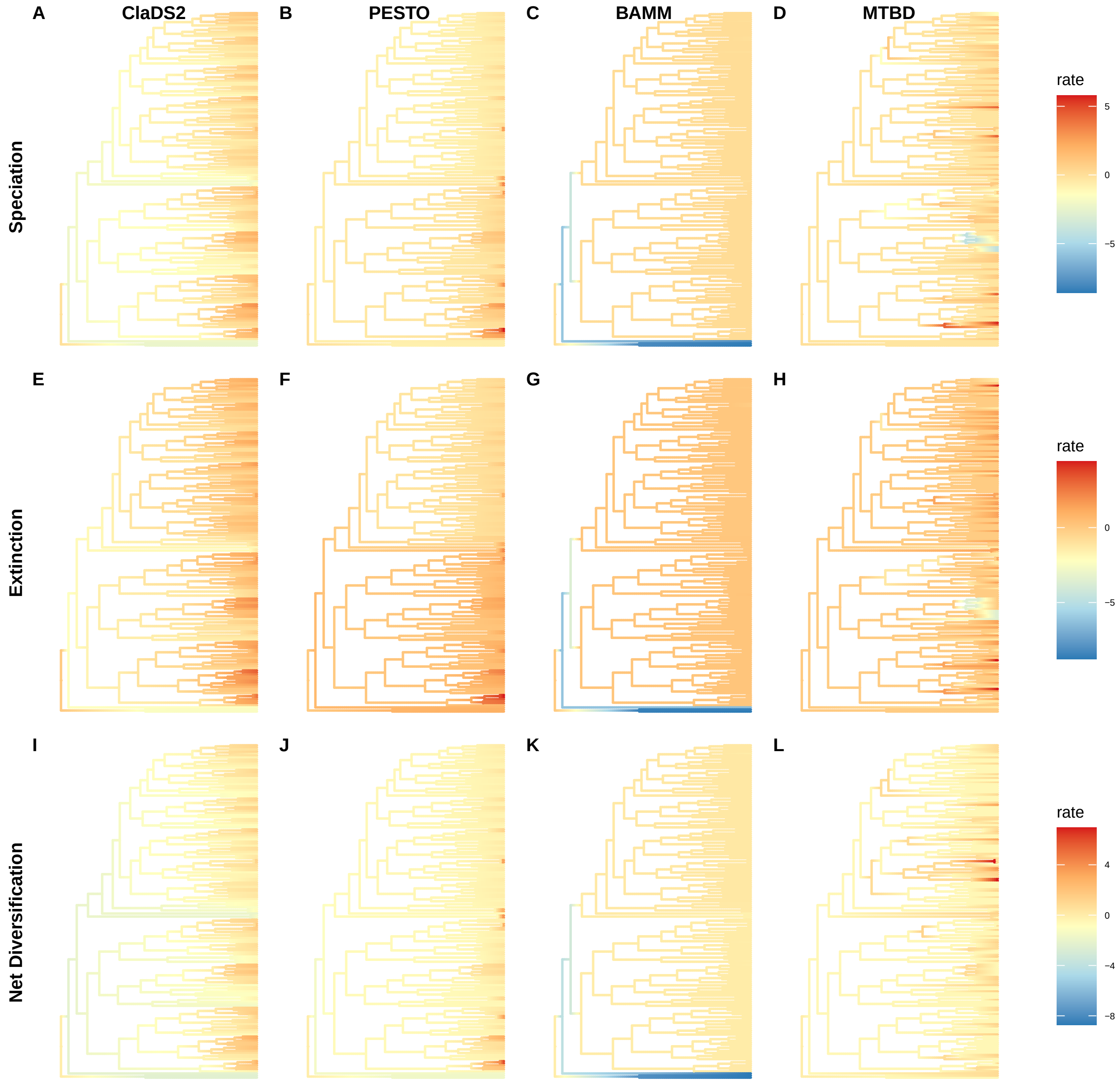
A
ClaDS2
B
PESTO
C
BAMM
D
MTBD
E
F
G
H
I
J
K
L
Speciation
Extinction
Net Diversification
rate
5
0
−5
rate
0
−5
rate
4
0
−4
−8

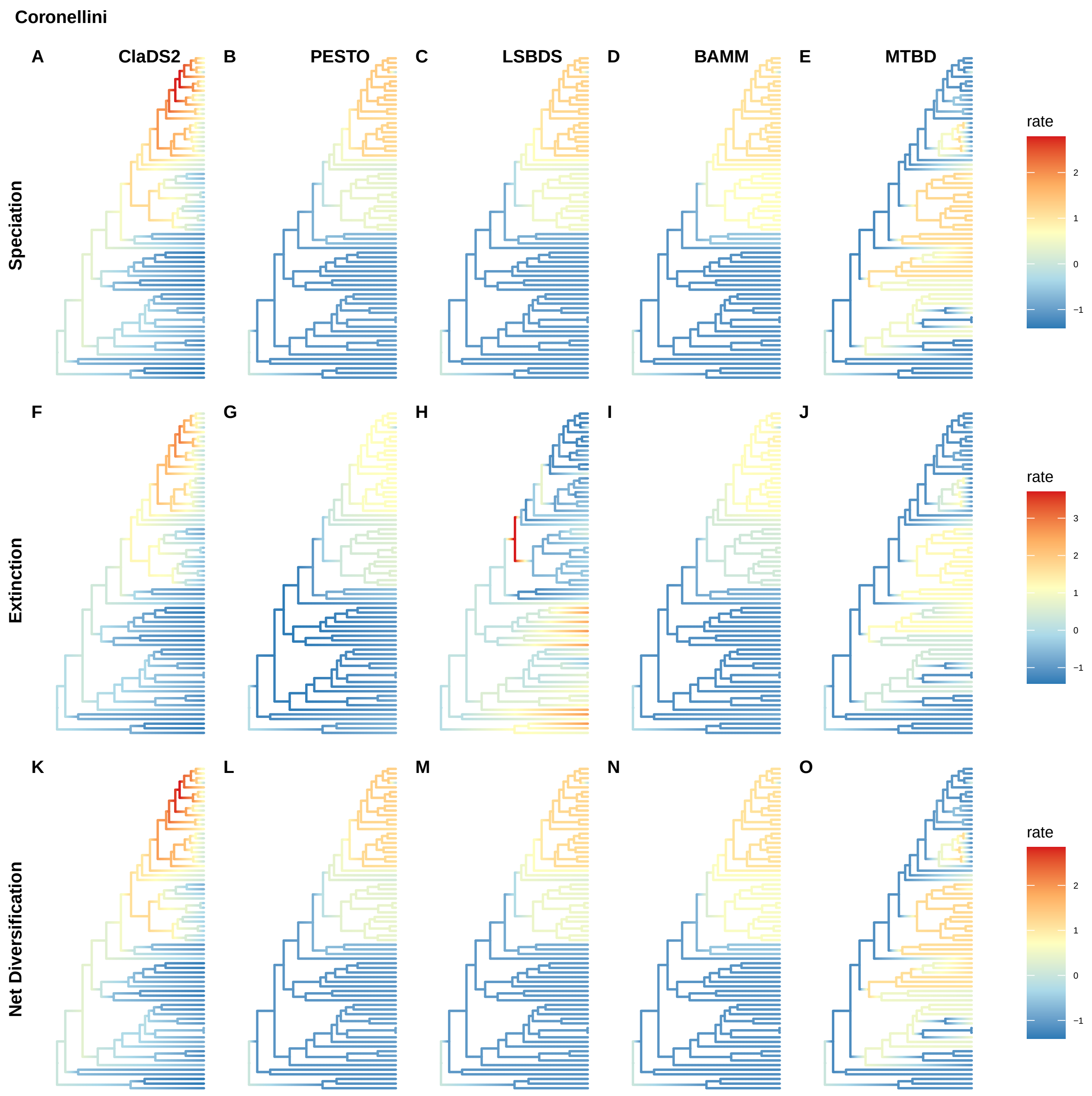
Coronellini
A
ClaDS2
B
PESTO
C
LSBDS
D
BAMM
E
MTBD
Speciation
rate
2
1
0
−1
F
G
H
I
J
Extinction
rate
3
2
1
0
−1
K
L
M
N
O
Net Diversification
rate
2
1
0
−1

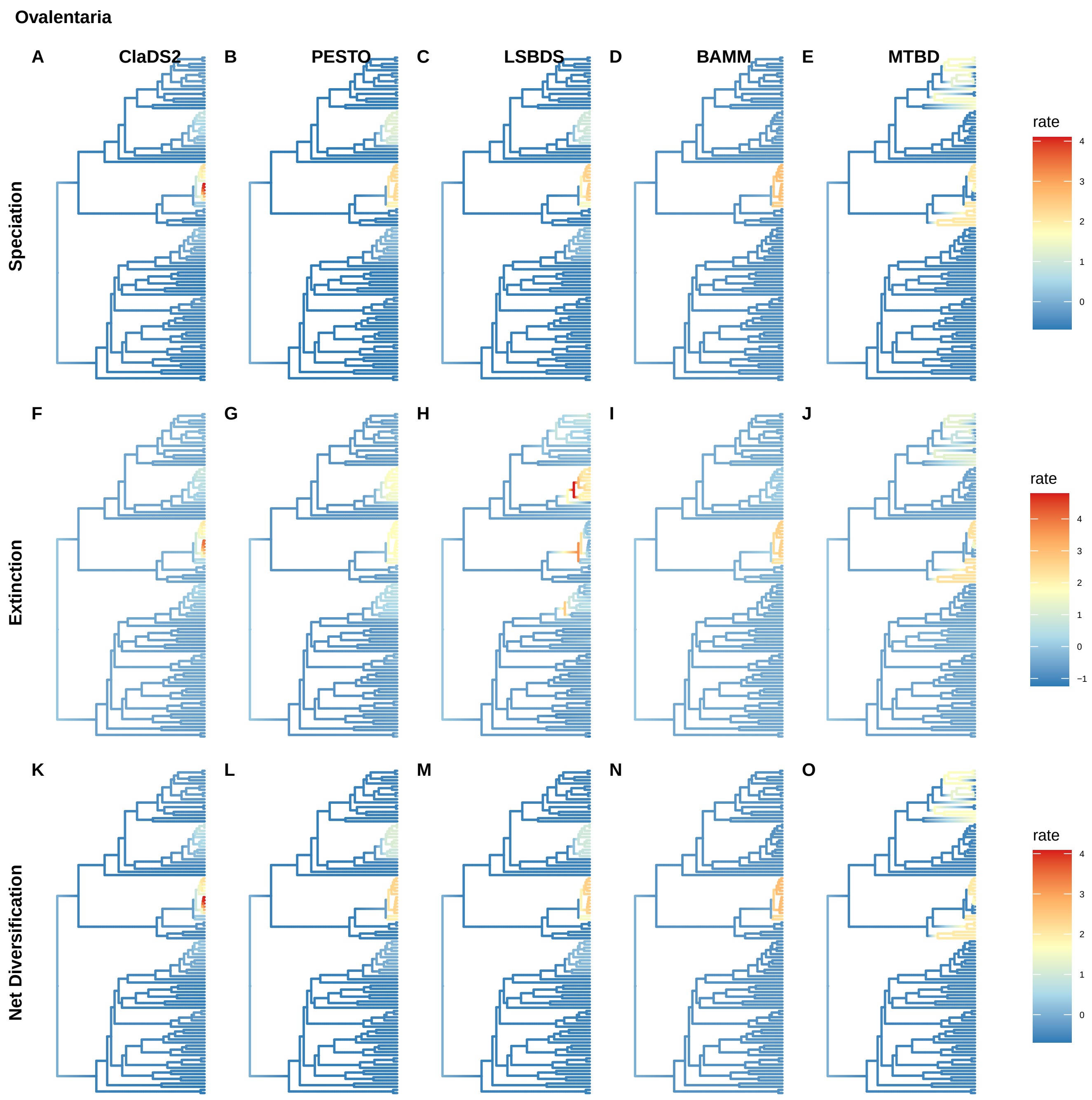

Ovalentaria
A
ClaDS2
B
PESTO
C
LSBDS
D
BAMM
E
MTBD
Speciation
rate
4
3
2
1
0
F
G
H
I
J
Extinction
rate
4
3
2
1
0
−1
K
L
M
N
O
Net Diversification
rate
4
3
2
1
0

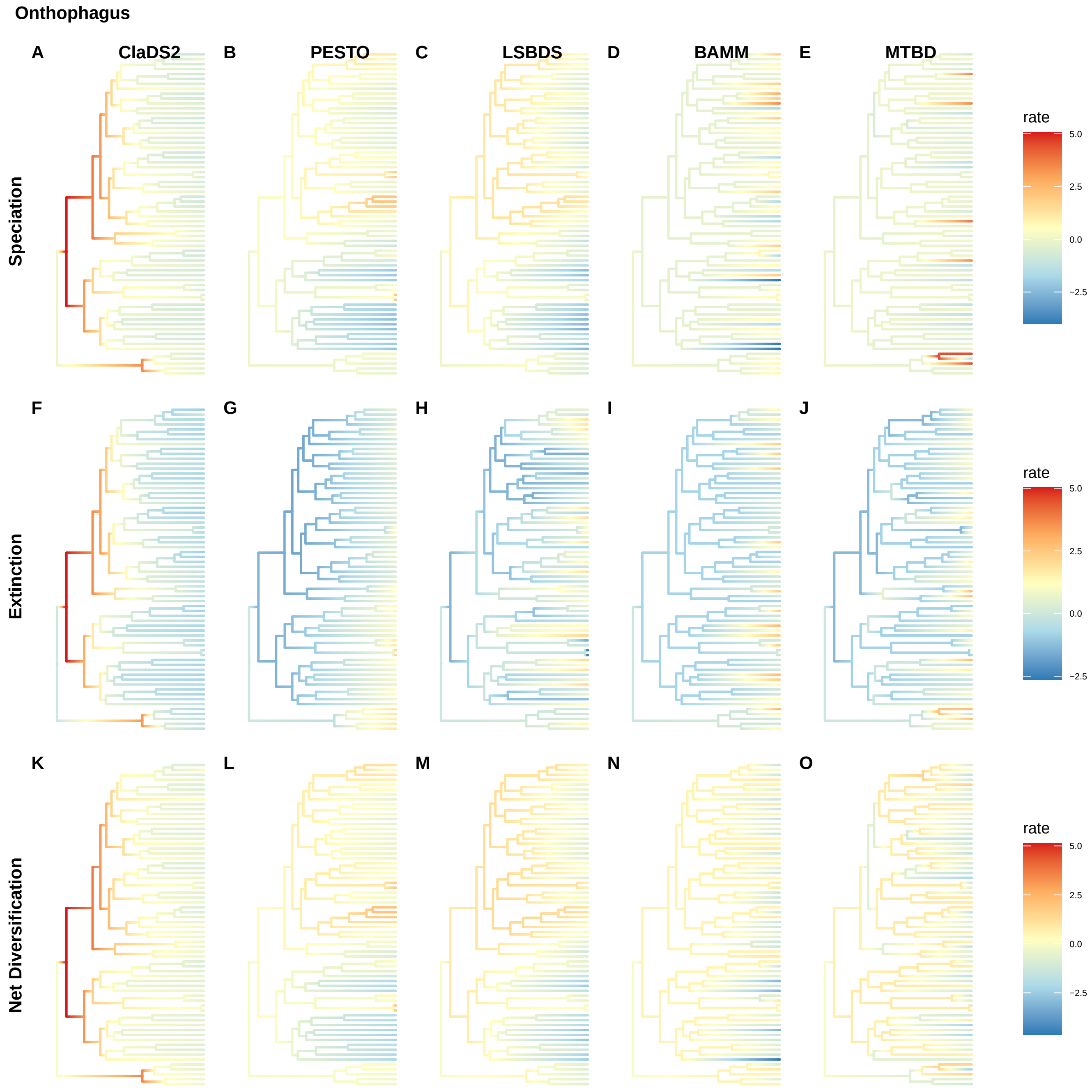
Onthophagus
A
ClaDS2
B
PESTO
C
LSBDS
D
BAMM
E
MTBD
Speciation
F
G
H
I
J
Extinction
K
L
M
N
O
Net Diversification
rate
5.0
2.5
0.0
−2.5
rate
5.0
2.5
0.0
−2.5
rate
5.0
2.5
0.0
−2.5

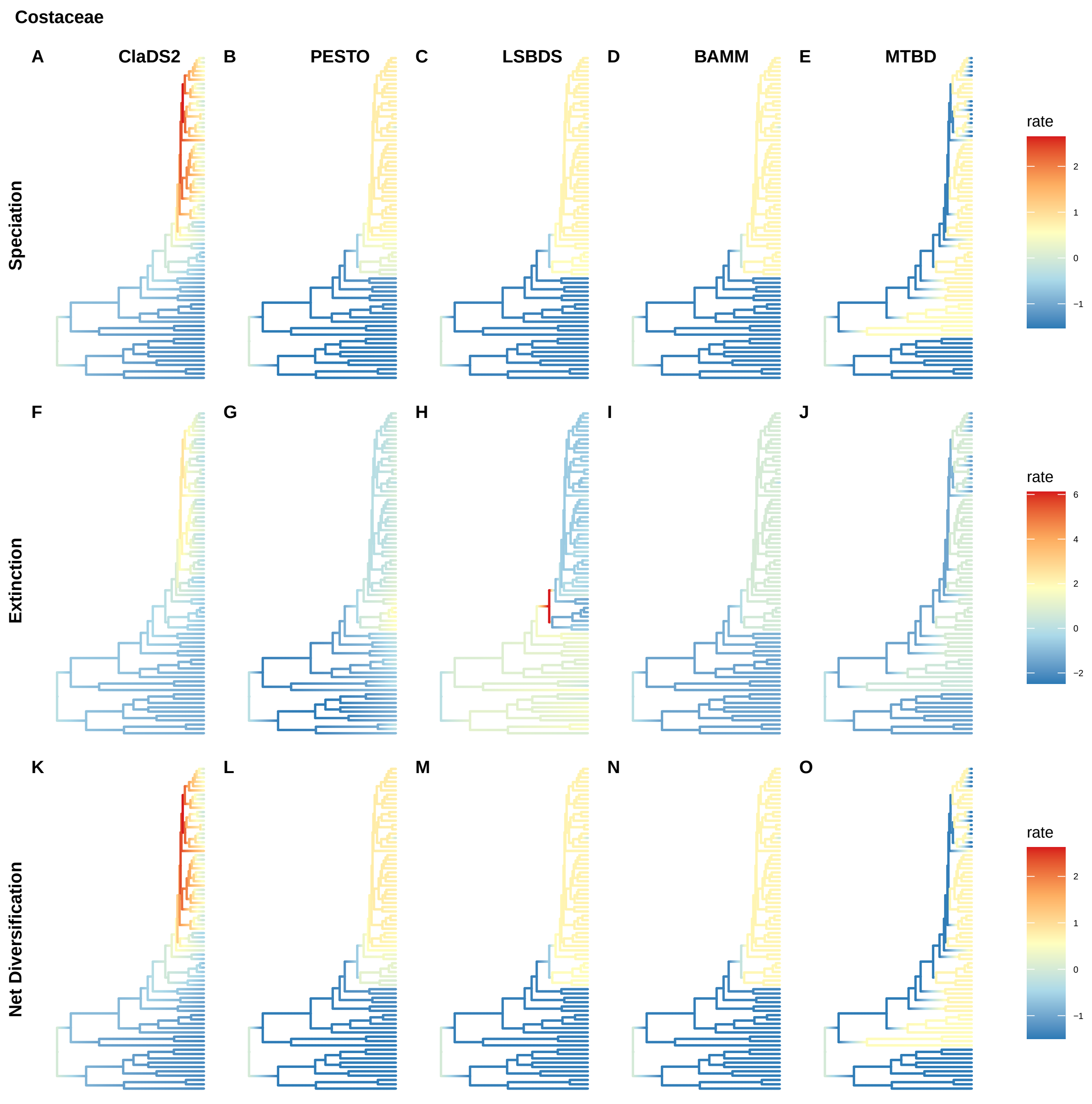

Costaceae
A
ClaDS2
B
PESTO
C
LSBDS
D
BAMM
E
MTBD
Speciation
rate
2
1
0
−1
F
G
H
I
J
Extinction
rate
6
4
2
0
−2
K
L
M
N
O
Net Diversification
rate
2
1
0
−1

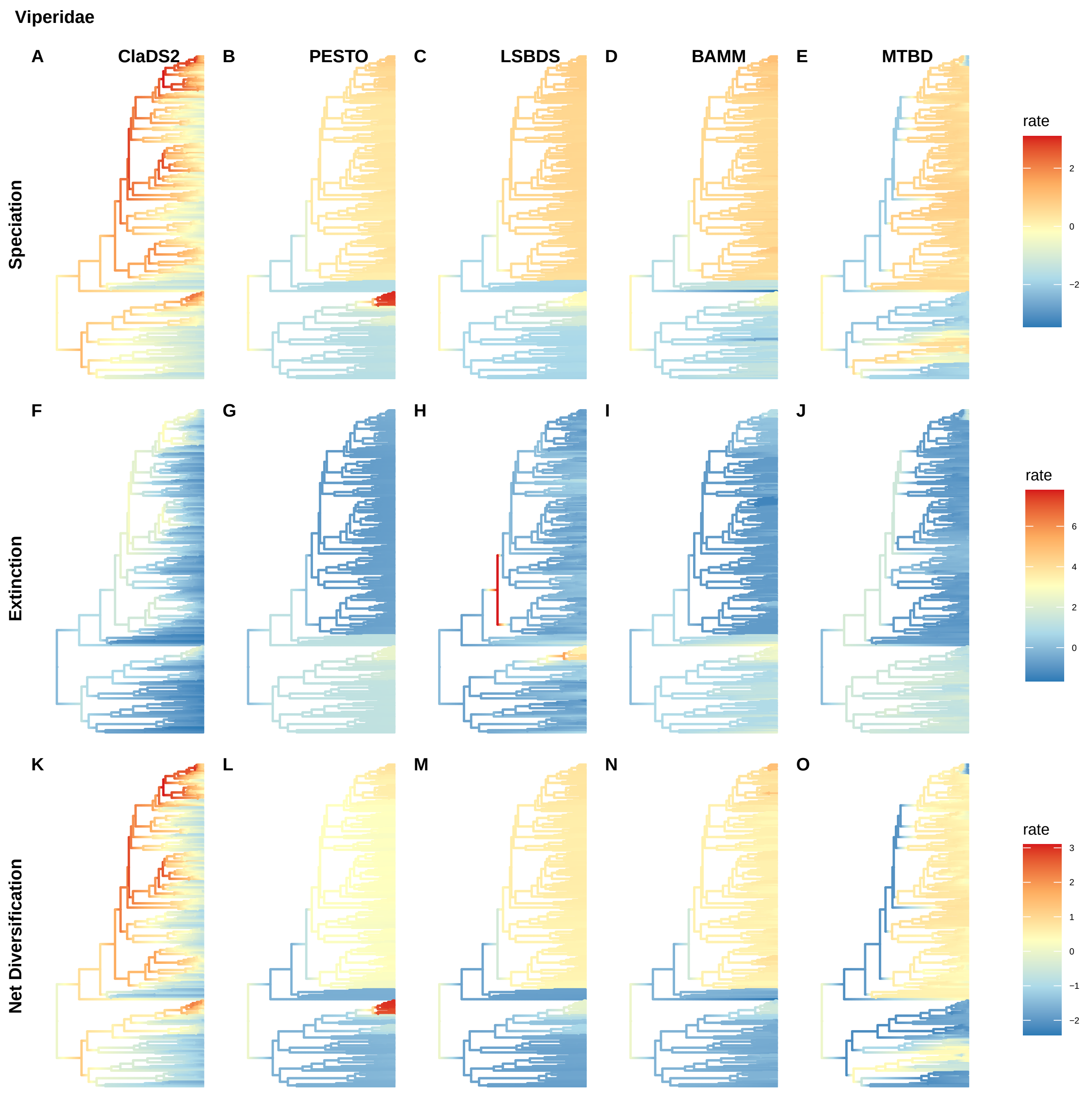

Viperidae
A
ClaDS2
B
PESTO
C
LSBDS
D
BAMM
E
MTBD
Speciation
rate
2
0
−2
F
G
H
I
J
Extinction
rate
6
4
2
0
K
L
M
N
O
Net Diversification
rate
3
2
1
0
−1
−2

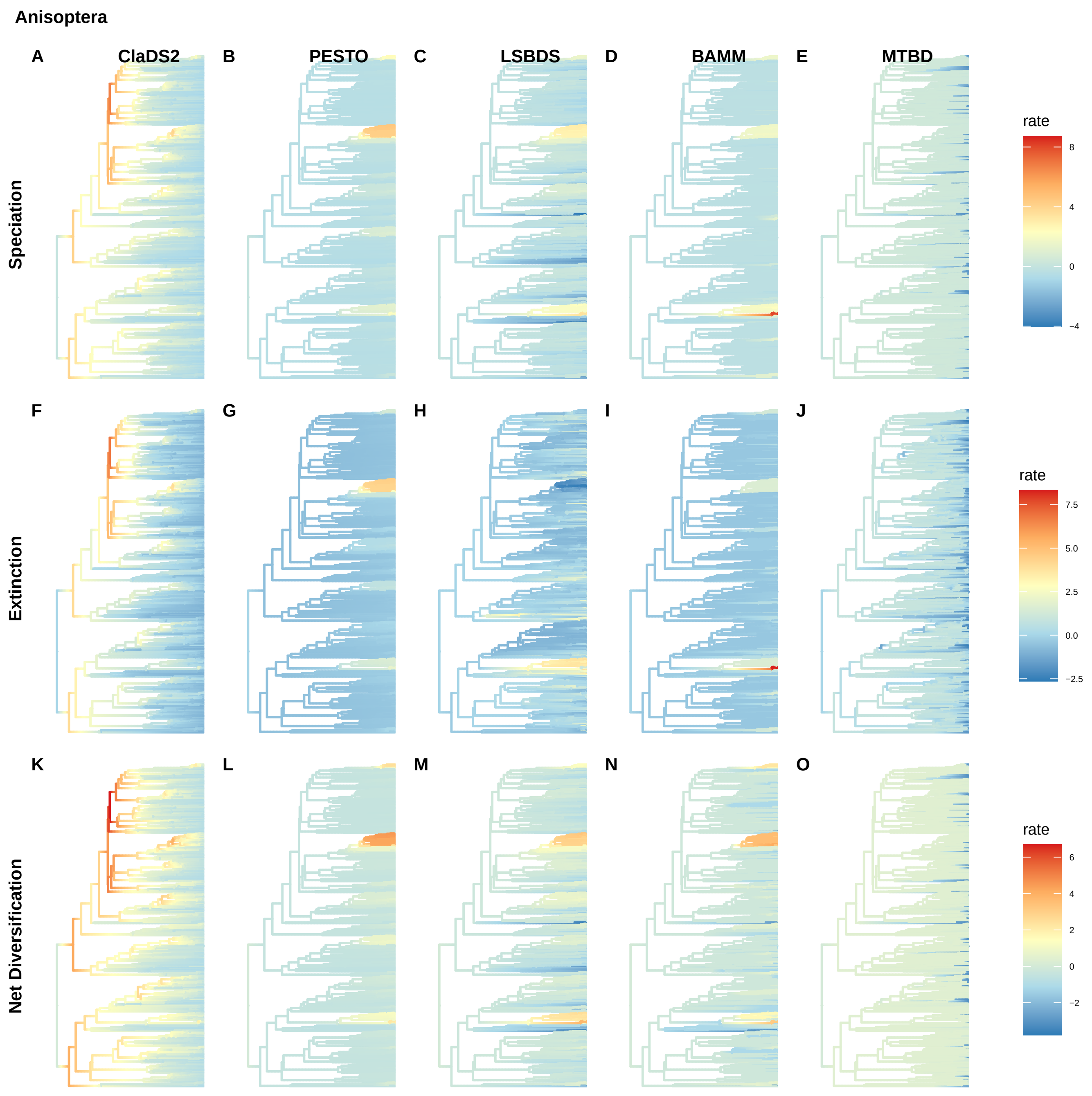
Anisoptera
A
ClaDS2
B
PESTO
C
LSBDS
D
BAMM
E
MTBD
Speciation
rate
8
4
0
−4
F
G
H
I
J
Extinction
rate
7.5
5.0
2.5
0.0
−2.5
K
L
M
N
O
Net Diversification
rate
6
4
2
0
−2

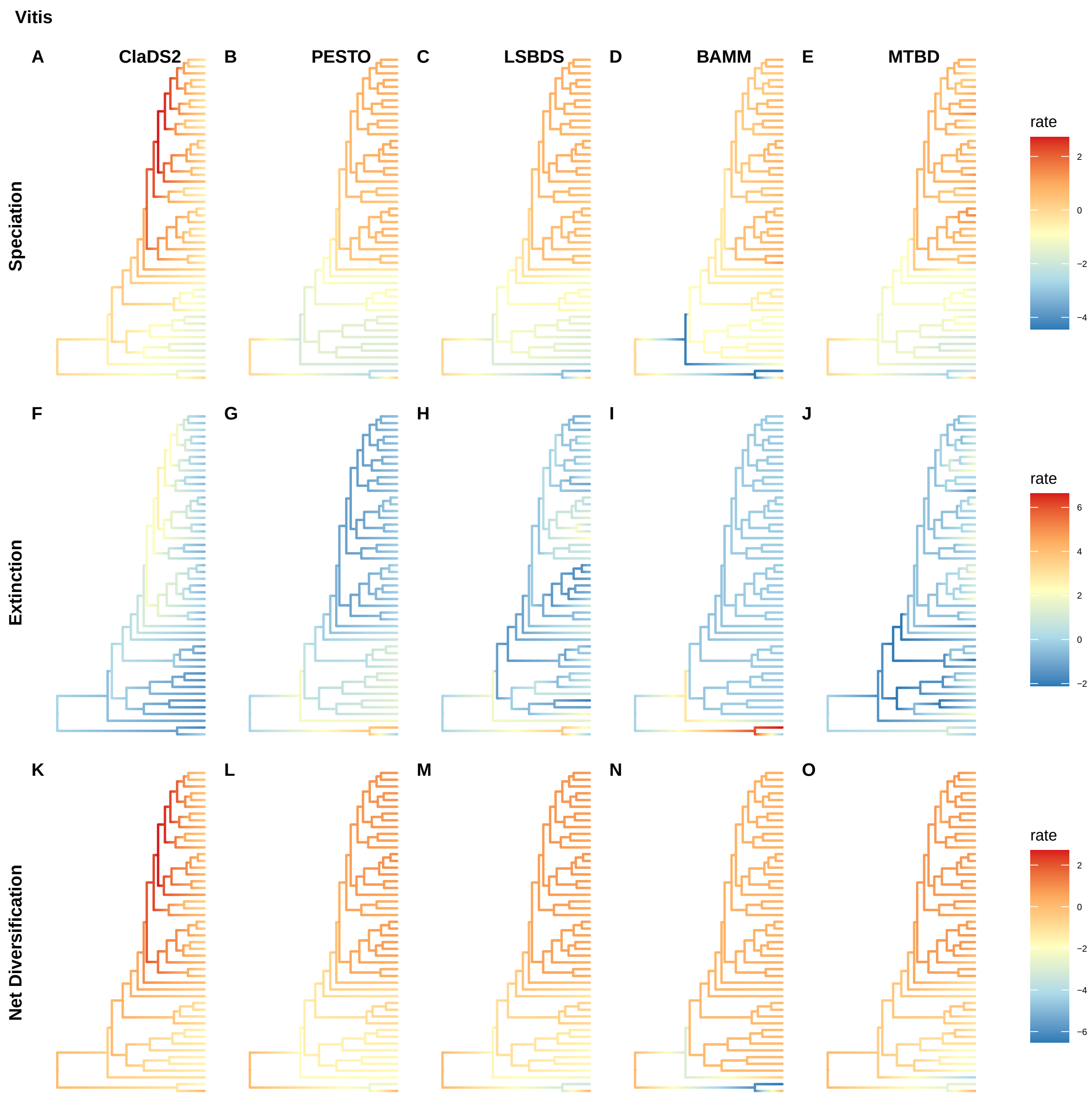
Vitis
A
ClaDS2
B
PESTO
C
LSBDS
D
BAMM
E
MTBD
Speciation
rate
2
0
−2
−4
F
G
H
I
J
Extinction
rate
6
4
2
0
−2
K
L
M
N
O
Net Diversification
rate
2
0
−2
−4
−6

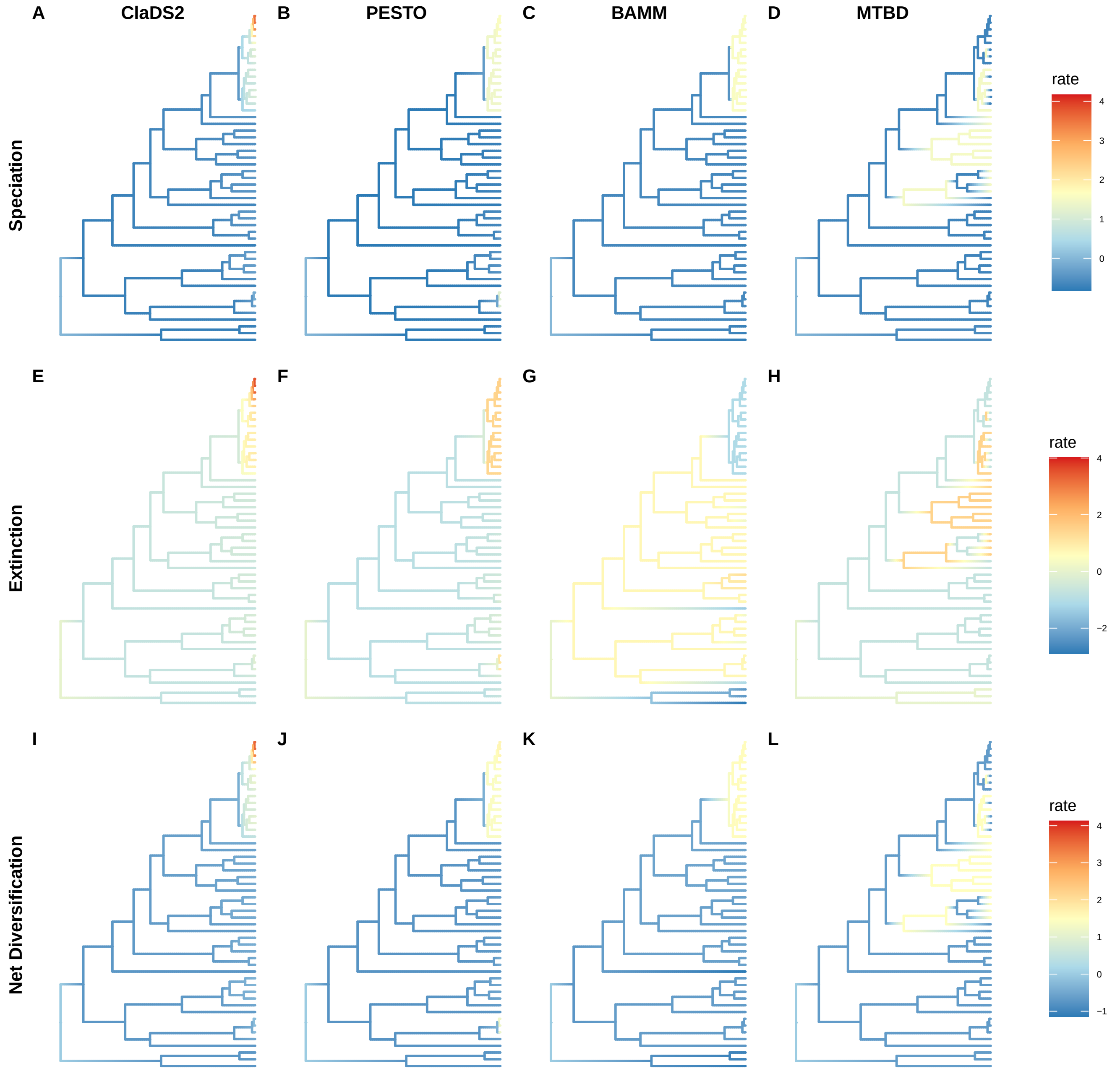

A
ClaDS2
B
PESTO
C
BAMM
D
MTBD
Speciation
rate
4
3
2
1
0
E
F
G
H
Extinction
rate
4
2
0
−2
I
J
K
L
Net Diversification
rate
4
3
2
1
0
−1

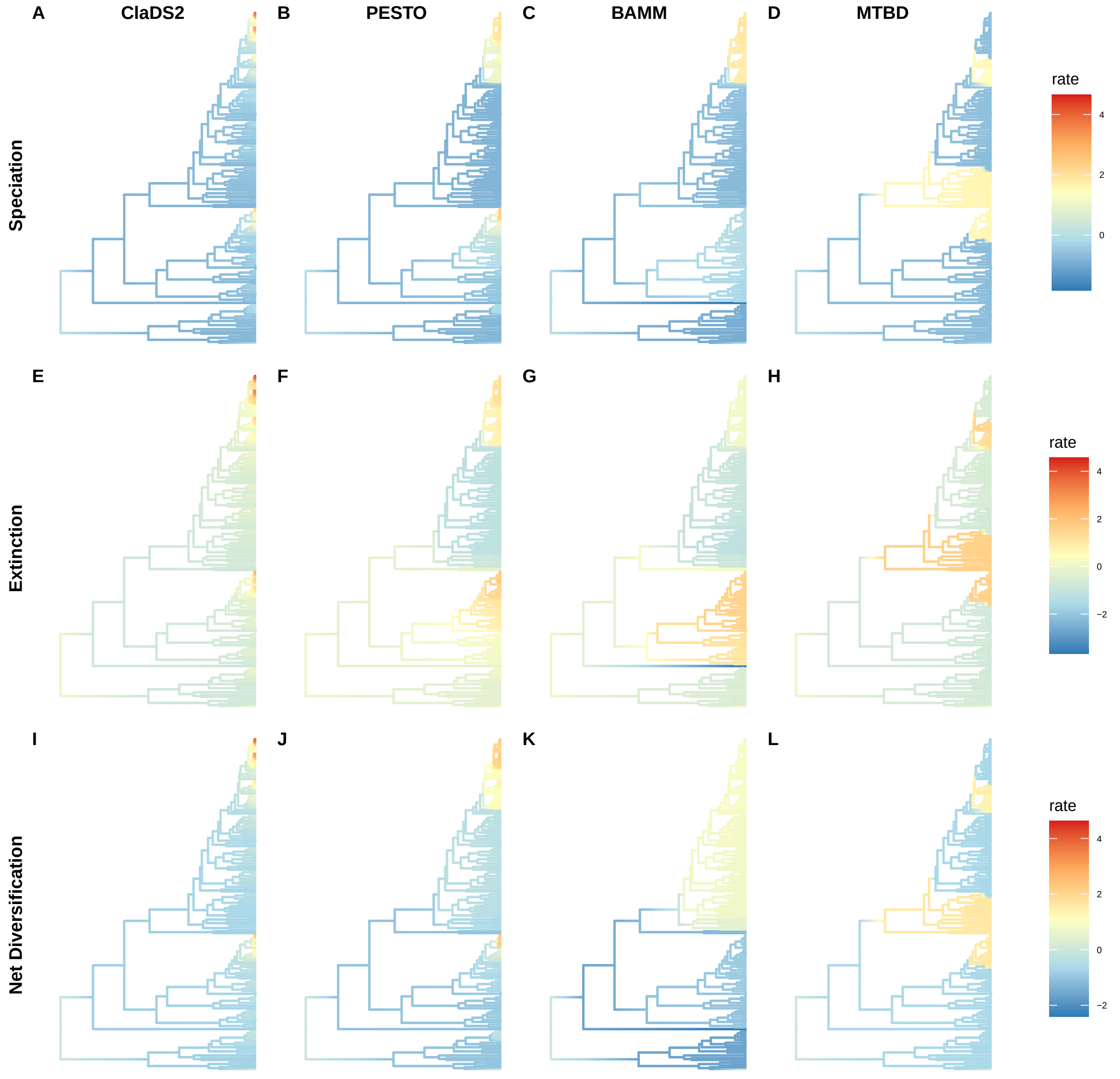

A
ClaDS2
B
PESTO
C
BAMM
D
MTBD
Speciation
rate
4
2
0
E
F
G
H
Extinction
rate
4
2
0
−2
I
J
K
L
Net Diversification
rate
4
2
0
−2

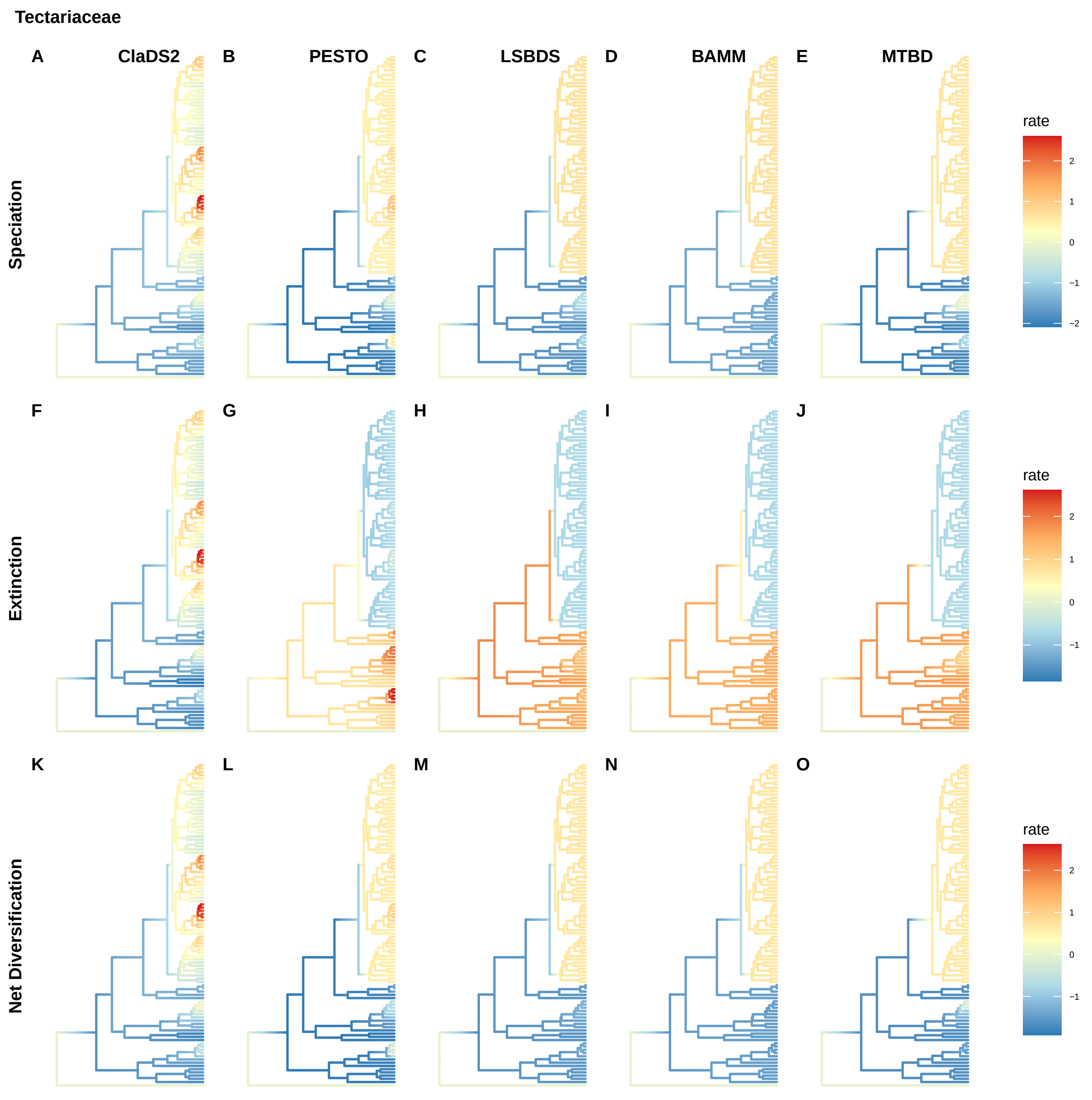
Tectariaceae
A
ClaDS2
B
PESTO
C
LSBDS
D
BAMM
E
MTBD
Speciation
rate
2
1
0
−1
−2
F
G
H
I
J
Extinction
rate
2
1
0
−1
K
L
M
N
O
Net Diversification
rate
2
1
0
−1

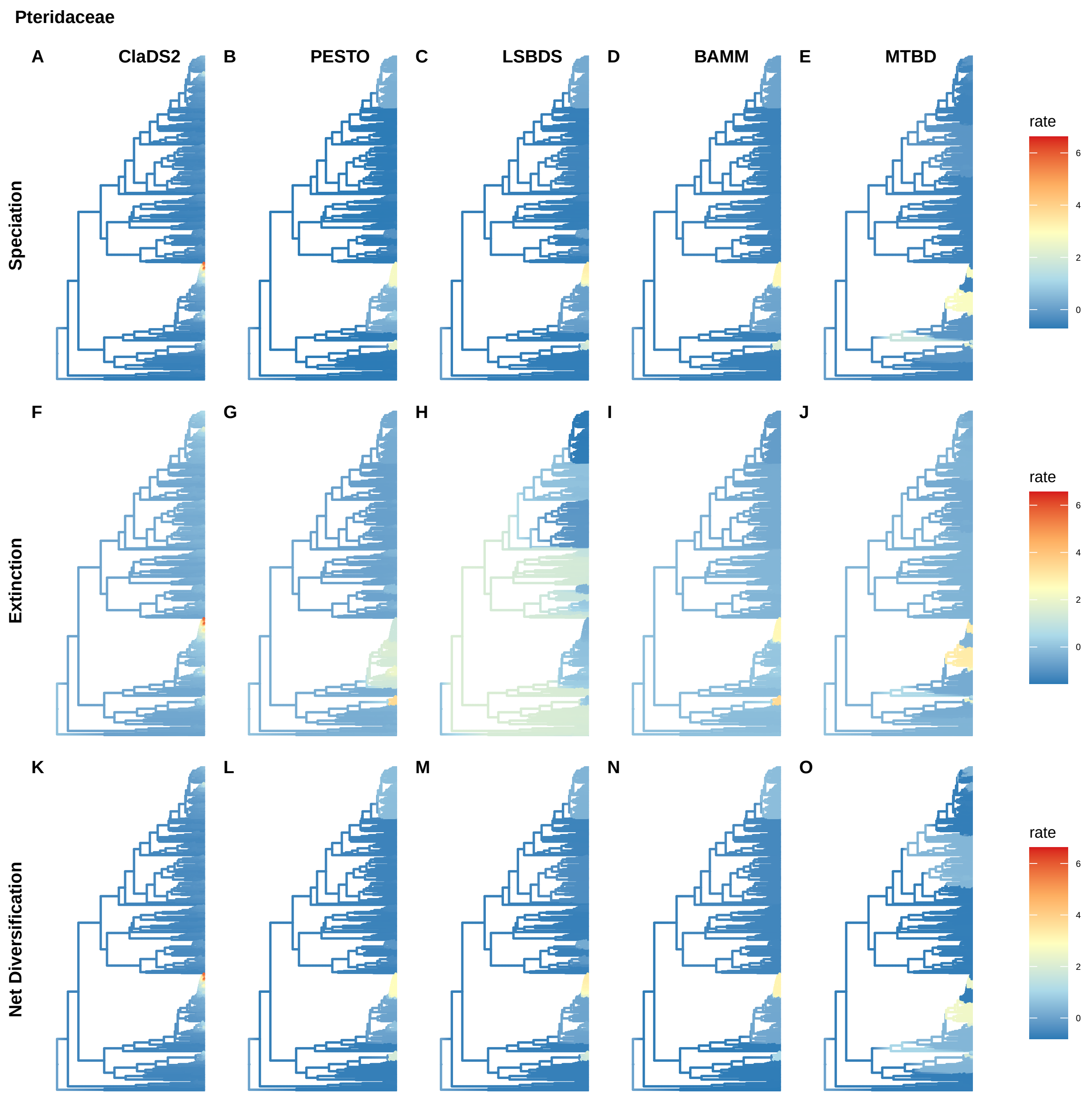

Pteridaceae
A
ClaDS2
B
PESTO
C
LSBDS
D
BAMM
E
MTBD
Speciation
rate
6
4
2
0
F
G
H
I
J
Extinction
rate
6
4
2
0
K
L
M
N
O
Net Diversification
rate
6
4
2
0

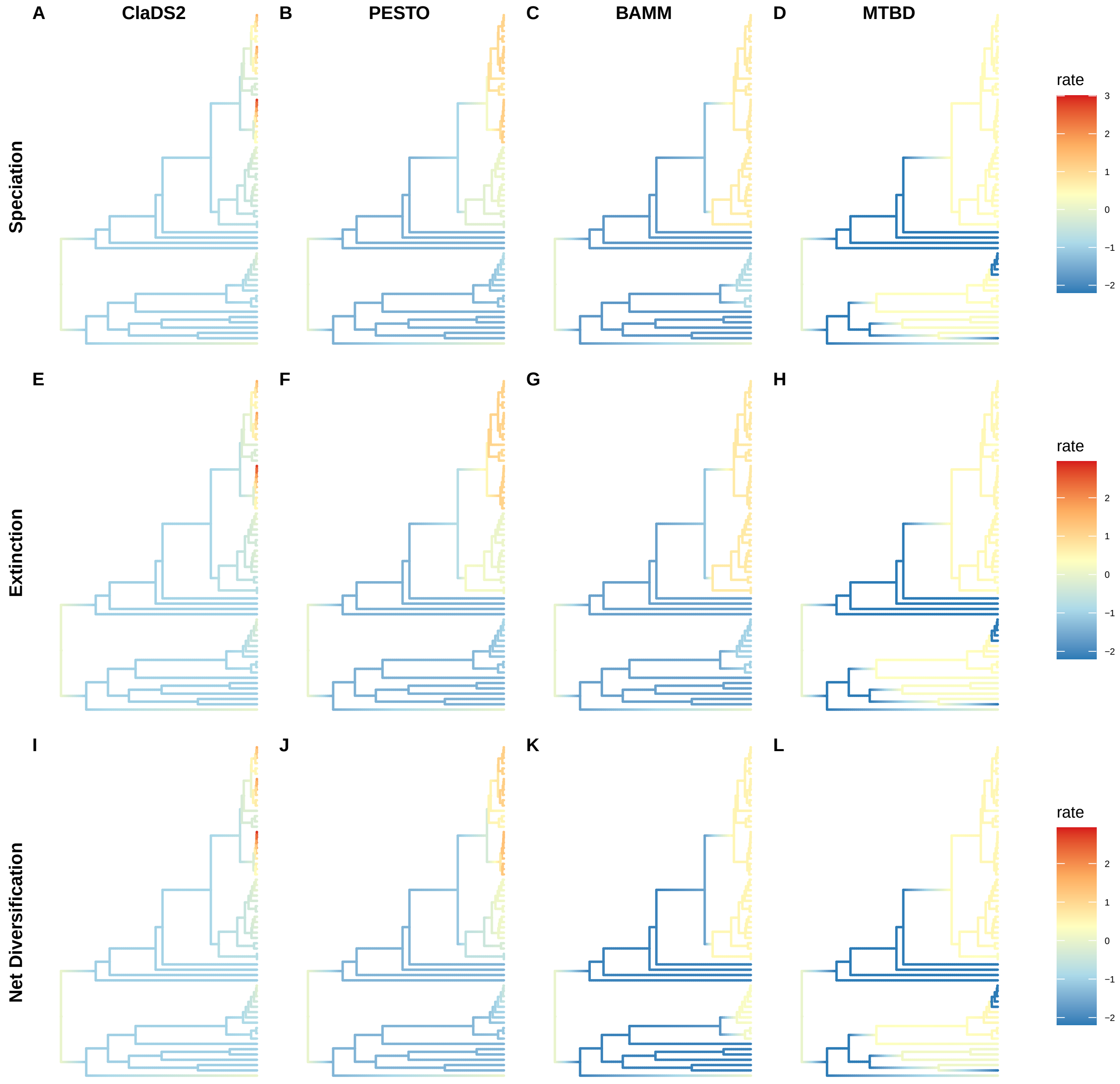

A
ClaDS2
B
PESTO
C
BAMM
D
MTBD
Speciation
rate
3
2
1
0
−1
−2
E
F
G
H
Extinction
rate
2
1
0
−1
−2
I
J
K
L
Net Diversification
rate
2
1
0
−1
−2

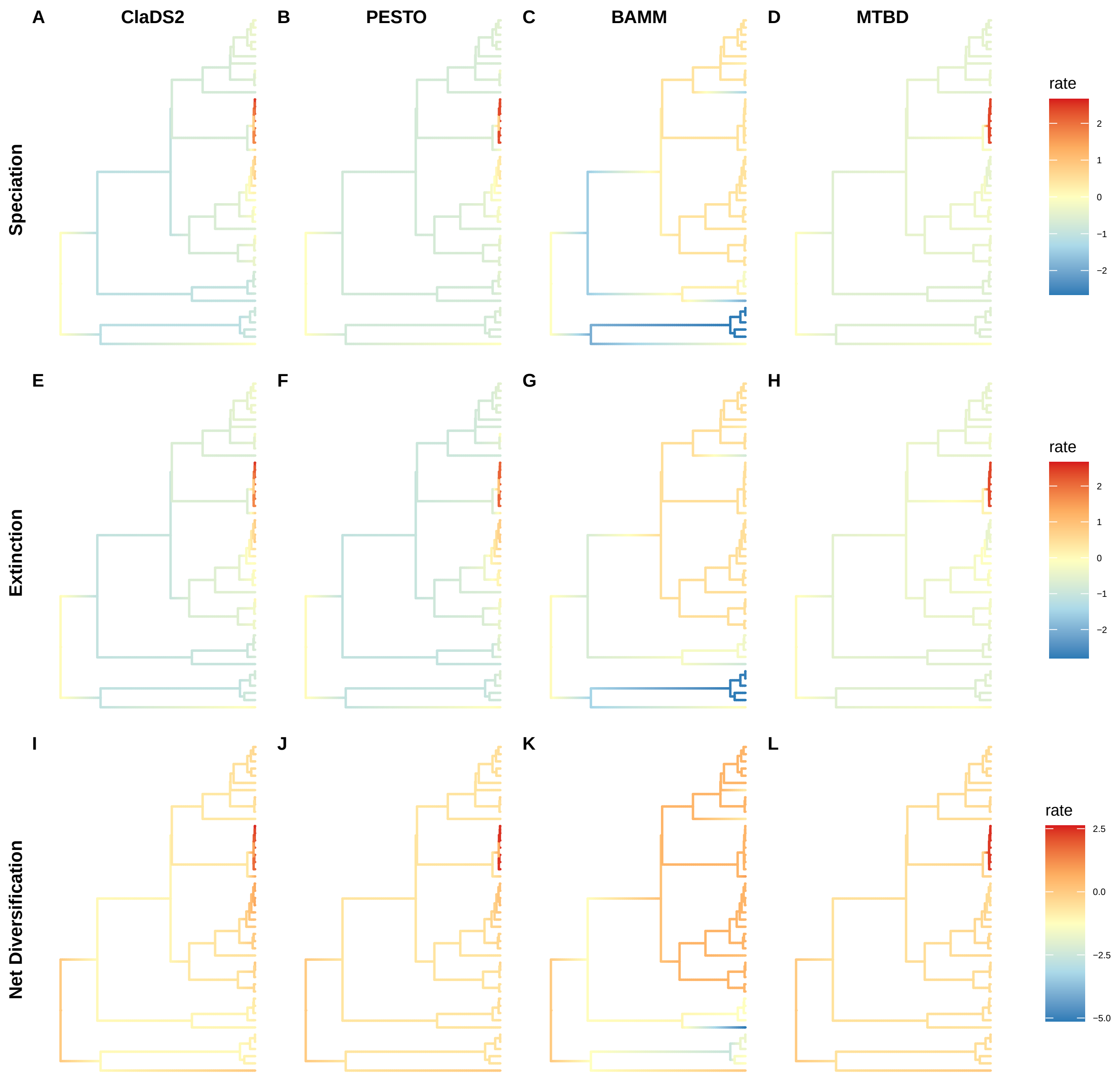
A
ClaDS2
B
PESTO
C
BAMM
D
MTBD
Speciation
rate
2
1
0
−1
−2
E
F
G
H
Extinction
rate
2
1
0
−1
−2
I
J
K
L
Net Diversification
rate
2.5
0.0
−2.5
−5.0

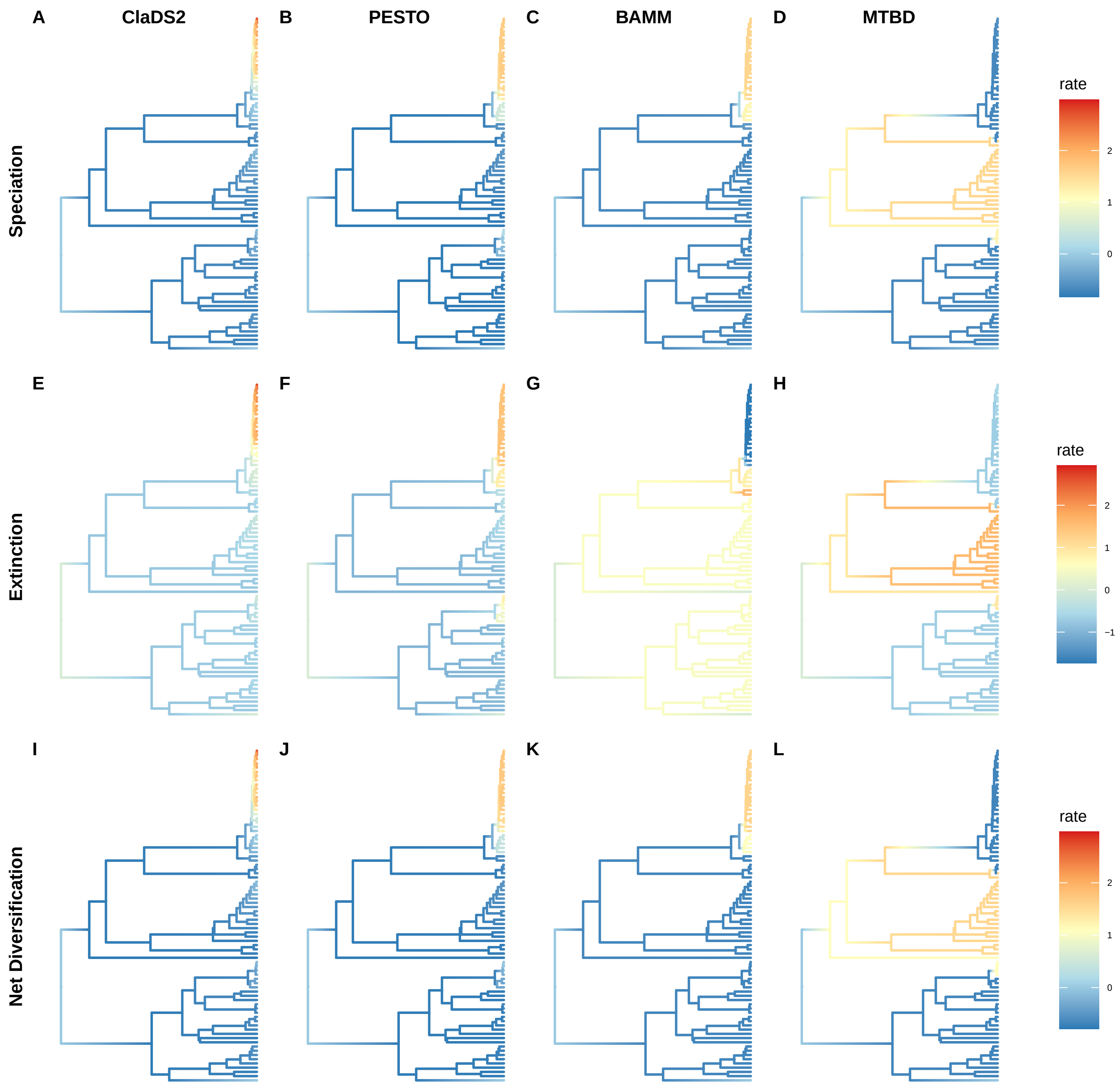
A
ClaDS2
B
PESTO
C
BAMM
D
MTBD
Speciation
rate
2
1
0
E
F
G
H
Extinction
rate
2
1
0
−1
I
J
K
L
Net Diversification
rate
2
1
0

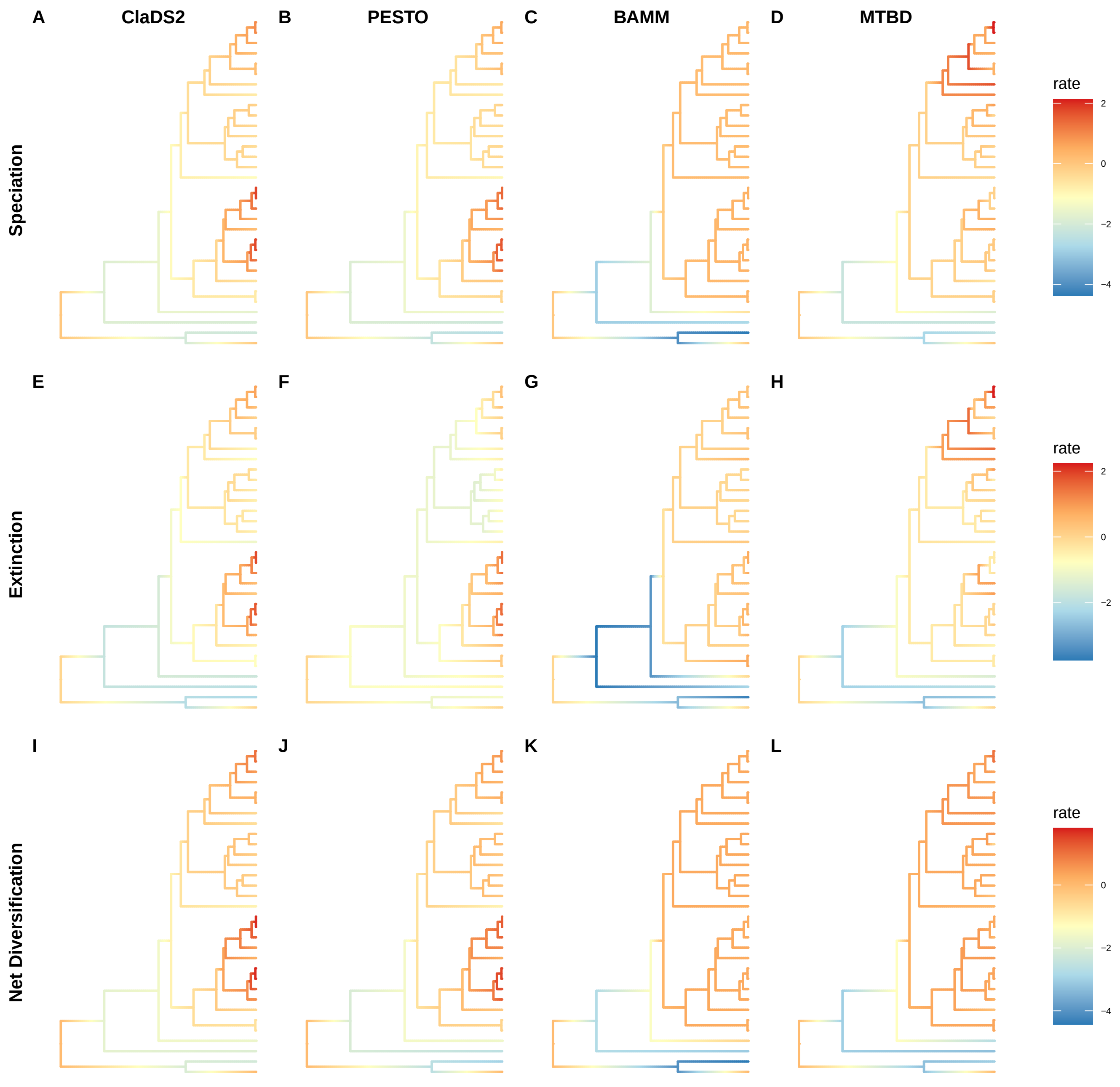

A
ClaDS2
B
PESTO
C
BAMM
D
MTBD
Speciation
rate
2
0
−2
−4
E
F
G
H
Extinction
rate
2
0
−2
I
J
K
L
Net Diversification
rate
0
−2
−4

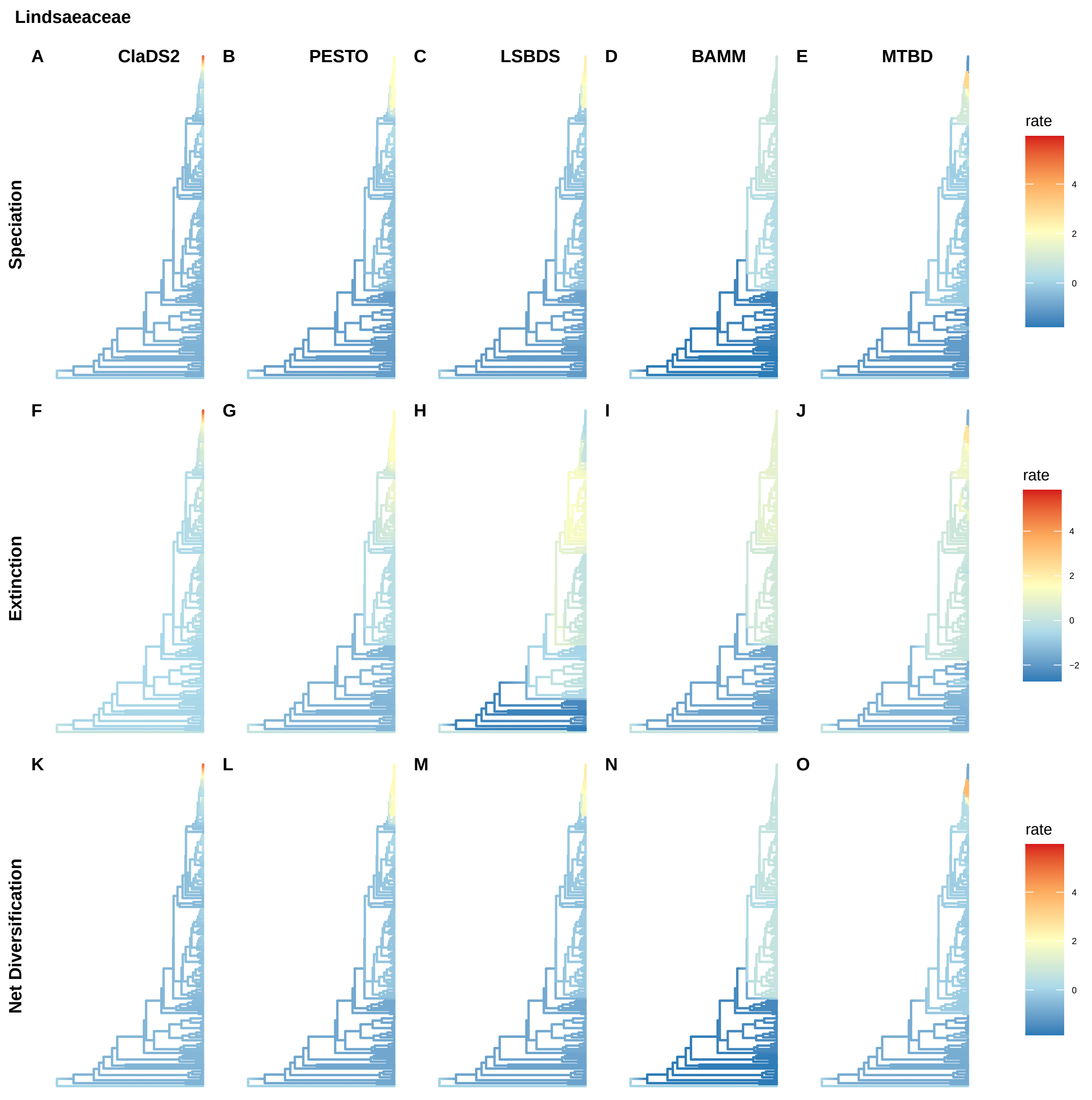
Lindsaeaceae
A
ClaDS2
B
PESTO
C
LSBDS
D
BAMM
E
MTBD
Speciation
rate
4
2
0
F
G
H
I
J
Extinction
rate
4
2
0
−2
K
L
M
N
O
Net Diversification
rate
4
2
0

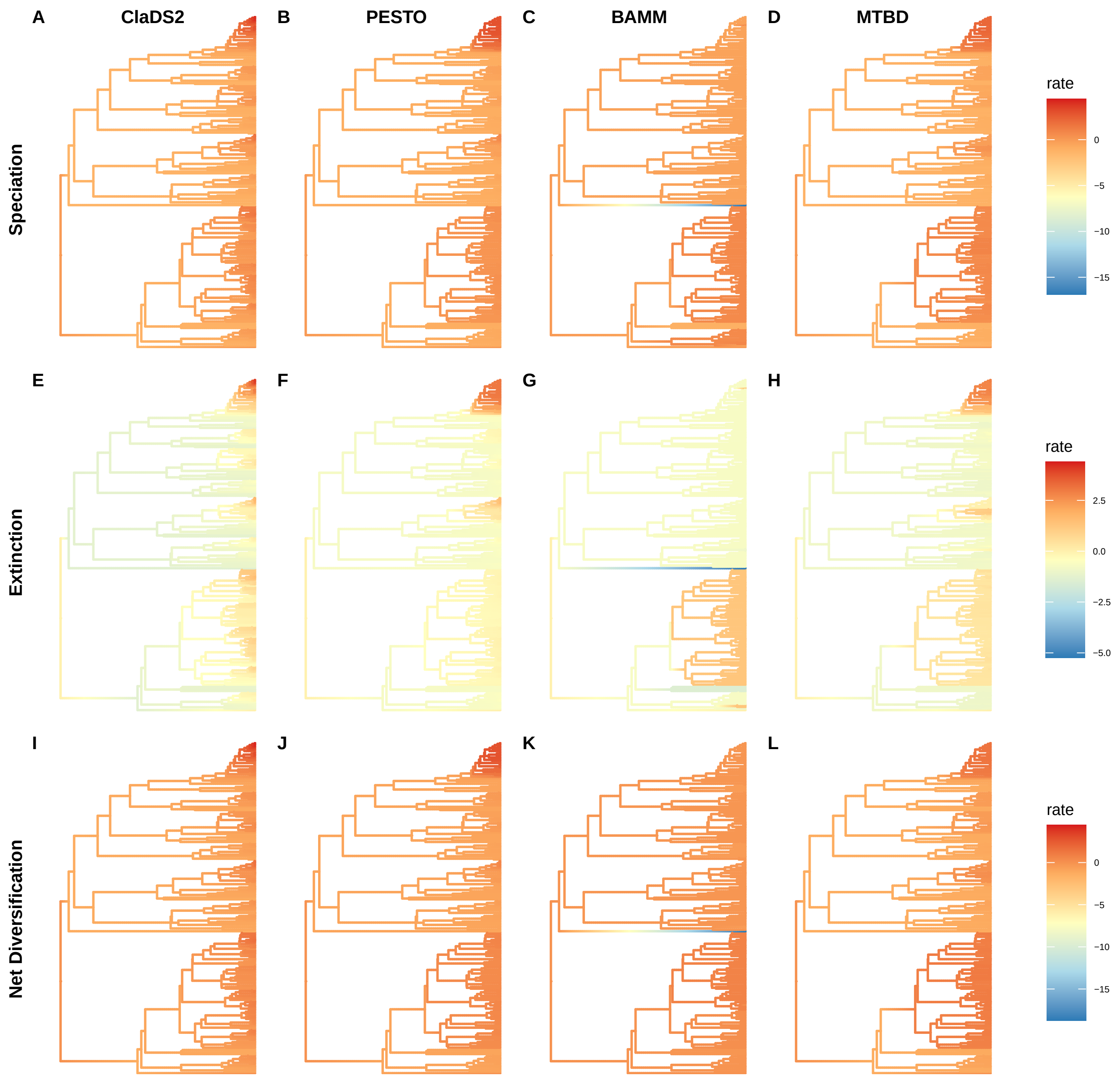
A
ClaDS2
B
PESTO
C
BAMM
D
MTBD
Speciation
rate
0
−5
−10
−15
E
F
G
H
Extinction
rate
2.5
0.0
−2.5
−5.0
I
J
K
L
Net Diversification
rate
0
−5
−10
−15

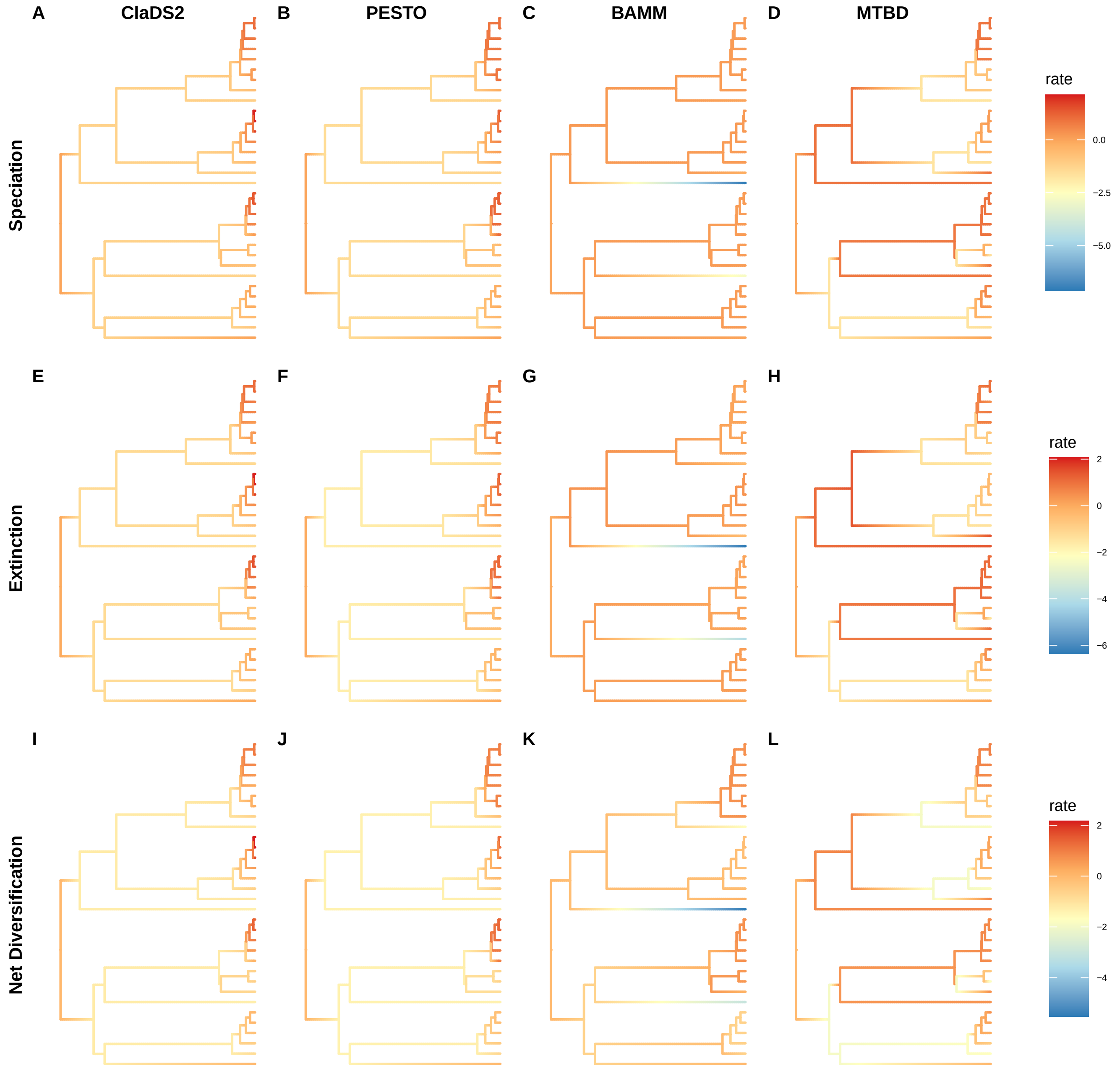
A
ClaDS2
B
PESTO
C
BAMM
D
MTBD
Speciation
rate
0.0
−2.5
−5.0
E
F
G
H
Extinction
rate
2
0
−2
−4
−6
I
J
K
L
Net Diversification
rate
2
0
−2
−4

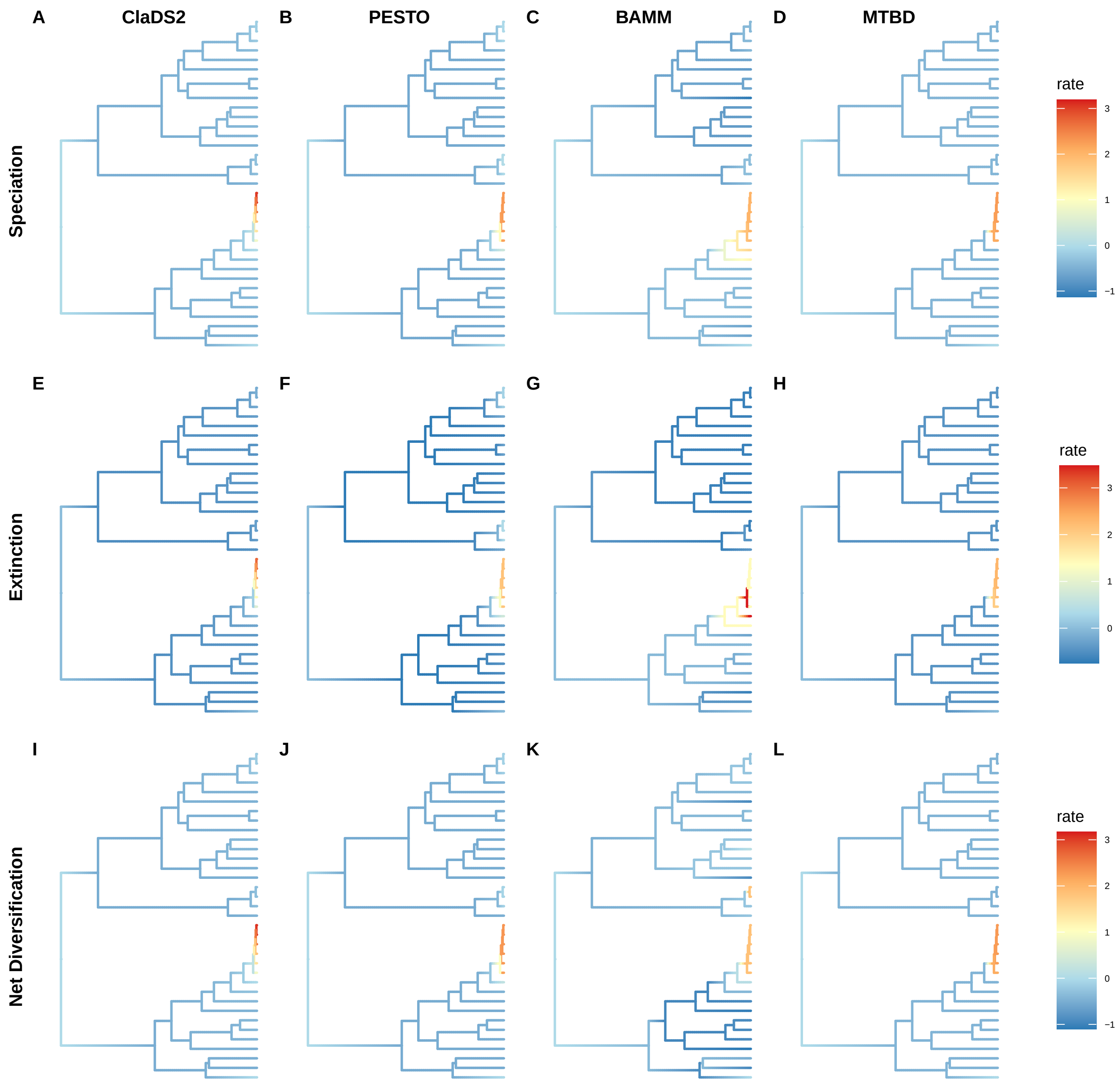
A
ClaDS2
B
PESTO
C
BAMM
D
MTBD
Speciation
rate
3
2
1
0
−1
E
F
G
H
Extinction
rate
3
2
1
0
I
J
K
L
Net Diversification
rate
3
2
1
0
−1

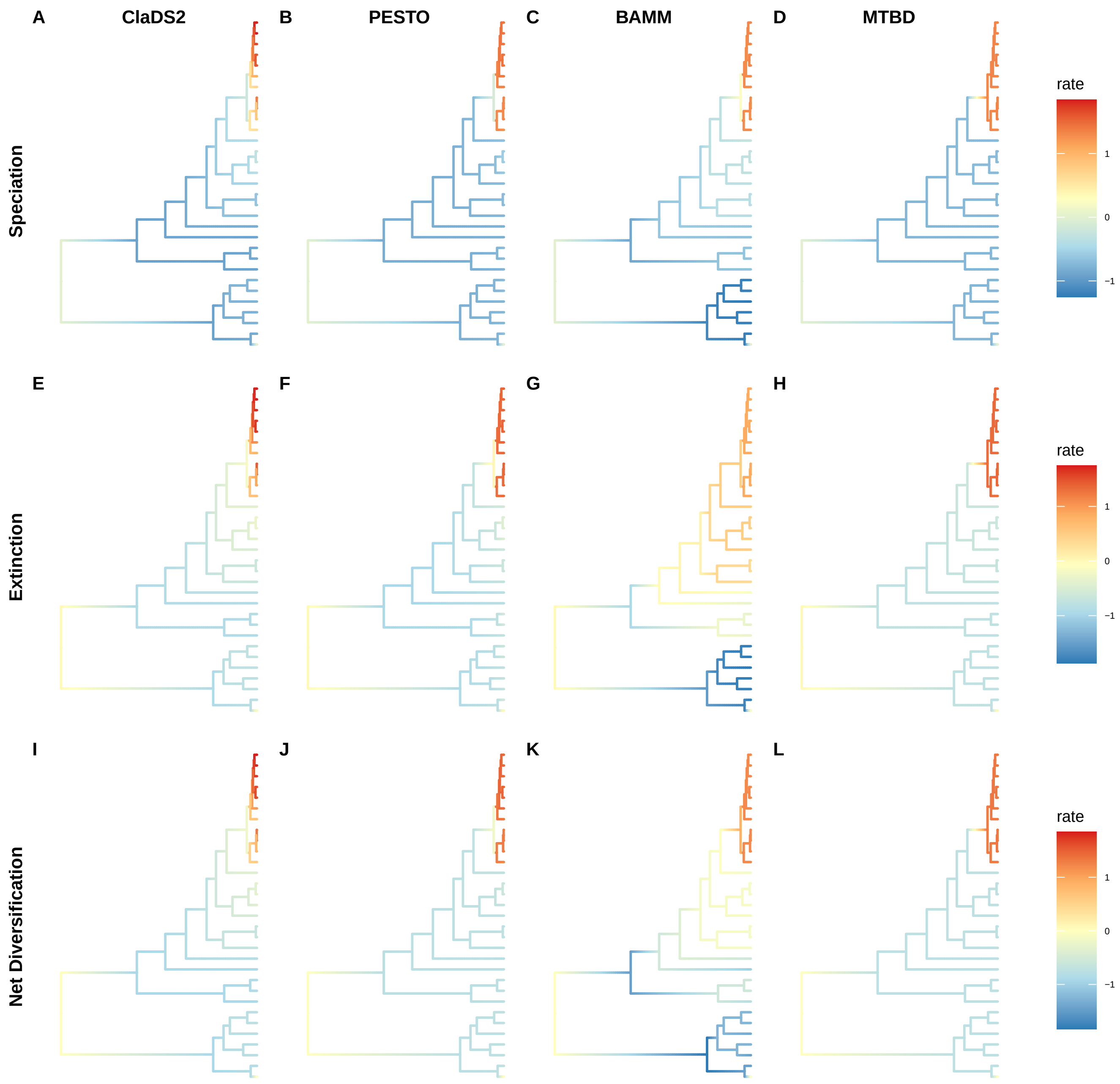
A
ClaDS2
B
PESTO
C
BAMM
D
MTBD
Speciation
rate
1
0
−1
E
F
G
H
Extinction
rate
1
0
−1
I
J
K
L
Net Diversification
rate
1
0
−1

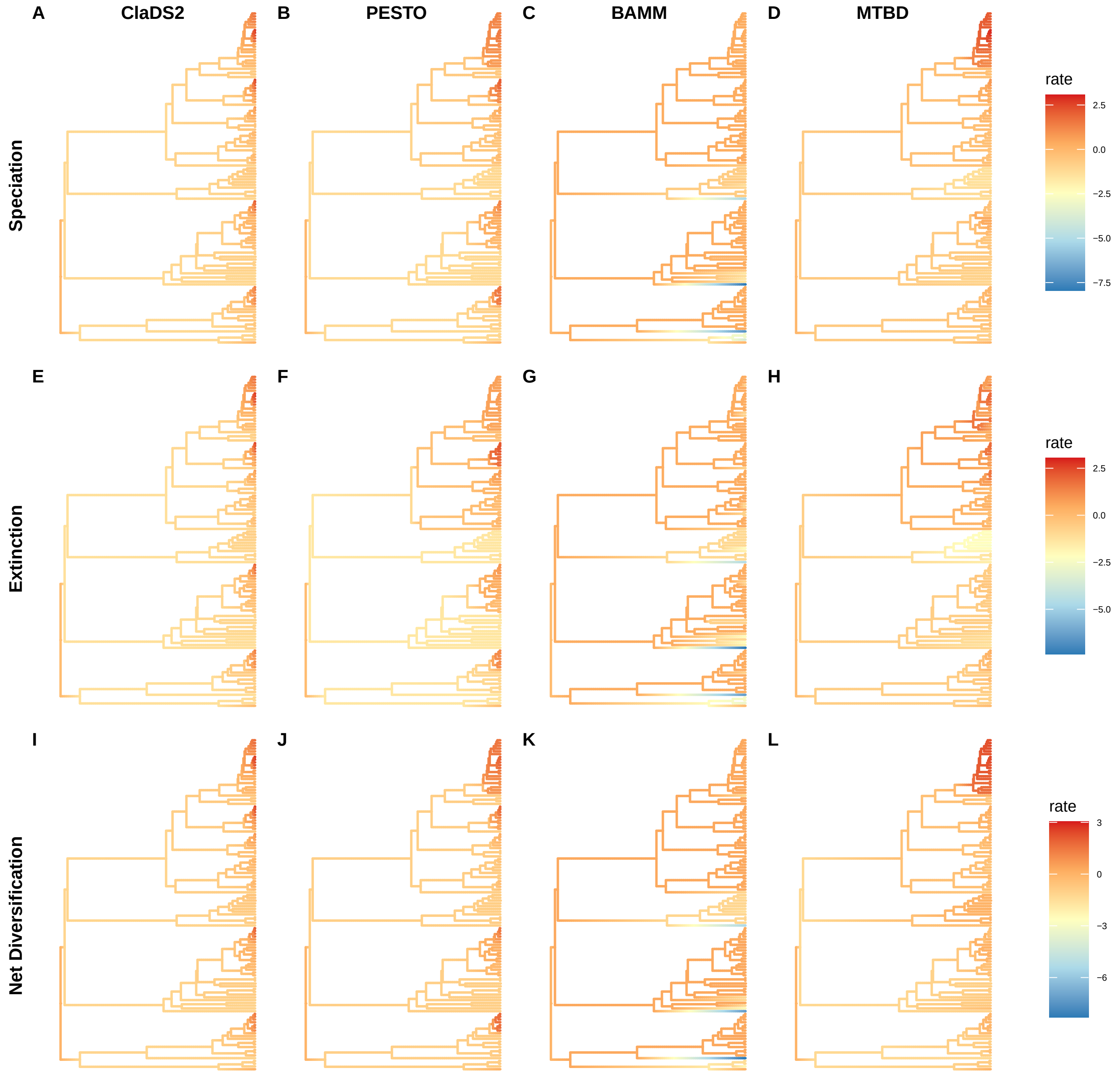
A
ClaDS2
B
PESTO
C
BAMM
D
MTBD
Speciation
rate
2.5
0.0
−2.5
−5.0
−7.5
E
F
G
H
Extinction
rate
2.5
0.0
−2.5
−5.0
I
J
K
L
Net Diversification
rate
3
0
−3
−6

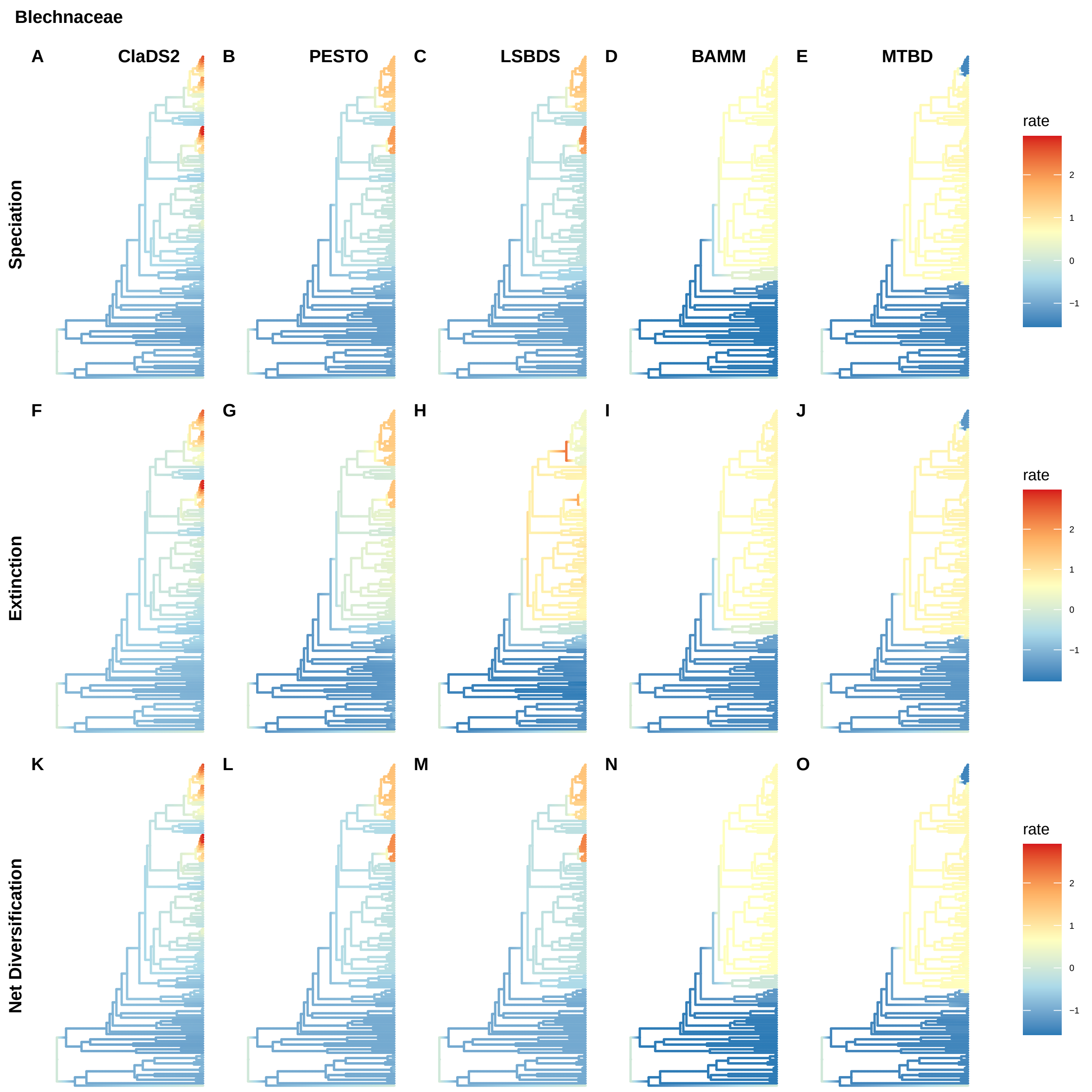
Blechnaceae
A
ClaDS2
B
PESTO
C
LSBDS
D
BAMM
E
MTBD
Speciation
rate
2
1
0
−1
F
G
H
I
J
Extinction
rate
2
1
0
−1
K
L
M
N
O
Net Diversification
rate
2
1
0
−1

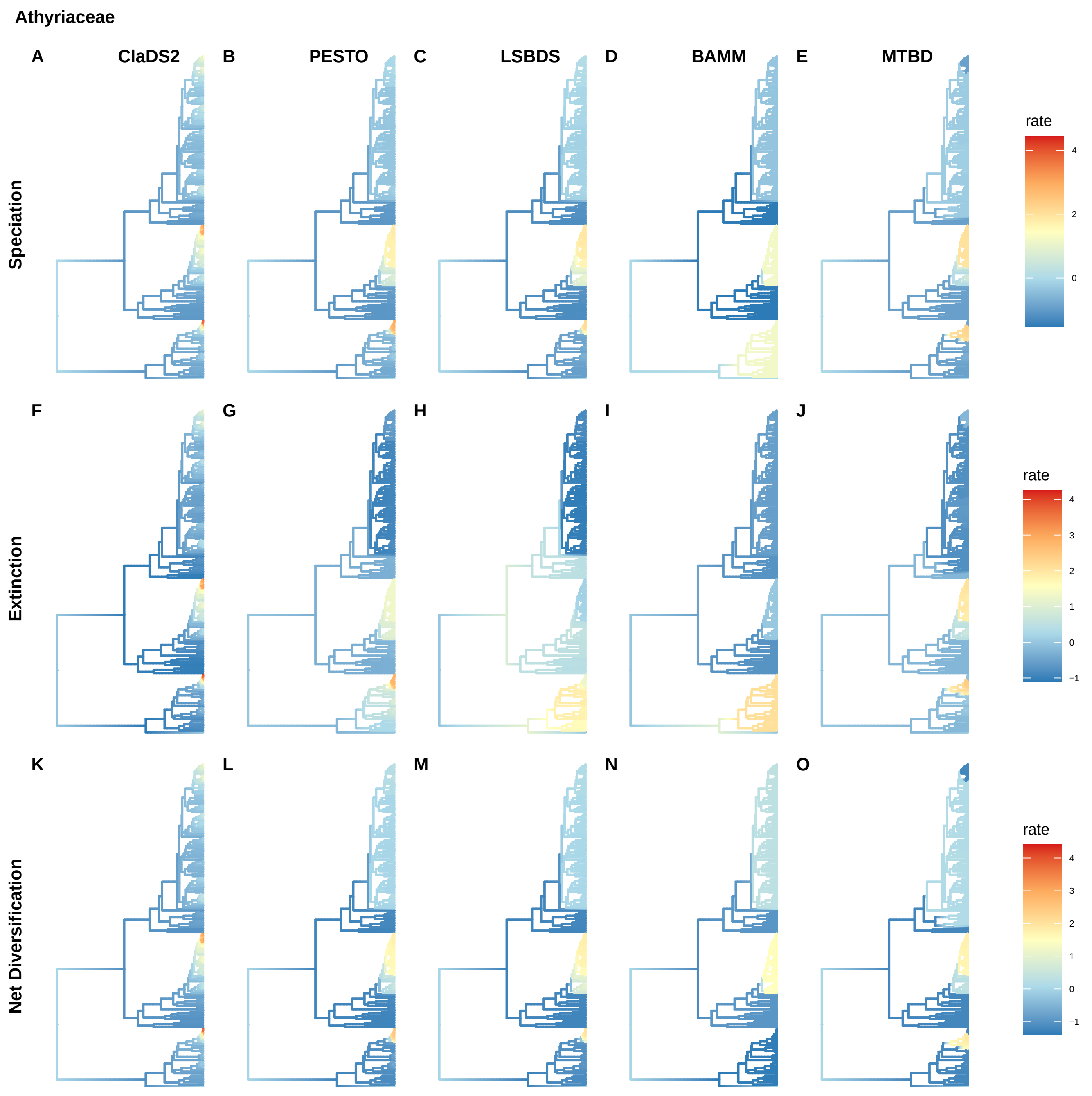

Athyriaceae
A
ClaDS2
B
PESTO
C
LSBDS
D
BAMM
E
MTBD
Speciation
rate
4
2
0
F
G
H
I
J
Extinction
rate
4
3
2
1
0
−1
K
L
M
N
O
Net Diversification
rate
4
3
2
1
0
−1

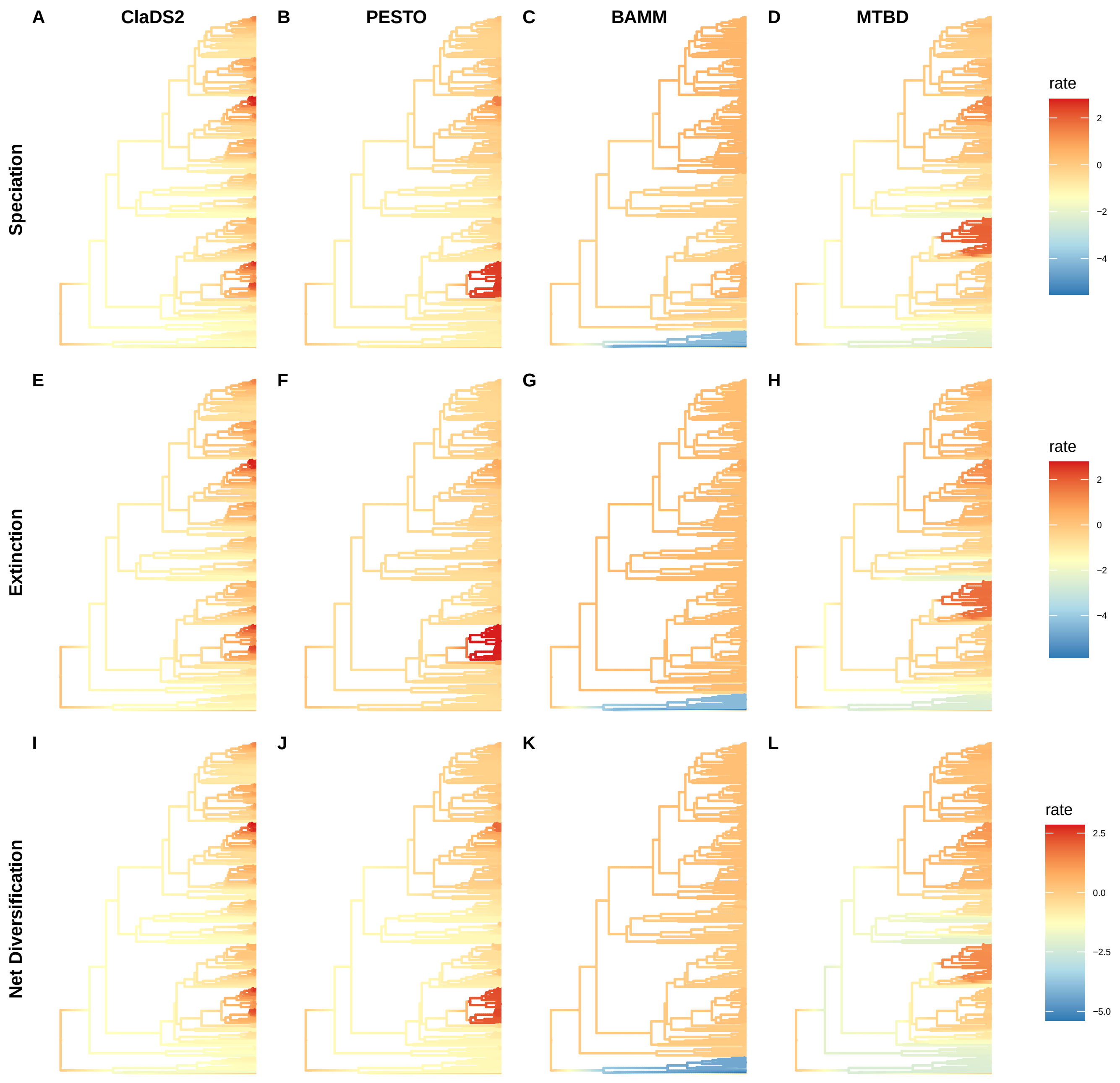

A
ClaDS2
B
PESTO
C
BAMM
D
MTBD
Speciation
rate
2
0
−2
−4
E
F
G
H
Extinction
rate
2
0
−2
−4
I
J
K
L
Net Diversification
rate
2.5
0.0
−2.5
−5.0

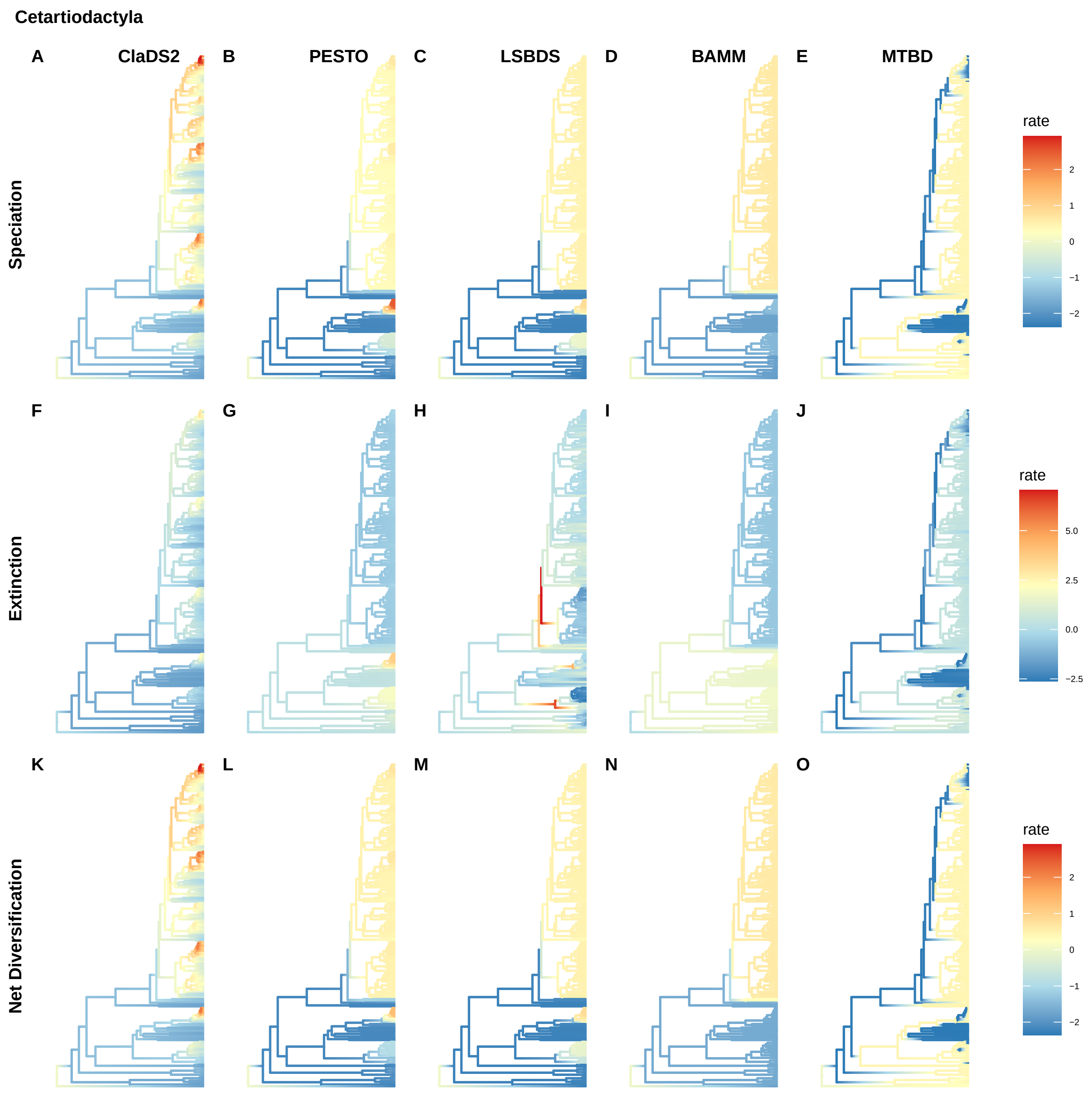
Cetartiodactyla
A
ClaDS2
B
PESTO
C
LSBDS
D
BAMM
E
MTBD
Speciation
rate
2
1
0
−1
−2
F
G
H
I
J
Extinction
rate
5.0
2.5
0.0
−2.5
K
L
M
N
O
Net Diversification
rate
2
1
0
−1
−2
